# Stochastic pausing at latent HIV-1 promoters generates transcriptional bursting

**DOI:** 10.1101/2020.08.25.265413

**Authors:** Katiana Tantale, Encarnation Garcia-Oliver, Adèle L’Hostis, Yueyuxio Yang, Marie-Cécile Robert, Thierry Gostan, Meenakshi Basu, Alja Kozulic-Pirher, Jean-Christophe Andrau, Florian Muller, Eugenia Basyuk, Ovidiu Radulescu, Edouard Bertrand

**Author notes:** Equal contribution (Katiana Tantale, Encarnation Garcia-Oliver).

## Abstract

Promoter-proximal polymerase pausing is a key process regulating gene expression. In latent HIV-1 cells, it prevents viral transcription and is essential for latency maintenance, while in acutely infected cells the viral factor Tat releases paused polymerase to induce viral expression. Pausing is fundamental for HIV-1, but how it contributes to bursting and stochastic viral reactivation is unclear. Here, we performed single molecule imaging of HIV-1 transcription, and we developed a quantitative analysis method that manages multiple time scales from seconds to days, and that rapidly fits many models of promoter dynamics. We found that RNA polymerases enter a long-lived pause at latent HIV-1 promoters (>20 minutes), thereby effectively limiting viral transcription. Surprisingly and in contrast to current models, pausing appears stochastic and not obligatory, with only a small fraction of the polymerases undergoing long-lived pausing in absence of Tat. One consequence of stochastic pausing is that HIV-1 transcription occurs in bursts in latent cells, thereby facilitating latency exit and providing a rationale for the stochasticity of viral rebounds.

## Introduction

Transcription initiation is a complex process that comprises chromatin opening, assembly of a pre-initiation complex (PIC), polymerase recruitment and finally its maturation into an elongation-competent form (see ^1^ for review). In Drosophila and mammals, this last step is highly regulated and appears to be a key point in the control of gene expression (^2^ for review). RNA polymerase II (RNAPII) is recruited by the PIC in a hypo-phosphorylated form and is then loaded on a short stretch of single stranded DNA, which is melted by TFIIH. The initiating polymerase starts elongating about a dozen of nucleotides and must undergo a number of modifications before leaving the promoter and entering productive elongation ^3^. First, the TFIIH-associated CDK7 kinase phosphorylates the Serine 5 of the heptad repeats of the C-terminal domain (CTD) of RNAPII, thereby disrupting interaction with Mediator and facilitating promoter escape (^4,5^ for reviews). The S5 phosphorylated CTD also recruits the RNA capping enzymes that access the RNA 5’-end when it emerges from the polymerase ^6^. The polymerase then transcribes an additional 10-80 nucleotides and typically enters a paused state. Two factors appear particularly important to trigger pausing, in relation with TFIID ^7^: DSIF (DRB sensitivity-inducing factor), which is composed of SPT4 and SPT5, and NELF (negative elongation factor), a four subunit complex that also interacts with the cap via the cap-binding complex ^8^ (CBC). A recent structure of the pausing complex indicates that the RNA-DNA hybrid adopts a tilted conformation within the polymerase that prevents further nucleotide addition ^9^. This structure is stabilized by NELF and DSIF, which also prevent binding of TFIIS, a factor that can trigger cleavage of the RNA at the active site to restart backtracked polymerases ^10^. Release from the paused state requires the positive transcription elongation factor b (P-TEFb), which is composed of Cyclin T1 or T2 associated with the kinase CDK9 ^11^, sometimes in association with the super-elongation complex ^12,13^ (SEC). P-TEFb is activated by CDK7 ^4,5,14^ and it phosphorylates a number of components of the pausing complex to enable formation of an elongation-competent polymerase ^9,15,16^. Phosphorylation of NELF triggers its dissociation from the polymerase, and this frees a binding site for PAF, an elongation factor that is required for transcription through chromatin. P-TEFb also phosphorylates the RNA polymerase CTD on its Serine 2, as well as the linker between the polymerase core and the CTD, creating a binding site for the elongation factor SPT6 ^9^. DSIF functions both as a repressor and activator of elongation, and it is also phosphorylated by P-TEFb (^17^ and ref therein). The structures of the paused and active elongation complex show that DSIF adopts different conformations in the two complexes. In particular, phosphorylated DSIF frees the nascent RNA and allows the polymerase to clamp around the DNA, promoting elongation while preventing release of the polymerase from DNA. Overall, P-TEFb mediated phosphorylation thus disrupts the pausing complex and triggers formation of an active elongation complex comprising the polymerase associated to DSIF, SPT6, and PAF.

While pausing is thought to be a key regulatory point for many cellular promoters in mammals and Drosophila, it is often revealed by a peak of RNAPII near the promoter that can in fact correspond to different molecular processes ^18^, such as slow elongation, polymerase arrest, or defective processivity (i.e. abortive initiation). Recent efforts have been made to clarify these mechanisms by measuring pausing duration. These studies indicated that pausing time vary from less than a minute up to an hour in Drosophila, depending on the promoter ^19-23^. This revealed a surprising variability in pausing kinetics, with widely different regulatory potential.

Another major finding of the last 15 years is that transcription is a discontinuous process in vivo (^24^ see ^25,26^ for reviews), with “active” genes going through active and inactive periods in a stochastic manner, a phenomenon also called transcriptional noise or gene bursting. In particular, recent evidences suggest that for many genes, expression levels are dynamically encoded in the time domain by controlling the periods during which a gene is active, rather than by regulating the initiation rate ^27-29^. Major efforts have been made to decipher the causes of gene bursting and in particular the molecular status of the postulated ON and OFF states. Indeed, the transitions between these states are kinetically rate limiting and therefore represent key regulatory checkpoints. However, despite these efforts and the importance of pausing in regulating gene expression, how pausing affects gene bursting remains not understood.

An important implication of gene bursting is that it creates cell-to-cell heterogeneity and this has multiple consequences on the phenotypes of single cells or multicellular organisms. For instance, stochasticity in the expression of Heat-Shock genes in yeasts is thought to help a fraction of the yeast population survive sublethal stresses ^30^, while in C. Elegans, mutations in a small gene regulatory network create a high expression variability, ultimately leading to variable phenotypic penetrance of the mutation ^31^. In the case of HIV-1, transcriptional noise is thought to play a crucial role in the control of latency. Indeed, HIV-1 infection generates latent cells that can persist in the body for decades and can re-establish viral propagation when antiviral treatments are interrupted. Previous studies from the Siliciano and Weinberger labs have shown that latency exit is stochastic and linked to random fluctuations of viral transcription ^32-34^. How the viral promoter creates bursts of gene expression in latent cells is not understood, but nevertheless fundamental as it is triggering latency exit. A better knowledge of mechanistic and quantitative aspects of the reactivation dynamics is indeed essential for the development of new strategies in combinatorial anti-retroviral therapies such as “shock and kill” and “block and lock”.

The ability of the virus to alternate between acute and latent forms lies in a positive transcriptional feedback loop established by the viral protein Tat (^32^, see ^35,36^ for reviews). In latent cells, Tat levels are very low and viral transcription remains silent or also low. In acutely infected cells, Tat levels are elevated, strongly inducing viral transcription. It is well established that in absence of Tat or when Tat levels are low, P-TEFb is limiting for viral transcription and the polymerases that initiate transcription enter a paused state after transcribing about 60 nucleotides and fail to enter productive elongation (reviewed in ^35,36^; Figure 1A, left). Tat alleviates this block by binding both P-TEFb and the TAR stem-loop at the 5’-end of nascent HIV-1 RNAs, leading to the formation of a ternary complex that promotes elongation by recruiting P-TEFb and its associated super-elongation complex to paused polymerases ^11-13^ (Figure 1A, right). The HIV-1 promoter is thus strictly regulated at the level of pausing and P-TEFb recruitment, and these steps are controlled by Tat, which overall can activate viral transcription by more than 100 fold. These properties make HIV-1 an attractive model to decipher how pausing affects gene bursting, with direct relevance for HIV-1 latency and pathogenesis ^37,38^.

**Figure 1.**
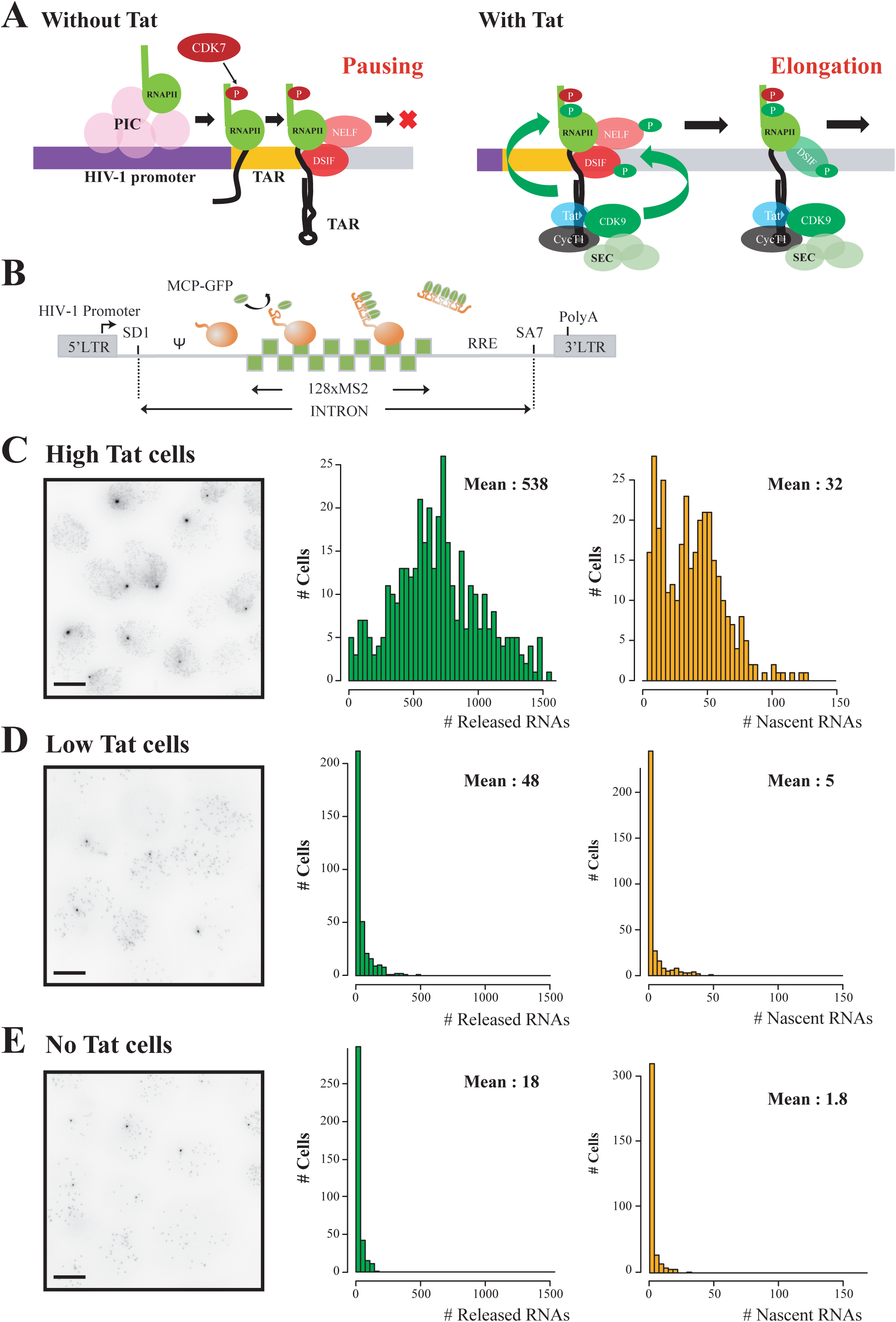
Single cell characterization of HIV-1 gene expression, with and without Tat. **A**-Schematic of HIV-1 transcriptional regulation. Left: in absence of Tat, pTEFb is not recruited and polymerases binds NELF and DSIF and pause near the promoter. Right: in presence of Tat, pTEFb, composed of Cyclin T1 and Cdk9 associated to the super elongation complex, is recruited to the nascent TAR RNA. Cdk9 phosphorylates NELF, DSIF and RNA polymerase II, thereby triggering pausing exit and processive elongation. **B**-Schematic of the HIV-1 reporter construct. SD1: major HIV-1 splice site donor; SA7: last HIV-1 splice site acceptor; ψ: packaging signal; RRE: Rev-responsive element; LTR: long terminal repeat. **C**-Expression of the 128xMS2 HIV-1 tagged reporter in cells expressing high levels of Tat. <u>Left panel:</u> microscopy images of High Tat HeLa cells where the unspliced HIV-1 pre-mRNA is detected by smFISH with probes against the 128xMS2 tag. Cells bear a single copy of the reporter gene integrated with the Flp-in system. The bright spots in the nuclei correspond to nascent RNA at their transcription sites, while the dimmer spots correspond to single pre-mRNA molecules. Scale bar : 10 μm. <u>Middle panel:</u> distribution of the number of released HIV-1 pre-mRNAs per cell, in High Tat cells. Experimental RNA distribution are from smFISH data. X-axis: number of HIV-1 pre-mRNA molecules per cell; y-axis: number of cells; inset: mean number of HIV-1 pre-mRNAs per cell. <u>Right panel:</u> distribution of the number of nascent HIV-1 pre-mRNAs per transcription site, in High Tat cells. Experimental RNA distribution are from smFISH data. X-axis: number of nascent HIV-1 pre-mRNA molecules per transcription site; y-axis: number of transcription sites; inset: mean number of nascent HIV-1 pre-mRNAs per cell. **D**-Expression of the 128xMS2 HIV-1 tagged reporter in cells expressing low levels of Tat. Legend as in C, except that experiments are from Low Tat cells. **E**-Expression of the 128xMS2 HIV-1 tagged reporter in cells not expressing Tat. Legend as in C, except that experiments are from No Tat cells. Image contrast adjustment is identical for panels C, D and E.

Here, we imaged HIV-1 transcription in live cells at the level of single polymerases. We characterized the effect of pausing on gene bursting by modulating the levels of Tat, which controls pausing at the HIV-1 promoter. We provide the first fully quantitative description of the stochastic activity of the HIV-1 promoter in basal and induced conditions, on timescales ranging from second to tens of hour. Surprisingly, we found that promoter-proximal pausing is a stochastic event that generates large viral bursts even in cells that do not express Tat. In HIV-1 latent cells with a functional but inactive Tat loop, stochastic pausing may be a key phenomenon that determines latency exit.

## Results

### Single molecule imaging of HIV-1 transcription with different levels of the pause release factor Tat

We previously developed an improved MS2 tagging system based on a 128xMS2 tag and designed for long term tracking of single RNAs ^28^. To image viral transcription, we inserted this tag in the intron of an HIV-1 vector that had all the viral sequences responsible for transcription and RNA processing (Figure 1A-B). The corresponding pre-mRNA splices entirely post-transcriptionally, enabling imaging of transcription independently of splicing ^28,39^. The high number of MS2 stem-loops present in this reporter allows for a 5-fold increase in signal as compared to our original 24xMS2 repeat ^40^. This enables the use of a low illumination power to limit photo-bleaching, allowing to capture five times more images while still detecting single RNA molecules. By using the 128xMS2 tag and monitoring the brightness of the transcription site over time, it is possible to measure promoter activity with a temporal resolution in the second range and for hours.

It has been demonstrated by numerous studies that the HIV-1 promoter is regulated at the level of promoter proximal pausing (see ^35,36^ for reviews). Indeed, latent cells do not express a significant amount of Tat and in this case, polymerases that start transcribing are blocked ∼60 nucleotides downstream the transcription start site and do not enter productive elongation. This block is relieved by Tat, which directly alleviates pausing by recruiting P-TEFb to the nascent viral RNAs and allowing polymerases to elongate throughout the entire viral genome. To characterize how pausing affects HIV-1 transcription, we therefore created isogenic cell lines expressing different levels of Tat. These lines all contained the 128xMS2 reporter integrated at the same chromosomal location. We previously generated a HeLa cell line that expressed in *trans* a saturating amount of Tat (*High Tat* cells). In these cells, transcription was high and a further increase in the amount of Tat did not lead to more viral transcription ^28^. We then created two new reporter cell lines with low levels of Tat to mimic the situation of latent cells where Tat is not expressed or only at very low levels ^35,36^. The first cell line expresses Tat from the second cistron of a bicistronic vector (referred to as *Low Tat* cells), and Tat was not detected by Western blot although it promoted HIV-1 transcription by 2.7 fold (Figure 1C-E and Figure S1A). The second cell line entirely lacked Tat (referred to as *No Tat*). We first determined the expression levels of the HIV-1 reporter by performing smFISH experiments with probes binding the 128xMS2 repeat. We found that expression of the HIV-1 reporter depended on Tat as expected (Figure 1C-E), as the number of pre-mRNA molecules present in the nucleoplasm dropped from ∼500 copies per nucleus in *High Tat* cells, to ∼50 and ∼20 in *Low Tat* and *No Tat* cells, respectively. This was mirrored by a similar decrease in the level of the nascent RNAs present at the transcription sites, with a mean of 32 copies for the *High Tat* cells, and only 5 and 1.8 for the *Low Tat* and *No Tat* cells, respectively (Figure 1C-E).

Next, we aimed at confirming that pausing was limiting viral transcription in *No Tat* cells. To this end, we overexpressed the two subunit of P-TEFb, Cdk9 and Cyclin T1, by transient transfection. We observed that this increased viral transcription as previously reported in other cellular systems (Figure S1B; ^41^). Then, we fused CDK9 to a fluorescent catalytically inactive Cas9 variant (dCas9-tagBFP), and we transfected the resulting construct in *No Tat* cells together with vectors expressing three Cas9 guide RNAs targeting the HIV-1 promoter. By performing smFISH with probes against the 128xMS2 repeat, we found that expressing dCas9-CDK9-tagBFP alone increased HIV-1 RNA levels by 4 fold, while further targeting it to the HIV-1 promoter with three guide RNAs led to a 10-fold increase in expression (Figure S1C). Moreover, the basal HIV-1 transcriptional activity in *No Tat* cells was blocked when P-TEFb was inactivated with KM05283, a drug that specifically inhibits CDK9 kinase activity (Figure S2A). This indicated that P-TEFb was both required for basal transcription and also limiting viral expression, providing functional indications that pausing was limiting in *No Tat* cells. Next, we tested whether the basal viral transcription observed without Tat was due to sporadic activation of the NF-κB pathway, as it is a well-known activator of the HIV-1 promoter that can recruit P-TEFb ^42,43^. We treated cells with BAY11-7082, a drug that inhibits the IKK kinase and traps NF-κB subunits in the cytoplasm. No difference in HIV-1 expression was seen after 16h of treatment, indicating that the basal viral transcription was independent of NF-κB (Figure S2B-C). Taken together, these data indicate that in our cellular system, the basal HIV-1 transcription occurring in absence of Tat is P-TEFb dependent, and that the recruitment of this factor is a key step limiting viral transcription, as expected from a large body of previous studies.

### The absence of Tat does not affect the formation of polymerase convoys but creates long inactive periods

When Tat is in excess, HIV-1 transcription occurs in the form of polymerase convoys, i. e. sets of closely spaced polymerases that transcribe the gene together (see schematic in Figure 2D; ^28^). In average, the Tat-activated HIV-1 promoter produces convoys of 19 polymerases, each polymerase spaced every ∼4 seconds, with a convoy being fired every ∼2 minutes. In order to characterize how a limiting amount of Tat affects the viral transcriptional output, we performed live-cell imaging using MCP-GFP and monitored the brightness of transcription sites over time. The single molecules of unspliced pre-mRNA present in the nucleoplasm were used to calibrate the signal at the transcription site, which could then be expressed as an absolute number of RNA molecules (Figure 2). We previously showed that the Tat-activated HIV-1 promoter fluctuates on time scales ranging from minutes to hours, and we therefore recorded two types of movies to cover the entire temporal range of transcriptional fluctuations ^28^. ‘Short movies’ capture one image stack every 3 seconds for 15 to 20 minutes, and they allow a detailed characterization of rapid transcriptional fluctuations such as polymerase convoys. ‘Long movies’ last for 8 hours with a rate of one image stack every three minutes, and they allow to measure the frequency and duration of long inactive periods. Note that since a nascent RNA resides 2.8 minutes at the transcription site ^28^, this frame rate ensures that all the initiation events are detected in the long movies.

**Figure 2.**
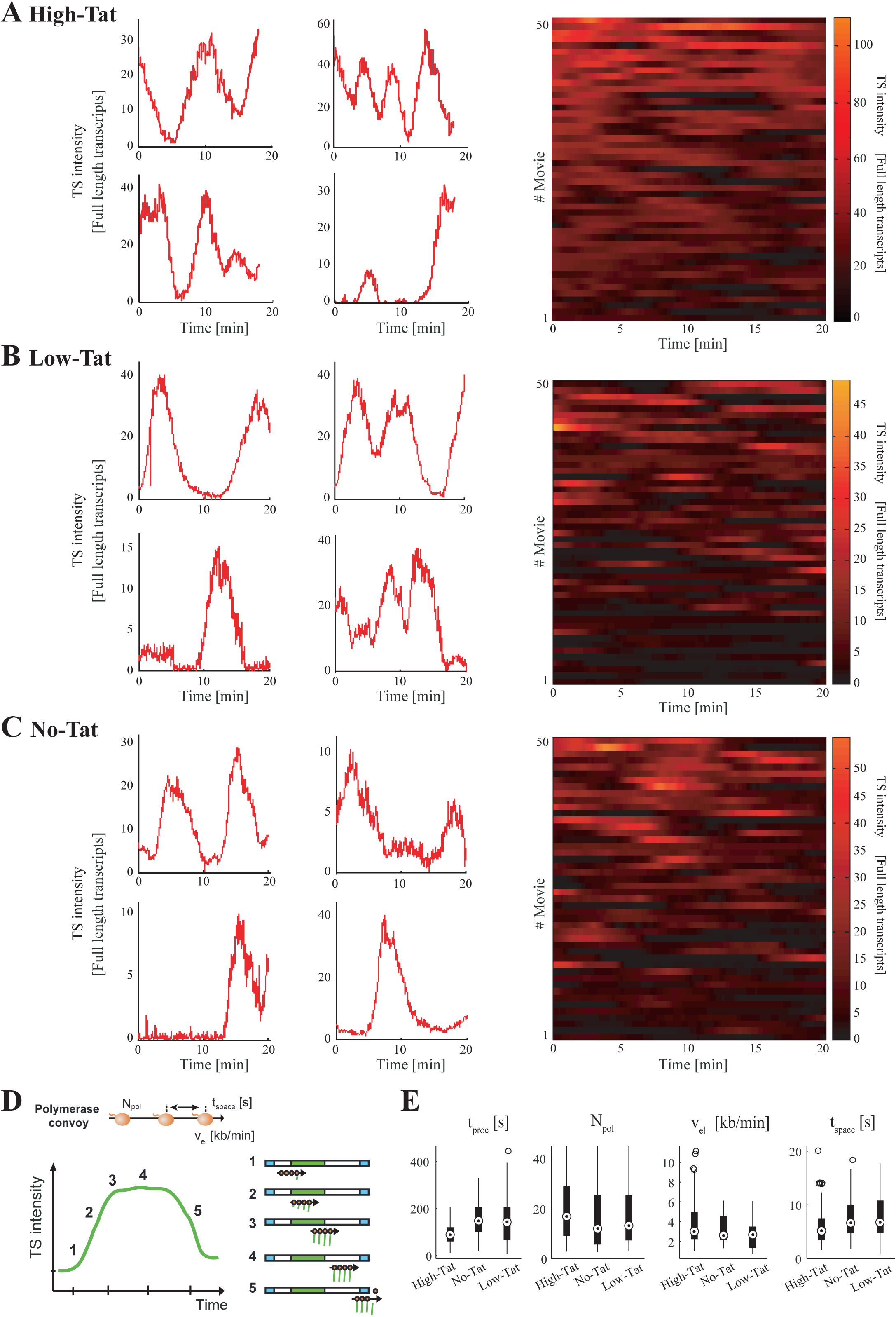
Fluctuation of HIV-1 transcription over short time periods, with and without Tat. **A-C** Fluctuations of HIV-1 transcription over 15-20 minute periods, with one image stack recorded every 3 seconds. Left: each graph is a single transcription site; the x-axis represents the time (in minutes) and y-axis represents the intensity of transcription sites, expressed in equivalent numbers of full-length pre-mRNA molecules. Right: each line is a cell and the transcription site intensity is color-coded (scale on the right). A: High-Tat cells; B: Low-Tat cells; C: No-Tat cells. **D**-Schematic of a polymerase convoy. Top: a polymerase convoy, with polymerases in orange and the gene represented as a black horizontal arrow. *N*_*pol*_ : number of polymerases; *t*_*space*_ : spacing between successive RNA polymerases (in seconds); *v*_*el*_ : elongation rate. Bottom: schematics describing the different phases of a transcription cycle (left) and the position of the polymerase convoy on the MS2 tagged gene (right; the green box is theMS2 tag). **E**-Box-plots representing the parameters values of the best-fit models, measured for a set of isolated transcription cycles in each cell line. *t*_*proc*_ is the 3’-end RNA processing time; *Npol* is the number of polymerases in the convoy; *V*_*el*_ is the elongation rate (in kb/min); *t*_*space*_ is the spacing between successive polymerase (in seconds). The bottom line displays the first quartile, the box corresponds to the second and third quartile, the top line to the last quartile, and the double circle is the median. Small circles are outliers (1.5 times the inter-quartile range above or below the upper and lower quartile, respectively).

In the short movies, we observed transient increases in the brightness of transcription sites for all three cell lines: *High Tat, Low Tat* and *No Tat* (Figure 2A-C). They were in the minute range and quantification of the signals indicated that they corresponded to the synthesis of multiple RNA molecules (Figure 2A-C). Thus, viral transcription occurred in large bursts even in absence of Tat, resulting in the formation of polymerase convoys. To better characterize these rapid fluctuations, we focused on transcription cycles in which an inactive transcription site transiently turned on, and we fitted these data with a model of polymerase convoys (^28^; see schematic in Figure 2D). Surprisingly, the convoys formed in the *Low Tat* and *No Tat* cells were roughly similar to those formed when Tat was saturating (19 polymerases initiating every 4 seconds in *High Tat* cells, compared to 14 polymerases every 6 s in *Low Tat* cells and 12 polymerases every 8 s in absence of Tat; Figure 2E). This result was unexpected because decreasing Tat levels should increase pausing, which should increase in lag time between successive polymerases, possibly until convoys are no longer formed. It is also interesting to note that the differences observed at this rapid time-scale were small and could not account for the 30 fold difference in expression induced by Tat (Figure 1C-E).

Next, we analyzed fluctuations on slow time scales using long movies. The HIV-1 promoter was almost always active in cells expressing an excess of Tat (Figure 3A-B, left panels). In contrast, *No Tat* and *Low Tat* cells displayed long inactive periods that lasted for hours (Figure 3A-B, middle and right panels). In addition, active periods were brief and rare, yet yielded initiation of multiple polymerases in the form of convoys as for *High Tat* cells (see Figure 3A). The activity of the HIV-1 promoter in absence of Tat thus occurs mainly in the form of sparse, yet large bursts, with long inactive period explaining most of the difference in promoter activity with and without Tat.

**Figure 3.**
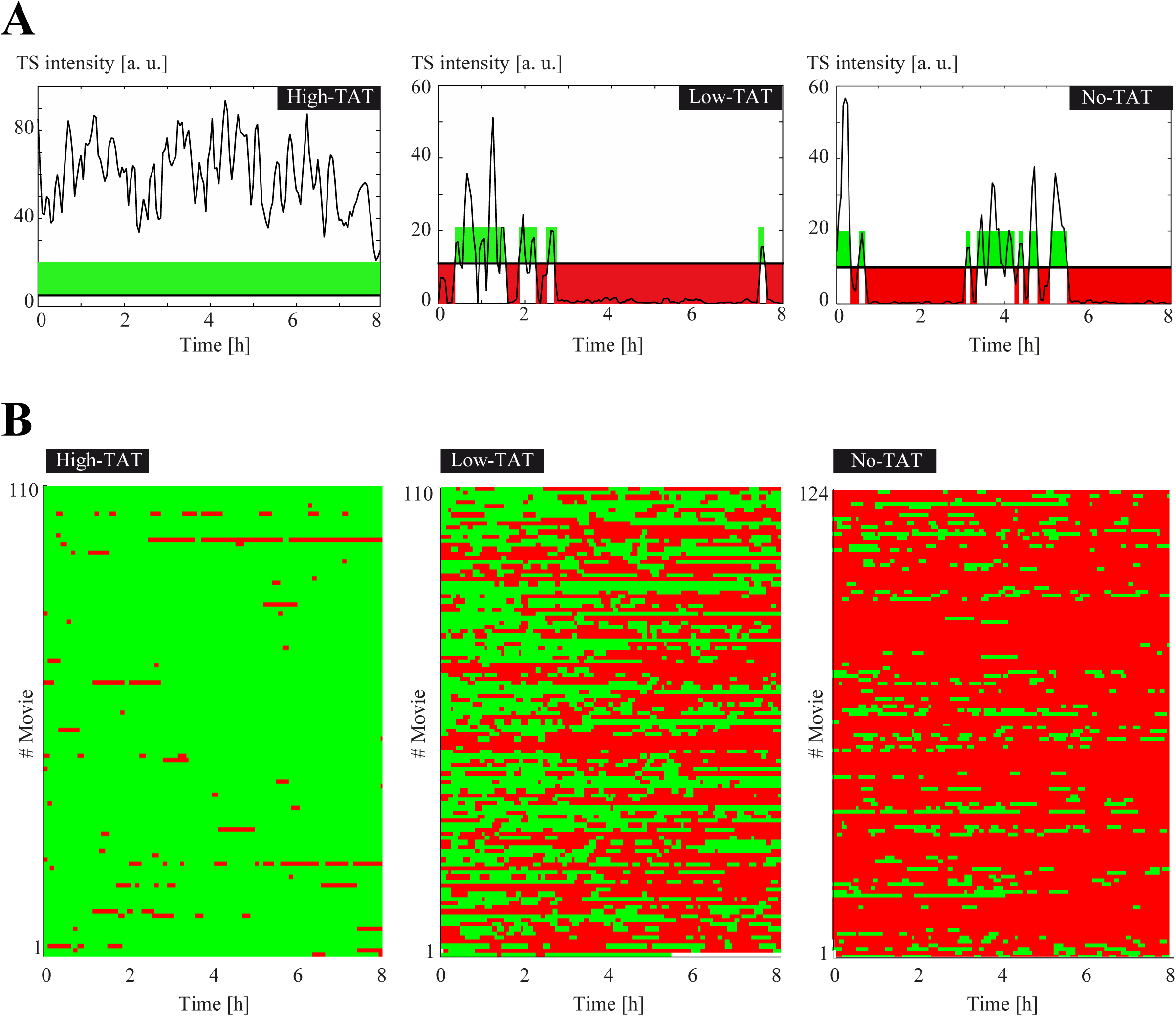
Fluctuation of HIV-1 transcription over long time periods, with and without Tat. **A**-Fluctuations of HIV-1 transcription over 8 hours, with one image stack recorded every 3 minutes. The x-axis represents the time (in hours) and y-axis represents the intensity of transcription sites, expressed in arbitrary units. Periods of HIV-1 promoter activity are colored in green, and periods of inactivity in red. **B**-Active and inactive periods of the HIV-1 promoter, for the indicated cell lines. Each line is a cell and the activity of the HIV-1 promoter is color-coded (green: active; red: inactive), using the threshold shown in panel A. x-axis: time in hours.

### Development of a novel analysis pipeline to characterize the fluctuations of promoter activity on multiple timescales

The fluctuations of promoter activity arise from stochastic transitions between active and inactive promoter states (^25,26^; Figure 4A). These transitions correspond to steps that are kinetically rate-limiting, and the characterization of these promoter states can thus yield important information on how promoters function and are regulated. To better understand how pausing and Tat control the activity of the HIV-1 promoter, we turned to machine learning and modeling approaches with the aim of elucidating how the promoter switches between active and inactive states. The analysis of the fluctuations of transcription sites brightness can be done by auto-correlation strategies ^44,45^. This gives a direct measurement of the dwell time of the nascent RNAs and allows to estimate the elongation and 3’-end processing rates. However, there is currently no theoretical framework that can easily extend autocorrelation methods to models containing multiple promoter states besides a simple ON/OFF switch. In addition, correlation approaches are difficult to use when fluctuations are slow and approach the recording time of the movies. Other analysis strategies hypothesize a theoretical transition model and infer parameters using Bayesian or maximum likelihood approaches ^29,46-48^. These strategies rarely compare several models and do not directly characterize features such as polymerase convoys. To circumvent these difficulties, we turned to the direct analysis of polymerase waiting times, i.e. the lag time between two successive initiation events. Indeed, transcription can be modelled as a continuous time Markov chain in which a promoter stochastically switches between various non-productive states until it reaches an active state where it can initiate transcription (Figure 4A). In this case, waiting times between successive initiation events are interesting to consider because their distribution directly relates to transition rates of the Markov chain (see Supplemental Text). Moreover, we obtained for many different models the closed-form equations expressing the distribution of waiting times as a function of the model parameters (for full solutions to this direct problem, see Methods and Supplemental Text), as well as closed-form equations allowing to compute the model parameters directly from the distribution of waiting times, the so-called inverse problem (for full solutions, see Methods and Supplemental Text). In particular, if we consider a class of models containing several consecutive OFF states and one ON state that can initiate transcription (Figure 4A), the survival function of polymerase waiting times, which is one minus their cumulative distribution, is the sum of several exponentials with the number of exponentials corresponding to the number of promoter states (Figure 4A; see Supplemental Text). Thus, by fitting the survival function with various sums of exponentials, one can determine the number of states in the promoter model. In addition, the rates of promoter switching can be directly calculated from the coefficients of the fitted sum of exponentials (see Methods and Supplemental Text, inverse problem). Hence, if the distribution of waiting times can be extracted from the experimental data, it is straightforward to determine both the number of promoter states, as well as the rates of switching between these states.

**Figure 4.**
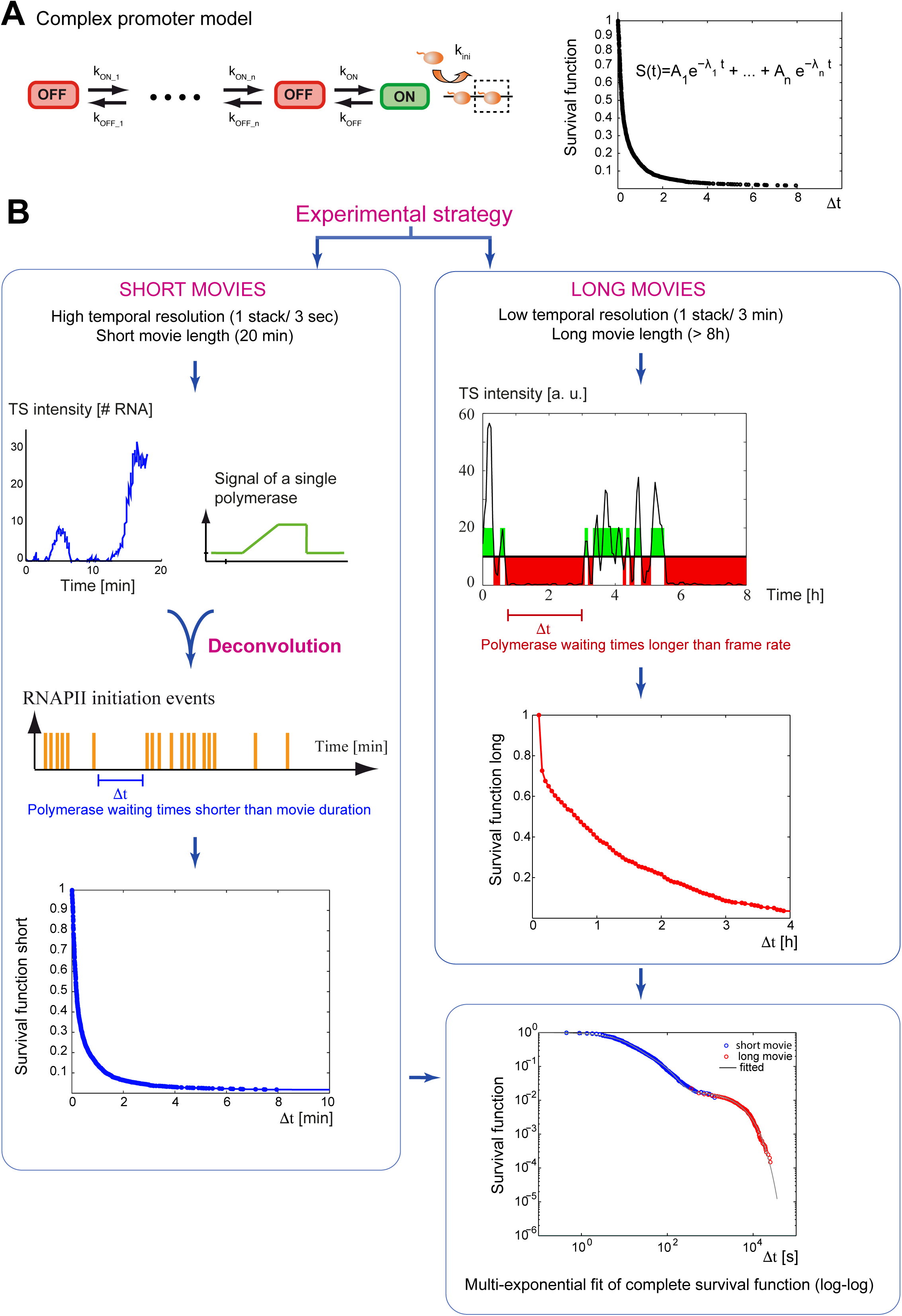
Analysis and modeling strategy for the live cell transcriptional data. A-Determination of models for transcription initiation. Left: example of a complex promoter models describing the different steps leading to transcription initiation and their kinetic relationship. OFF: inactive promoter state; ON: active promoter state; orange ball: RNA polymerase. Right: the survival function (equal to one minus the cumulative function) describes the distribution of polymerase waiting times (delay between two successive initiation events). For linear models such as the one depicted on the left, the survival function can be fitted by a sum of exponentials, with the number of exponentials being equal to the number of promoter states. Branched models also lead to sums of exponentials (see text). **B**-Experimental and machine learning strategy to determine the survival function of polymerase waiting times. Left: signals of short movies made at high temporal resolution result from the convolution of the signal from a single polymerase and the sequence of temporal positions of initiation events. The sequence of initiation events can thus be reconstructed by a deconvolution numerical method, provided that the signal of a single polymerase is known. This allows to estimate the distribution of waiting times for waiting times shorter than the movie duration (i.e. a conditional distribution). Right: long movies made with a lower temporal resolution, in the order of the residency time of RNA polymerase on the gene (3 minutes), allow to estimate the distribution of polymerase waiting times for waiting times greater than the temporal resolution. The two conditional survival functions, short and long, can then be combined to reconstitute the complete survival function, with the constraint that waiting times of short movies, smaller than the frame rate of the long movie, must fill the active periods in the long movie. Finally, the complete survival function is fitted with a sum of exponentials to determine the number of promoter state, the kinetics of transitions between them, and the initiation rate. Multiple models can be easily fitted to the same survival function and the most appropriate one is selected based on parsimony, parametric indeterminacy and consistency with complementary experiments.

### Calculation of polymerase waiting times from short and long movies

We first reasoned that the inactive periods seen in the long movies correspond to long polymerase waiting times (Figure 3A-B). Since the frame rate is 3 minutes while a nascent RNA remains 2.8 minutes at the transcription site ^28^, these movies should detect all initiation events and thus to measure all the polymerase waiting times longer than 3 minutes. The short waiting times could be calculated from the short movies, which have a much higher frame rate (3 seconds). However, a difficulty is that the signal generated by a polymerase persists several minutes after it initiated, as the labelled nascent RNA leaves the transcription site only after it is transcribed to the end of the gene and 3’-end processed (see schematic in Figure 2D). Consequently, if the next polymerase appears before the nascent RNA disappears, the transcription site remains continuously fluorescent and it is not possible to directly calculate the polymerase waiting times. To circumvent this difficulty, we reasoned that the intensity of transcription sites over time is the result of the convolution of two functions: the signal produced by a single polymerase and the time sequence of firing events (see ^25^ and Figure 4B, left panels). The signal produced by a single polymerase depends on the polymerase elongation rate and the rate of 3’-end formation, which we determined previously for this HIV-1 reporter gene ^28,39^. If we assume that all polymerases behave identically, it is thus possible to calculate the temporal position of polymerase initiation events by finding the best sequence of these events that reproduces the experimental transcriptional fluctuations (Figure 4B). It should thus be possible to extract polymerase waiting times from the short movies, keeping in mind that the waiting times longer than the movie will be truncated and require a correction (see Supplemental Text).

Altogether, the long movies give access to waiting times longer than the frame rate (waiting times in the 3 min-10h range), and the short movies provide waiting times shorter than the movie length (in the 3s-20min range). The combination of these movies thus allows to reconstruct and estimate the distribution of polymerase waiting times over 4 logarithmic decades, i.e. 3s-10h (see Supplemental Text for the reconstruction procedure). This analysis pipeline has three advantages. First, by determining the number of exponentials required to fit the survival function, one can directly determine the number of promoter states in the model. Second, given that equations describing the distribution of waiting times can be obtained for many models, it is straightforward to fit these models and to estimate which model best fits the experimental data. Finally, this pipeline enables to combine data acquired at multiple time-scales, from seconds to ten hours, and therefore provides an ideal framework to quantify transcriptional dynamics in live cells.

### Validation of the analysis pipeline by simulations

To evaluate the precision and reliability of the analysis pipeline, we first tested the performance of the deconvolution algorithm on simulated datasets. The initiation times of several polymerases were simulated and the signal of an imaginary transcription site was calculated using experimentally measured elongation and 3’-end processing rates ^28,39^. We then added a realistic amount of noise and tested the ability of the deconvolution algorithm to reconstruct the proper initiation timing from the noisy signal (Figure 5A and Supplemental Text). The algorithm is composed of two parts: a genetic algorithm to obtain the rough position of initiation events, and by a local optimization to refine the position of initiation events. In both presence and absence of noise, the combination of the two steps allowed an accurate positioning of the initiation events.

**Figure 5.**
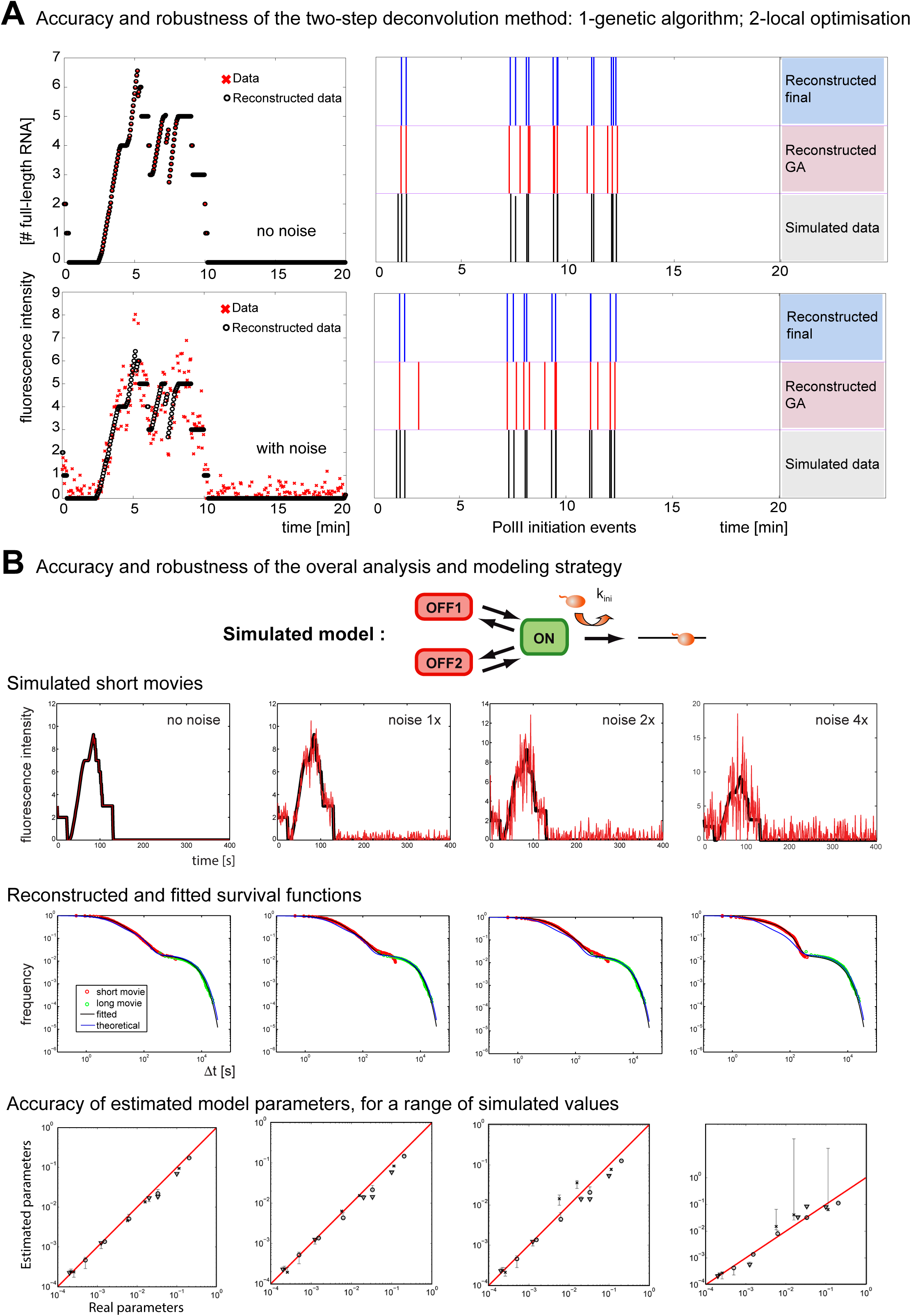
Accuracy and robustness of the analysis and modeling pipeline. **A**-Fidelity and robustness of the deconvolution method. Left panels: simulation of short movies for an artificial set of polymerase initiation events, with noise added (bottom), or without (top). x-axis is time in minutes; y-axis is the intensity of transcription sites (expressed in number of RNA molecules). Right panels: positions of the transcription initiation events (vertical bars), for the original artificial data (black; bottom lines), the reconstructed data from the simulated short movies after the genetic algorithm (GA, red, middle lines), or the final reconstruction after both the GA and the local optimization (blue; top lines). x-axis is time in minutes. **B**-Fidelity and robustness of the overall analysis pipeline. Top schematic: the linear three state promoter model used for Monte Carlo simulations. Top graphs: examples of artificial short movies (black lines), with various levels of noise added (red lines). Note that the noise level measured experimentally corresponds to the 1x condition. x-axis is time in seconds; y-axis is the intensity of transcription sites expressed in number of RNA molecules. Middle graphs: survival functions reconstructed from artificial short and long movies (red and green circles, respectively), and fitted to a sum of three exponentials (black line). The theoretical survival function obtained with the model parameters used for the simulation is shown for comparison (blue line). x-axis: time intervals between successive initiations events, in seconds and in log_10_ scale. y-axis: probability of Δt > x (log_10_ scale). **C**-Accuracy of determining the model parameters. Graphs plot the parameters used to generate the artificial data (x-axis), against the parameter measured by the deconvolution and fitting procedure (y-axis). Vertical bars: confidence intervals. Three parameter sets were used, corresponding to the values obtained with the experimental data from the High Tat cells (circles), Low Tat cells (crosses), and No Tat cells (triangles).

Next, we validated the entire analysis pipeline by simulating a three state branched promoter model with the Gillespie algorithm, using several realistic sets of parameters (i.e. corresponding to values obtained with our cell lines, see below). We computed the brightness of many statistically equivalent imaginary transcription sites as above, and added different amounts of noise (1x, 2x and 4x), with the 1x condition corresponding to the noise observed in our experimental data (Figure 5B, upper panels; see Supplemental Text). The intensities of the simulated transcription sites were then resampled to create artificial short and long movies, which were treated exactly as real data. Simulated short movies were deconvolved and the distribution of waiting times was computed separately for the short and long movies. These distributions were then combined to reconstruct the entire distribution of waiting times (Figure 5B, middle panels), which was fitted to a sum of three exponentials to calculate the parameters of the promoter model. In absence of noise, all the model parameters were recovered accurately and with high precision (i.e. a small confidence interval), for the three sets of parameter value used to generate the artificial data (Figure 5B, lower panels). With the 1x and 2x amount of noise, parameter recovery was still accurate, while for the 4x noise condition, some parameters were recovered with a low precision, in particular those corresponding to rapid transition rates. Overall, these simulations indicated that our analysis pipeline worked well, even with complex promoter models, and was robust with respect to noise.

### Modeling indicates that pausing is stochastic and that pauses are long-lived

We analyzed the movies produced from cells expressing different amounts of Tat and created several models describing how the HIV-1 promoter may operate. The simplest model has two promoter states, ON and OFF as shown in Figure 6A, and assumes that once initiated, RNAPII enters directly into productive elongation without a pausing step. This would likely be the case when expression of Tat is high and pausing not rate-limiting, but not when Tat is limiting or absent. We thus created a model that included a pausing step. It consisted of the same simple model with two promoter states (OFF and ON), but with initiating polymerases undergoing an obligatory pause (PAUSE), before either progressing into elongation or aborting (Figure 6A middle; model M3). Note that once the polymerase exits the pause or aborts, the promoter goes immediately back in the ON state. A large body of work indicates that Tat is promoting elongation by recruiting P-TEFb and in agreement, P-TEFb is limiting for HIV-1 transcription in the *Low Tat* and *No Tat* cells used here (Figure S1B-C). Therefore, we expected to have a high abortion rate (k_abort_) and/or a low rate of pause release (k_release_) in absence of Tat, and the opposite when Tat is abundant. Conversely, the rates of switching between the ON and OFF state should not be much affected by the amount of Tat.

**Figure 6.**
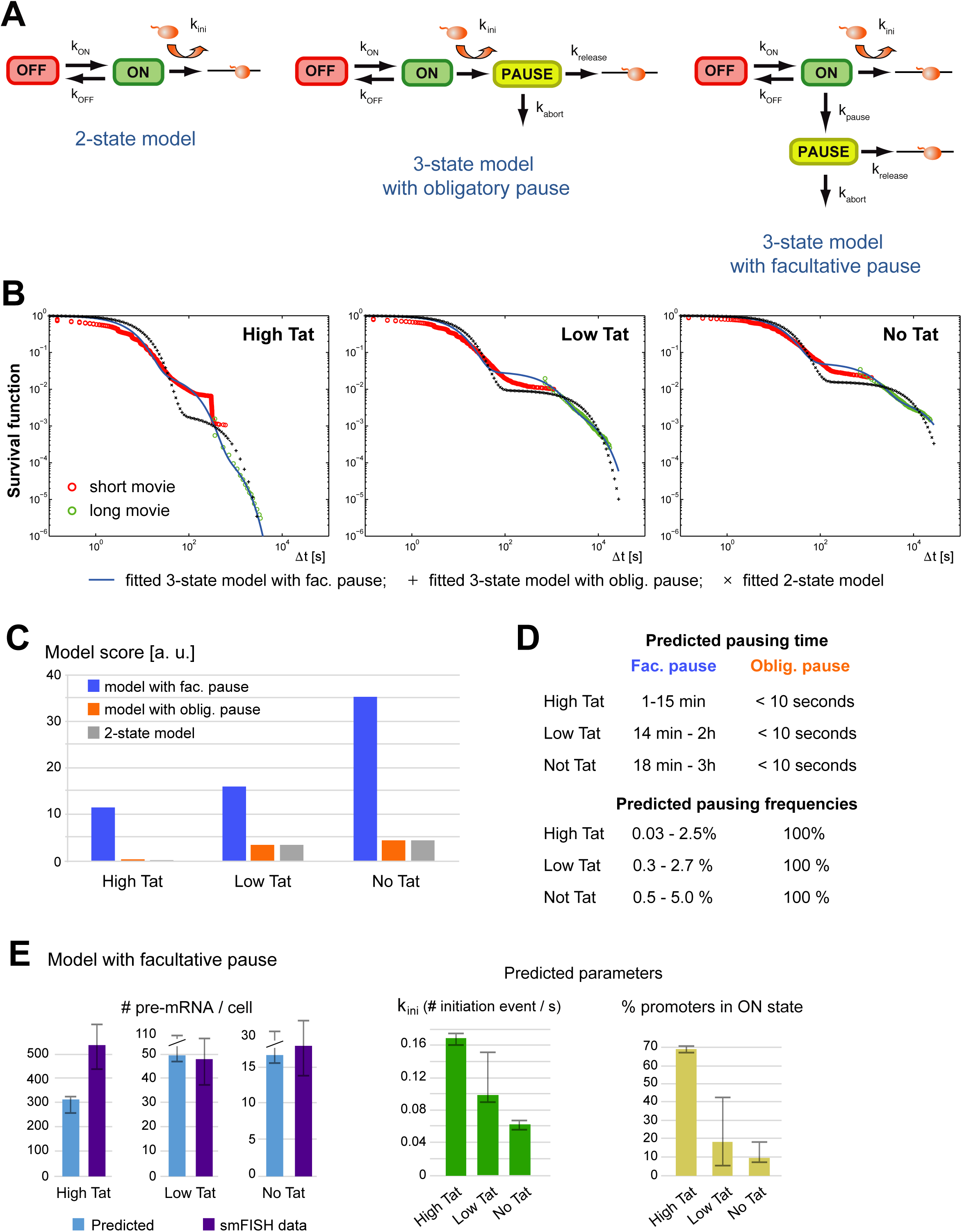
A facultative pausing model reproduces the live cell transcription data and predicts a long-lived pause. **A**-Schematics of the different models used to fit the live cell HIV-1 transcriptional data. Left: a two-state ON/OFF promoter mode; middle: a three state promoter model including an obligatory pause as traditionally represented (model M3); right: a three state promoter model with a facultative pause (model M2+). Polymerases are represented by small orange balls. **B**-Fits of the experimental survival functions. Graphs represent the survival functions reconstructed from the live cell data for the High Tat, Low Tat and No Tat conditions, with the part deriving from the short and long movies in red and green, respectively. Blue line: fit of the 3-state model with a facultative pause; “+”: fit of the 3-state model with an obligatory pause; “x”: fit with a facultative pause. x-axis: time intervals between successive initiations events, in seconds and in log_10_ scale. y-axis: probability of Δt > x (log_10_ scale). **C**-Model scores. The graph depicts the score of each model (inverse of the minimal value of the fitted Objective Function), for each of the model and cell line. **D**-Pausing characteristics predicted by the models. Top: predicted pausing times, for the relevant models and cell lines (see text for details). Bottom: predicted pausing frequencies (in %), for the indicated cell line and model. For the model with the facultative pause, the two indicated values come from the two branches of the model that could each correspond to the paused state (see model M2 in the Supplemental Text). **E**-Features of the model with the facultative pause. Left: the graphs represent the number of mRNA per cell measured by smFISH experiments (violet bars), or predicted from the model parameters (blue bars). Error bars are the standard deviation for the smFISH data (estimated from replicate measurements) and the confidence intervals for the prediction from the model. Middle: initiation rate (in s^-1^), for the three cell lines. Error bars are confidence intervals. Right: fraction of the cells with the promoter in the ON state (in %), for the three cell lines. Error bars are confidence intervals.

For the obligatory pausing model, the symbolic solution describing the distribution of polymerase waiting times is the sum of three exponentials, but with one of the five parameter being constrained and expressed as a function of the others (see Supplemental Text section 4.6). After fitting the experimental distributions of polymerase waiting times using this symbolic solution, we estimated the quality of the fit with three criteria: (i) the sum of squared residuals, evaluated from the function minimized during the fit (i.e. the objective function, with the inverse of its minimal value giving the fit score); (ii) the certainty of the value of the fitted model parameters, evaluated by their confidence intervals; (iii) the realistic nature of the parameter values, and in particular the pausing times and the effects of Tat. According to these considerations, the fit of the 3 state model with an obligatory pause was poor, and this was the case of all the Tat cell lines. First, the model scores were low and not better than the simple 2 state model without pause, even in the *Low Tat* / *No Tat* cell lines were P-TEFb recruitment limits viral transcription (Figure 6B-C). Second, the uncertainty in some parameter was high, as shown by the large confidence intervals of the parameters of the fitted exponentials (see Table 4 of Supplemental Text). Third, the pausing time, estimated from the rates of pause release and transcription abortion, was short (Figure 6D; less than 10 seconds whether Tat was present or not), while most of the regulation induced by Tat occurred at the transition between the ON and OFF state and not pausing (see Figure 17 of the Supplemental Text). It is also interesting to note that the fitted abortion rate was found to be > 100 fold faster than the rate of pause release (Figure 17 of Supplemental Text). Because the promoter goes directly to the ON state upon abortion or pause release, a high abortion or release rate creates a collapse between the ON and PAUSE states and therefore simplifies the 3 state model with pause into a simple 2 state ON/OFF model without pause. This explains why these two models have identical scores and fitted survival functions (Figure 6B, compare curves with ‘+’ and ‘x’). In order to try improving the model with obligatory pause, we made a four state model having two successive OFF states, one ON state and an obligatory pause (Model M4; see Figure 18 of Supplemental Text). This model fitted the data better and had a better overall score (see value of the objective function in Table 5 of the Supplemental text). However, it suffered from similar flaws as the previous model (see Figure 18 of Supplemental Text): (i) short pausing time whether Tat was present or not (<10 s); and (ii) high abortion rates, which similarly collapsed the 4 state model with an obligatory pause into a 3 state model without pause. Overall, increasing the number of OFF states in the model with an obligatory pause still yields short pausing times not regulated by Tat. Thus, an obligatory pause does not provide a benefit over a model without pause, with most of the effect of Tat occurring at the level of transitions between OFF and ON states. It is important to realize that given the high degree of bursting without Tat, with polymerases rapidly succeeding one another to form convoys during periods of gene activity (Figure 2), an obligatory pause necessarily means that pausing is short. In addition, since Tat mainly affects long inactive periods (Figure 3B), short pauses mean that the regulation by Tat cannot be on pausing, but rather on other steps able to produce long OFF periods. Hence, the occurrence of polymerase convoys in absence of Tat implies that an obligatory pause cannot be the step regulated by Tat to increase transcription.

This questioned the validity of the model and we thus sought for alternatives. In the previous models, pausing is an obligatory step, but it could be imagined that pausing is a facultative step, for instance if entry into the pause is stochastic. In this case, initiating polymerases have the choice of either directly progressing into productive elongation or entering a paused state, from which they can exit by either aborting or entering elongation (Figure 6A right, model M2+). To test this model, we first used a simplified variant of model M2+, in which polymerases systematically abort when exiting a facultative pause (model M2, see Figure 1 of the Supplemental Text). This model could fit the data from all the three cell lines, *High Tat, Low Tat* and *No Tat* (Figure 6B), with scores higher than the 3- or 4-state models with an obligatory pause (Figure 6C; Table 5 of Supplemental Text). Moreover, all parameters had a high precision with small confidence intervals (see Table 2 of Supplemental Text), and the model correctly predicted the number of pre-mRNA per cell (Figure 6E), with only a slight under-estimation for the *High Tat* cells. The fitted parameters indicate that pausing is infrequent, even in cells lacking Tat (Figure 6D). This implies that the fate of the paused polymerase will only marginally affect the promoter output, indicating that models in which the paused polymerase enters productive elongation would give similar results. Because the simplified model M2 is symmetrical, it is not possible to determine with certainty which parameters correspond to the ON-OFF transition, and which correspond to the ON-facultative pause. Nevertheless, both possibilities indicate a long pausing time from 15 minutes to 3h in *No Tat* cells, which is regulated by Tat as it decreases to either 1 or 15 minutes in *High Tat* cells. Pausing is also always predicted to be infrequent, varying from one every 20 to 180 polymerases in *No Tat* cells, down to one every 40 to 3900 in *High Tat* cells (Figure 6D).

### Measurement of pausing duration by biochemical approaches

To further assess models with obligatory or stochastic pausing, we attempted to test their most discriminative prediction. Obligatory pausing predicts a pausing time in the second range, while facultative pausing predicts a duration in the hour or sub-hour range (Figure 6D). Pausing duration can be estimated by measuring RNAPII residency time, and this can be achieved by performing chromatin immunoprecipitation (ChIP) during a time-course with Triptolide, a drug that inhibits TFIIH and prevents loading new polymerases without removing the ones that already initiated. We treated *High Tat* and *No Tat* cells with Triptolide for up to an hour and performed an RNAPII ChIP experiment. We analyzed the HIV-1 promoter as well as the GAPDH promoter, as a constitutively active control gene (Figure 7A). In the *High Tat* cells, similar levels of RNAPII were found on both the GAPDH and the viral promoters, while about 6-fold less polymerases was found on the HIV-1 promoter in absence of Tat, consistent with previous results (Figure S3; ^49^). Most importantly, treatment with Triptolide led to the rapid disappearance of RNAPII at the GAPDH promoter, with only ∼20% of the signal remaining after 10 min of treatment (Figure 7A). Interestingly, the kinetics observed at the HIV-1 promoter were dependent on Tat. In *High Tat* cells, the RNAPII signal also decreased rapidly and this was consistent with the rapid succession of polymerase firing that we measured in live cells (on every 4-6 seconds; ^28^). In contrast, the polymerases remained associated a much longer time with the viral promoter in absence of Tat, with 88% of the signal remaining after 10 minutes of treatment (Figure 7A). Extrapolation of the half-life of the promoter-associated polymerases indicated 10 minutes for the GAPDH promoter and for the HIV-1 promoter when Tat levels are high. However, this half-life raised to 38 minutes for the HIV-1 promoter when Tat was absent, consistent with a long pause. These long values may moreover be underestimated as hour-long treatment with Triptolide were shown to cause degradation of RNA polymerase II in human cells ^50^. Altogether, these data verify a key discriminative prediction of the facultative pausing model, namely that paused polymerases exhibit a half-life in the sub hour range and not in the second range as expected from an obligatory pausing scenario.

**Figure 7.**
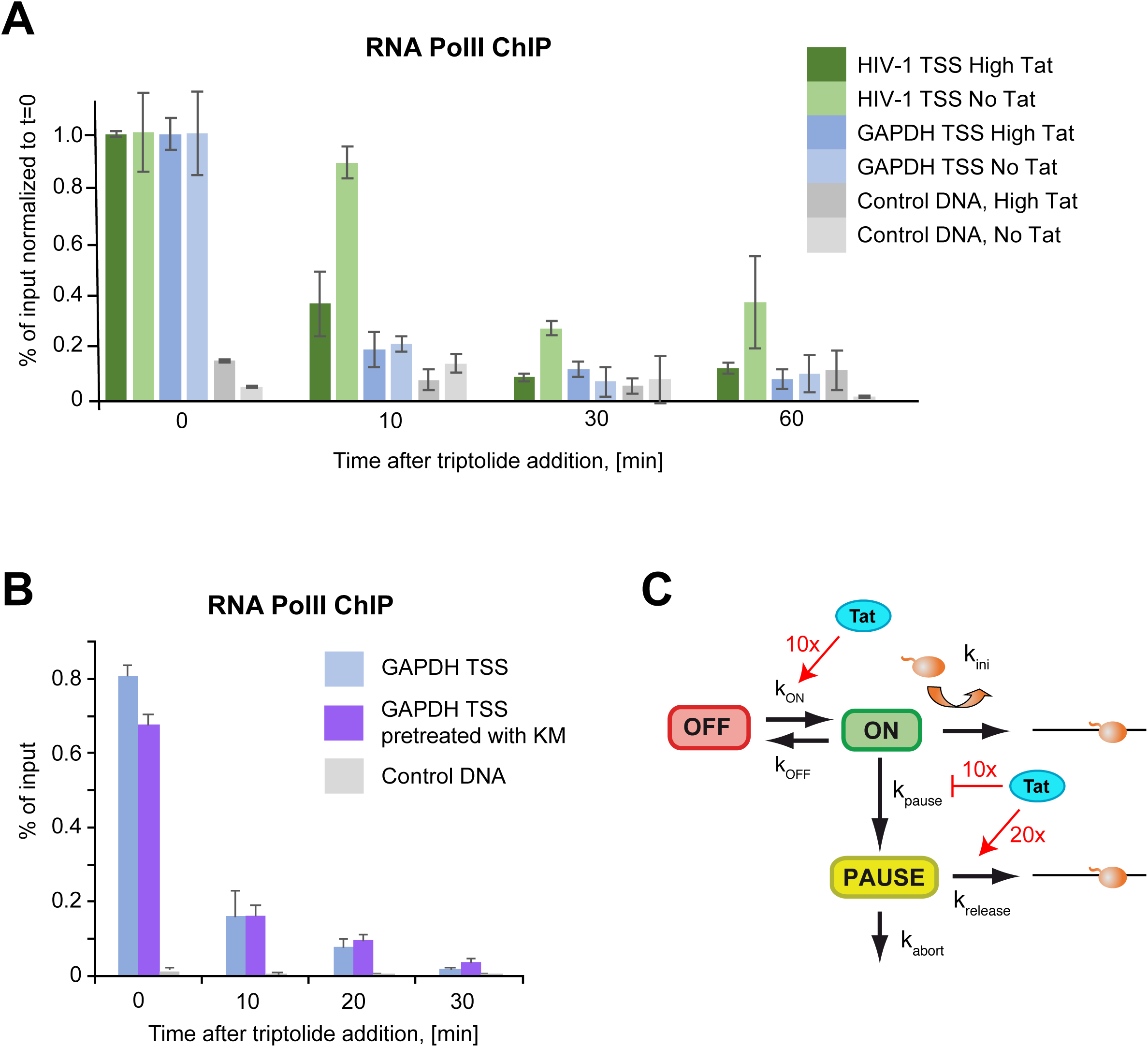
Biochemical measurements indicate a long-lived paused state at the HIV-1 promoter. **A**-Residency time of RNA polymerase II at the HIV-1 promoter. The graph depicts the RNA polymerase II ChIP signals at the HIV-1 and GAPDH promoters during a Triptolide time course experiment, for the High Tat and No Tat cell lines. GAPDH TSS: transcription start site of the human GAPDH gene; HIV-1 TSS: transcription start site of the HIV-1 promoter; Control DNA: a non-transcribed genomic locus. ChIP signals were measure by qPCR and values are expressed as percent of input and normalized to the zero time point. For the control genomic regions (Control DNA), values are normalized to that of GAPDH TSS at time zero. **B**-Effect of pTEFb inhibition on the residency time of RNA polymerase II at the GAPDH promoter. Legend as in panel A, except that the KM sample was pretreated with the Cdk9 inhibitor KM05382 for 2h before triptolide addition. **C**-Model depicting the dynamics of the HIV-1 promoter and highlighting the positive and negative effects of Tat. The numbers are from the facultative pausing model fitted to the High Tat and No Tat data (see Figure 6C and supplemental text, Table 3). The model with facultative pausing has two symmetrical branches (see model M2 in the Supplemental Text), and each branch of the model could correspond to the paused state. The values indicated attribute the pause state to the branch that is most affected by the presence of Tat.

Next, we wished to determine whether long pausing time requires a specific feature of the HIV-1 promoter or could be induced at any promoter by depleting P-TEFb. We thus repeated the GAPDH RNAPII ChIP time course, but pretreated cell with the Cdk9 inhibitor KM05382 for 2h before performing the Triptolide time course. The residency time of RNAPII at the GAPDH promoter was similarly short whether cells were pretreated with KM05382 or not (Figure 7B), indicating that the lack of P-TEFb activity is not sufficient in itself to induce long pauses. This suggests that the HIV-1 promoter likely has additional features that specify this property.

## Discussion

Cells latently infected with HIV-1 prevent patients from clearing the virus, as the stochastic activation of these cells can re-establish viral propagation ^32,34^. Latent cells do not express the viral genome and pausing of RNA polymerases at the viral promoter is a key block that prevents HIV-1 transcription ^35-38^. Pausing thus plays a fundamental role in HIV-1 biology, and yet, how it contributes to bursting and stochastic reactivation of the virus is not known. Here, we harnessed the power of single molecule transcriptional imaging and modeling to study how pausing affects HIV-1 transcription in single cells. We find that pausing is a stochastic process, and modeling as well as biochemical experiments indicate that it is long lived inhibitory state that impacts only a small fraction of the initiating polymerases. Stochastic pausing therefore generates viral transcriptional bursts in absence of Tat, which may cause viral reactivation, latency exit and viral rebounds in patients.

### A frequentist approach accurately and robustly model transcriptional fluctuations

Single molecule transcription imaging is a powerful technique that becomes indispensable for understanding transcriptional regulation in vivo. However, the signal produced by this technique integrates processes with widely distributed timescales, and not directly accessible by simple data processing. Hence, new modeling methods are needed to cope with the multiscale nature of transcription. To this end, we developed a new machine learning and modeling method. Using numerical deconvolution, this approach generates a time map of transcriptional initiation events indicating, for each transcription site, when RNAPII molecules start producing an mRNA. This feature is unique in our analysis pipeline and not available in other approaches that directly fit a particular transcription model to experimental data ^29,45-48^. Our method generates a multiscale cumulative distribution function of polymerase waiting times, which separate successive transcription initiation events. This distribution function has the unique advantage of integrating temporal information on transcriptional processes with an unprecedented dynamic range from seconds to days. Moreover, we have analytically solved the inverse problem consisting in computing the model parameters as a function of the waiting time distribution, for a large number of models. By allowing easy and quick comparison of many different models of promoter dynamics, this method removes a bottleneck essential for hypothesis testing in gene regulation studies.

### Polymerase pausing generates long-lived inactive states that limit HIV-1 transcription

P-TEFb is an essential elongation factor that is required for both the basal and Tat-induced activity of the HIV-1 promoter ^11,35,36^. By default, the HIV-1 promoter leads to pausing and inefficient elongation, and Tat functions as a promoter specific elongation factor by recruiting P-TEFb to the nascent viral RNAs. When Tat is present in saturating concentrations, we observe that polymerases initiate rapidly, one after another (every 4-6s in average; ^28^). This indicates that the maturation of initiating polymerases into a processive complex is rapid, in agreement with the fact that P-TEFb recruitment is not rate-limiting when Tat is abundant. When Tat is limiting or absent, we observe a biphasic behavior. HIV-1 promoters are mostly inactive, and yet sometimes transcribe the viral genome in brief pulses containing tens of polymerases. These polymerases are fired in rapid succession (one every 7-15s), and they form convoys resembling the ones observed when Tat is saturating. Modelling the imaging data in *Low Tat* and *No Tat* cells confirms this biphasic behavior and further indicates that in average, 20-35 polymerases initiate during active periods of 5 minutes, followed by inactive periods of 20 minutes or 3h. Given that pausing is limiting transcription in the absence of Tat, these long inactive periods are likely caused by long-lived pauses at the HIV-1 promoter. Indeed, direct measurements of RNAPII residency time at the HIV-1 promoter indicate that the absence of Tat generates long pauses in the sub-hour range, which are therefore responsible for at least some of the long periods without viral transcription.

Recent genome-wide data obtained in Drosophila with Triptolide time-course experiments indicate that the half-life of polymerases at cellular promoters varies from less than a minute to about an hour ^19-23^. Analysis of a series of promoter variants further indicates that an initiator element with a G at position +2 is a key determinant of long pausing time (>40 minutes; ^51^). It is not known whether this rule also applies to vertebrates, but it is worth noting that the HIV-1 promoter has an unusual initiator element required for Tat activation that contains a G at +2 ^52^. Moreover, inhibiting P-TEFb does not generate long pauses at the GAPDH promoter, suggesting that some promoter specific features exist. In the future, it will be interesting to determine whether long-lived and short-lived paused polymerases have a similar 3D structure. Indeed, recent data in NELF KO cells suggests that polymerases can have several pausing sites and states ^53^. Because of their half-life, long-lived paused polymerase may display additional features such as backtracking or other stabilizing properties, and backtracking was indeed shown to occur at the pause site in the case of HIV-1 ^54^. Long-lived paused polymerases are especially interesting because of their properties, which effectively limit transcription but maintain the promoter in an open state ^22^.

### Stochastic pausing generates transcriptional bursting

In the traditional model of transcription initiation, polymerase pausing is an obligatory step during the formation of the elongation complex ^2,55^. In contrast, our live cell data on HIV-1 transcription suggest that pausing is a stochastic event that occurs rarely: 1 every 20-180 polymerase in absence of Tat and down to 1 every 40-3900 when Tat is abundant. A model reconciling these views would be the existence of two fates during pausing: a pause could lead to either rapid enzyme maturation, or to a long-lived inactive state that would inhibit further transcription. In this scenario, the polymerases initiating at the HIV-1 promoter would mature into a processive elongation complex but would have a low probability of entering a long-lived paused state (Figure 7C). A key feature of this model is that long-lived pauses are stochastic, and this changes the nature of this process as long-lived pauses would not be a step required for proper polymerase maturation but an inhibitory state preventing transcription. In essence, stochastic long-lived pauses are analogous to an inactive promoter state (Figure 7C). In the case of HIV-1, long-lived pauses would be a key regulatory step in transcriptional regulation, and by ensuring an efficient recruitment of P-TEFb, Tat would drastically reduce the probability of long inhibitory pauses (see model in Figure 7C). This is consistent with the fact that the HIV-1 promoter is fully occupied in a model of latent cells ^56^, even if in some cases Tat can slightly enhance PIC occupancy ^49^. It is also consistent with the known function of Tat as a P-TEFb/SEC recruiting factor, with a major function in reducing pausing.

The basal activity of the HIV-1 promoter requires P-TEFb and it is surprising that the factors responsible for P-TEFb recruitment in absence of Tat allow for the firing of a series of polymerases before switching back to a long inactive state. Indeed, the HIV-1 promoter is active for periods of ∼ 5 minutes in absence of Tat, firing 20 polymerases on average. A possibility to explain this behavior would be a switching mechanism, in which P-TEFb would be present and active for several minutes at the HIV-1 promoter, and then leave for long time periods. Our data show that NF-κB is not involved in the basal transcriptional activity of HIV-1 in our cellular system, and we can thus rule out sporadic activation of this pathway as a cause of transcriptional bursts in absence of Tat. Another possibility would involve the diffusion dynamics of P-TEFb. Indeed, it has been shown that P-TEFb is a local explorer that repetitively visit the same location ^57^, and recent data further suggest that P-TEFb undergoes transient liquid-liquid phase transitions ^58^. FRAP studies showed that the residency time of P-TEFb is 11 seconds at the HIV-1 promoter in absence of Tat ^59^, and 55 seconds at the transcription site of a CMV-based reporter ^58^. While this is too short to explain the 5 minutes active periods without Tat, single particle tracking of P-TEFb subunits indicate a wide range of binding times ^58^. Moreover, P-TEFb also might exchange rapidly from longer-lasting liquid condensates. It is also possible that other phenomena are responsible for P-TEFb recruitment, or that long pauses arise from an inherently stochastic and inefficient process.

The stochastic nature of long-lived polymerase pausing and their low probability has important consequences for HIV-1 pathogenesis. There are evidences that the stochastic activation of the viral promoter is responsible for the stochasticity of latency exit, at least in part ^32-34,37^. Moreover, latent viruses do not express Tat or at very low levels ^35,36^, and we show that in these conditions the spontaneous release of a long-lived pause leads to the synthesis of a large series of viral RNAs. In some cases, this may be sufficient to activate the viral promoter and to initiate the Tat positive feedback loop, leading to acute viral replication. The stochastic nature of long-lived pausing may thus be an important feature of HIV-1 regulation that favorizes spontaneous latency exit ^34,37,38^. It is also possible that even if the viral RNAs produced do not initiate the Tat-feedback loop, they may still produce a small amount of viral particles, which may infect naive cells and could thus participate in the viral rebounds or viremia blips seen in patients. It is also important to note that quiescent memory T cells have a low P-TEFb activity ^35,36^, possibly leading to very long periods without HIV-1 transcription. Finally, stochastic pausing has also been reported in developing Drosophila embryos, where it may finely tune gene expression after zygotic genome activation (see accompanying paper). Stochastic pausing may be a general property of cellular promoters important for gene regulation.

## Methods

### Cell culture and drug treatments

HeLa Flp-in H9 cells (a kind gift of S. Emiliani) were maintained in DMEM supplemented with 10% fetal bovine serum, penicillin/streptomycin (10 U/ml) and glutamin (2.9 mg/ml), in a humidified CO_2_ incubator at 37°C. Cells were transfected with the indicated plasmids with JetPrime (Polyplus), following manufacturer recommendations. Drugs were used at the following concentrations: Triptolide, 1 μM; KM05382 100 μM; BAY11-7082, 2 μM.

Stable expression of MCP-GFP was achieved by retroviral-mediated integration of a self-inactivating vector containing an internal ubiquitin promoter (as described in ^28^). The MCP used dimerizes in solution and contained the deltaFG deletion, the V29I mutation, and an SV40 NLS. MCP-GFP expressing cells were grown as pool of clones and FACS-sorted to select cells expressing low levels of fluorescence. Isogenic stable cell lines expressing the 128xMS2 HIV-1 reporter gene were created using the Flp-In system and a HeLa H9 strain expressing various levels of Tat (see below) and MCP-GFP. Flp-In integrants were selected on hygromycin (150 μg/ml). For each construct, several individual clones were picked and analyzed by in situ hybridization.

*No Tat* cells expressed the 128xMS2 HIV-1 reporter gene but did not express any Tat protein. To obtain low level of Tat expression, a Tat-Flag fused to an Auxin-inducible degron (AID) and cloned as a second cistron after auxin receptor F-box protein AFB2 and instead of GFP in a previously described vector AAV-CAGGS-eGFP ^60^. The resulting vector was integrated in genomic AAVS1 site using CRISPR-Cas9 and clones were selected using puromycin as described ^60^. Cells were not treated with Auxin.

*High Tat* cells ^28^ were created using the plasmid pSpoII-Tat. In this plasmid, the CMV promoter transcribes a Tat-Flag cDNA followed by an IRES-Neo selectable marker. Following Neomycin selection (400 μg/ml), expression levels of individual clones were verified by western blotting and by immunofluorescence to ensure homogeneity both between clones and between cells of a clone.

### Plasmids

Sequences of the plasmids are available upon request. The 128xMS2 HIV-1 reporter and High Tat expression vector were described previously ^28^. AAV-CAGGS-eGFP vector, used to obtain low Tat cells, Cas9 encoding vector and AAVS1-site targeting RNA-guides were obtained from Dr. G. M. Church ^60^. pcDNA3-CDK9-GFP and pcDNA3-CyclinT1-GFP plasmids were obtained by Gateway technology, CDK9 and Cyclin-T1 were amplified by PCR from the vectors provided by Dr. L. Lania ^61^. pHR-SFFV-dCas9-BFP plasmid used for CDK9 cloning is #46911 from Addgene. The RNA guides were cloned in a home-made U6 expression vector with an optimized guide RNA scaffold ^62^.

### dCas9 tethering and pTEFb overexpression

For P-TEFb overexpression, Hela 128xMS2 HIV-1 No Tat cells without MCP-GFP were plated on coverslips and the next day transfected with CDK9-GFP, Cyclin-T1-GFP, or both, using jetprime (polyplus). pBluescript was used as a negative control and GFP-Tat as a positive control. 24-hours after transfection cells were fixed and the reporter RNA was detected by smFISH with Cy3-labeled fluorescent probes against MS2 repeats, the RNA expression was scored in transfected GFP-positive cells.

For CDK9 tethering, RfB gateway cassette was cloned in pHR-SFFV-dCas9-BFP between dCas9 and BFP. CDK9 was next introduced by LR recombination. The resulting plasmid pHR-SFFV-dCas9-CDK9-BFP was transfected in Hela *No Tat* cells without MCP-GFP together with 3 RNA guides encoding plasmids as described above. pHR-SFFV-dCas9-CDK9-BFP without guides and pHR-SFFV-dCas9-BFP were used as controls. 24 hours after transfection cells were fixed and subjected to smFISH with probes against 128xMS2. The numbers of RNA molecules in BFP-positive cells were counted using FISH-QUANT ^63,64^. The sequences of RNA guides were as follows CCGCCTAGCATTTCATCACG, CCACGTGATGAAATGCTAGG, TGCTACAAGGGACTTTCCGC.

### SmFISH and RNA quantification

SmFISH was performed as previously described ^28^, with a mix of 10 fluorescent oligos hybridizing against the MS2×32 repeat, each oligo containing four molecules of Cy3. Since each oligo bound four times across the 128xMS2 repeat, each molecule of pre-mRNA hybridized with 40 oligos, thereby providing excellent single molecule detection and signal-to-noise ratios.

To obtain the number of nascent and released pre-mRNAs per cell and the distribution of this parameter in the cell population, cells processed for smFISH were imaged on a ZEISS Axioimager Z1 wide-field microscope (63X∼, NA 1.4; 40X∼, NA 1.3), equipped with an sCMOs Zyla 4 .2 camera (Andor) and controlled by MetaMorph (Universal Imaging). 3D image stacks were collected with a Z-spacing of 0.3 μm. Figures were prepared with Image J, Photoshop and Illustrator (Adobe), and graphs were generated with R or MatLab.

Raw, 3D smFISH images were analyzed to count the number of pre-mRNA per nuclei, using populations of >300 cells per experiment. Briefly, nuclei were segmented using the DAPI signal with Imjoy ^65^, and transcription sites (TS) were identified manually. Isolated pre-mRNA molecules located in the nucleoplasm were then detected with *FISH-quant* ^63,64^, after manual thresholding of Laplacian on Gaussian filtered image. This defined the PSF and the total light intensity of single molecules, which were averaged to obtain an average PSF. The average PSF of single RNA molecule was used to determine the number of nascent pre-mRNA molecules at the TS.

### Live cell imaging

Cells were plated on 25 mm diameter coverslips (0.17 mm thick) in non-fluorescent media (DMEM gfp-2 with rutin; Evrogen). Coverslips were mounted in a temperature-controlled chamber with CO_2_ and imaged on an inverted OMXv3 Deltavision microscope in time-lapse mode. A 100x, NA 1.4 objective was used, with an intermediate 2X lens and an Evolve 512×512 EMCCD camera (Photometrics). Stacks of 11 planes with a z-spacing of 0.6 μm were acquired. This spacing still allowed accurate PSF determination without excessive oversampling. Illuminating light and exposure time were set to the lowest values that still allowed visualization of single molecules of pre-mRNAs (laser at 1% of full power, exposure of 15 ms per plane). This minimizes bleaching and maximizes the number of frames that can be collected. Yet, it guarantees that transcription can be detected early on, when one or a few nascent chains are in the process of being transcribed. For short movies, one stack was recorded every 3 seconds for 15 to 20 minutes. For long movies, one stack was recorded every three minutes for 8 hours.

### Quantification of short movies

Extract the TS signal in the short movies was done as previously described ^28^. We manually defined the nuclear outline and the region within which the TS is visible. The stack was corrected for photobleaching by measuring the fluorescence loss of the entire nucleus and fitting this curve with a sum of three exponentials. This fitted curve was then used to renormalize each time-point such that its nuclear intensity was equal to the intensity of the first time-point. We then filtered the image with a 2-state Gaussian filter. First, the image was convolved with a larger kernel to obtain a background image, which was then subtracted from the original image before the quantification is performed. Second, the background-subtracted image was smoothened with a smaller Kernel, which enhances the SNR of single particles to facilitate spot pre-detection.

We then pre-detected the position of the TS in each frame of the filtered image by determining in the user-specified region the brightest pixel above a user-defined threshold. If no pixel was above the threshold, the last known TS position was used. Pre-detected position was manually inspected and corrected. Then the TS signal was fitted with a 3D Gaussian estimating its standard deviation *σ*_*xz*_ and *σ*_*z*_, amplitude, background, and position. We performed two rounds of fitting: in the first round all fitting parameters were unconstrained. In the second round, the allowed range was restricted for some parameters, to reduce large fluctuations in the estimates especially for the frames with a dim or no detectable TS. More specifically, the *σ*_*xz*_ and *σ*_*z*_ were restricted to the estimated median value +/- standard deviation from the frames where the TS could be pre-detected, and the background was restricted to the median value. The TS intensity was finally quantified by estimating the integrated intensity above background expressed in arbitrary intensity units.

With the live cell acquisition settings, the illumination power was low and we could not reliably detect all individual molecules. We therefore collected right after the end of the movie one 3D stack – termed calibration stack - with increased laser intensity (50% of max intensity, compared to 1% for the movie), which allowed reliable detection of individual RNA molecules. We also collected slices with a smaller z-spacing for a better quantification accuracy (21 slices every 300 nm). Quantification of TS site intensity in the calibration stack was done with *FISH-quant* as follows: (a) when calculating the averaged image of single RNA molecules, we subtracted the estimated background from each cell to minimize the impact of the different backgrounds; (b) when quantifying the TS in a given cell, we rescaled the average image of single RNA molecules such that it had the same integrated intensity as the molecules detected in the analyzed cell.

To calibrate the TS intensities in the entire movie, i.e. to express the TS intensity as a number of equivalent full-length transcripts, we used the fact that the last movie frame was acquired at the same time as the calibration stack. We then normalized the extracted TS intensity in the movies, *I*_*MS2*_, to get the nascent counts *N*_*nasc;calib*_:

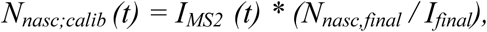

where *N*_*nasc,final*_ stands for the estimated number of nascent transcripts in the calibration stack and *I*_*final*_ for the averaged intensity of the last 4 frames. Note that the approach was limited to movies where the TS was active at the movie end since otherwise its intensity could not be quantified. More than 100 cells were used in each condition.

### Quantification of long movies

To quantify the long movies acquired at low frames rate (one 3D stack per 3 minutes), we used *ON-quant* ^28^, a rapid analysis tool that identified the ON and OFF periods and measured their length. This did not require an absolute quantification of the number of nascent pre-mRNAs and we therefore defined an intensity threshold, based on the mean intensity of single molecules, under which a TS is considered to be silent, and above which a TS is considered to be active. This threshold corresponded to the intensity of 1.5 pre-mRNA. For each cell line between 100 and 150 cells were analyzed.

### Mathematical modelling

A detailed description of the algorithm can be found in the Supplemental Text, and the software algorithms are in the Supplemental file.

#### Deconvolution and RNAPII Positioning

The RNAPII positions were found by combining a genetic algorithm with a local optimisation procedure. Before initiation of the analysis algorithm, several key parameters were established. The RNAPII elongation speed was fixed at 67 bp/s ^28^. The reporter construct transcript was divided into three sections consisting of the pre-MS2 fragment (PRE=700 bp), 128xMS2 loops (SEQ=2900 bp), and post-MS2 fragment (POST=1600 bp). An extra time *P*_*poly*_=100s was added to POST, corresponding to the polyadenylation signal (during this time the polymerase has finished transcription and waits on the transcription site). The temporal resolution of short movies was 3 s/frame. This frame rate is sufficient to detect processes that occur on the order of seconds.

The possible polymerase positions were discretized using a step of 30 bp. This step was chosen as it is smaller than the minimum polymerase spacing and large enough to have a reasonable computation time. For a movie of 20 min length this choice corresponds to a maximum number of 2680 positions. The deconvolution algorithm was implemented in Matlab R2020a using Global Optimization and Parallel Computing Toolboxes for optimizing RNAPII positions in parallel for all nuclei in a collection of movies. The resulting positions are stored for analysis in the further steps of our computational pipeline. The deconvolution step is common to all of the MS2 data analysis pipelines.

#### Long movies waiting time distribution

For long movies, the low resolution (3min) does not allow RNAPII positioning. In this case we binarize the signal by considering that the transcription site is active or inactive if the measured intensity is above or below a threshold level, respectively. The inactive intervals indicate long waiting times between successive polymerases. The active intervals are used to estimate the probability that waiting times are larger than the movie resolution (see Supplemental Text).

#### Multi-exponential regression fitting of the survival function and model reverse engineering using the survival function

Data from several short movies corresponding to the same phenotypes was first pooled together. Waiting times were extracted as differences between successive RNAPII positions from all the resulting traces and the corresponding data was used to estimate the nonparametric cumulative short movie distribution function by the Meyer-Kaplan method. Data from long movies and the same phenotype are also pooled and generate the nonparametric cumulative long movie distribution function. The two conditional distribution functions are fitted together into a multiscale cumulative distribution function using the total probability theorem and estimates of two parameters *p*_*l*_ and *p*_*s*_, representing the probabilities that waiting times are longer than the long movie resolution, and longer than the length of the short movie, respectively (see Figure 4 and Supplemental Text for details).

Then, a multi-exponential regression fitting of the multiscale distribution function produced a set of 2N-1 distribution parameters, where N is the number of exponentials in the regression procedure (3 for N=2 and 5 for N=3). The regression procedure was initiated with multiple log-uniformly distributed initial guesses and followed by local gradient optimisation. It resulted in a best-fit solution with additional suboptimal solutions (local optima with objective function value larger than the best fit).

The 2N-1 distribution parameters can be computed from the 2N-1 kinetic parameters of a N state transcriptional bursting model. Conversely, a symbolic solution for the inverse problem was obtained, allowing computation of the kinetic parameters from the distribution parameters and reverse engineering of the transcriptional bursting model. In particular, it is possible to know exactly when the inverse problem is well-posed, i.e. there is a unique solution in terms of kinetic parameters for any given distribution parameters in a domain.

The transcriptional bursting models used in this paper are as following:

For N=2, there were 3 distribution parameters and 3 kinetic parameters.

The distribution parameters are *A*_1_, *λ*_1_, *λ*_2_, defining the survival function

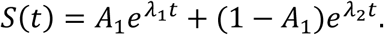

The solution of the inverse problem for the ON-OFF telegraph model (Figure 6A) is

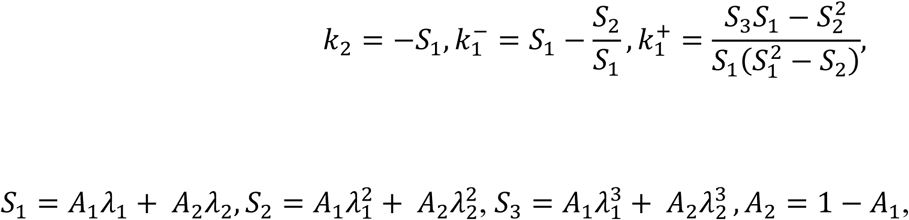

where 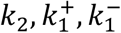 are the initiation rate, the OFF to ON and ON to OFF transition rates, respectively.

For N=3, there were 5 distribution parameters and 5 kinetic parameters.

The distribution parameters are *A*_1_, *A*_2_, *λ*_1_, *λ*_2_, *λ*_3_, defining the survival function

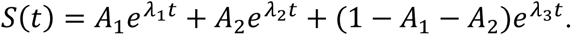

The inverse problem has a unique solution for the 3 state model (stochastic, facultative pause) with one OFF state, one PAUSE state and one ON state (Figure 6A, model M2 of Supplemental Text). Note that the kinetic parameter of Figure 6A (model M2+) are noted below as follow for model M2: k_ini_ = k3; k_pause_ = k2^-^; k_abort_ = k2^+^; k_release_ = 0.

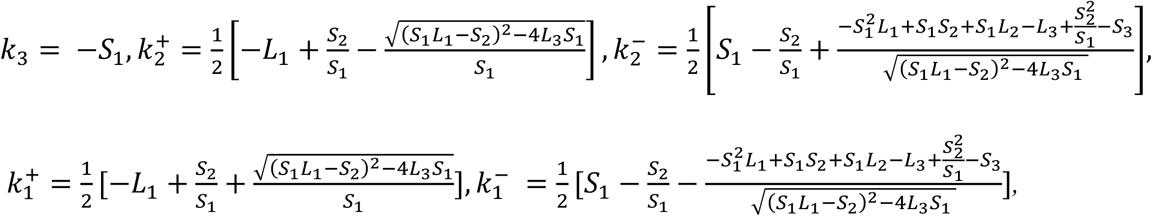

where

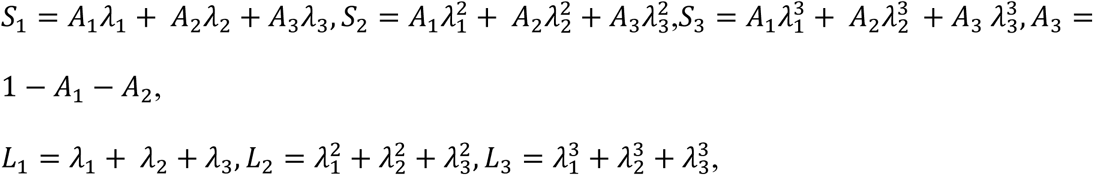

and 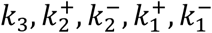 are the transcription initiation, OFF to ON, ON to OFF, PAUSE to ON, and ON to PAUSE rates, respectively.

Duration of the ON, OFF, and PAUSE states can be calculated thusly:

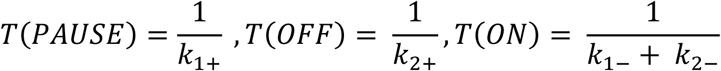

For this model, the steady state probability to be in a given promoter state is

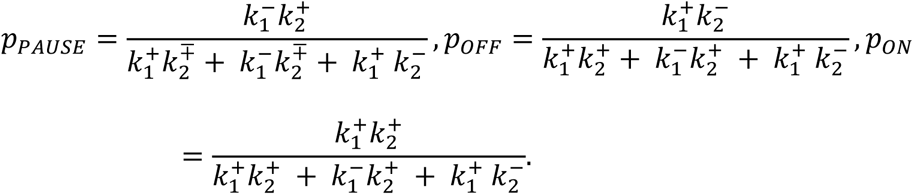

The alternative 3 state model with obligatory pause (Figure 6A, model M3) satisfies the following relation among distribution parameters (see Supplemental Text for a proof):

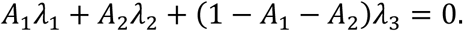

This means that only 4 and not 5 distribution parameters are free, which further constrains the three exponential fitting. In order to infer this model, a constrained fitting was performed but the bad quality of fitting recommended rejection of the model (Figure 6B-C; see results).

#### Testing the method with artificial data

The entire computational pipeline was tested using artificial data. Artificial traces were generated by simulating the model using the Gillespie algorithm with parameter sets similar to those identified from data. The simulations generated artificial polymerase positions, from which a first version of the signal was computed by convolution. The results are provided in Figure 5 and Supplemental Text.

#### Error intervals

Distribution parameters result from multi-exponential regression fitting using gradient methods with multiple initial data. These optimization methods provide a best fit (global optimum) but also suboptimal parameter values. Using an overflow ratio (a number larger than one, in our case 2) to restrict the number of suboptimal solutions, we define boundaries of the error interval as the minimum and maximum parameter value compatible with an objective function less than the best fit times the overflow.

#### mRNA levels

Steady state mRNA levels can be computed from the parameters of the multi-exponential fit. We showed in the Supplemental Text that:

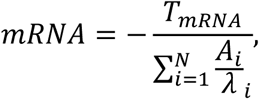

where *T*_*mRNA*_ is the mean lifetime of the mRNA. The formula is valid for all N and we have used *T*_*mRNA*_ = 45 min ^28^.

### Chromatin immunoprecipitation

*High Tat* and *No Tat* HeLa cells were treated with 1 μM of triptolide at 0, 10, 30 and 60 minutes. *High Tat* HeLa cells were treated with 100 μM of KM05382 during 1 hour followed by 1 μM of triptolide at 0, 10, 20 and 30 minutes. Cells were cross-linked by adding crosslinking solution (11% formaldehyde, 100 mM NaCl, 1 mM EDTA pH 8, 0.5 mM EGTA pH 8, 50 mM Hepes pH 7.8) directly to cultures (1% final) and incubated for 10 min at room temperature. Then, 250 mM final glycine was added, and cultures were incubated for 5 min at room temperature. Cells were then washed four times with cold PBS, scraped in cold PBS with Protease Inhibitor cocktail and centrifuged at 1350×g for 10 min.Crude nuclei were prepared by hypotonic lysis. The pellet was resuspended in 5 mL of BufferA (50 mM Hepes pH 8.0, 85 mM KCl, 0.5% Triton-X-100, Protease Inhibitor cocktail, 1 mM PMSF), incubated on ice for 10 min and centrifuged at 1350xg for 10 min. Then, the pellet was resuspended in 5 mL of BufferA’ (50 mM Hepes pH 8.0, 85 mM KCl, Protease Inhibitor cocktail, 1 mM PMSF) and centrifuged at 1350xg for 10 min. Finally, the pellet was resuspend in 0.9 mL of Buffer B (50 mM Tris-HCl pH 8, 1% SDS, Protease Inhibitor cocktail, 1 mM PMSF), incubated on ice for 10 min and then stored at the −80°. Pellets were sonicated at 4°C using a Bioruptor (Diagenode) to shear the chromatin to a mean length of 300 bp by repeated cycles (16 cycles of 30 s ON and 30 s OFF). After sonication cellular debris was removed by centrifugation at 20000×g for 10 min. The chromatin solution was diluted 10-fold in FA/SDS Like buffer (50 mM Hepes KOH pH 7.5, 150 mM NaCl, 1% Triton-X-100, 0.1% Na deoxycholate, Protease Inhibitor cocktail, 1 mM PMSF) and precleared for 1 hour at 4°C with 25 μl of protein G Dynabeads (Invitrogen). The precleared chromatin solution (1.5 × 106 cells) was incubated overnight with 50 μL of BSA-blocked protein G Dynabeads (previously bound with 3 ug of the corresponding antibody, POLII F-12 sc-55492 Lot K1516 Santacruz, during 1 hour at 4°C). Samples were washed once with FA/SDS buffer (50 mM Hepes KOH pH 7.5, 150 mM NaCl, 1% Triton-X-100, 0.1% Na deoxycholate, 1 mM EDTA, 0.1% SDS, Protease Inhibitor cocktail, 1 mM PMSF), three times with FA/SDS Buffer supplemented with 300mM NaCl, once with washing Buffer (10 mM Tris-HCl pH 8, 0.25 M LiCl, 1 mM EDTA, 0.5% NP40, 0.5% Na deoxycholate) and once with TE Buffer. Elution was performed adding 125 μl of Elution Buffer (25 mM Tris-HCl pH 7.5, 5 mM EDTA, 0.5% SDS) and incubating at 65°C for 25 min. The eluates were digested with 50 μg/mL of RNase A at 37°C for 30 min and with 50 μg/ml of proteinase K at 50°C for 1 h. Then, they were incubated at 65°C overnight to reverse cross-links. DNA was recovered by phenol extraction followed by a Qiaquick purification (PCR purification columns, Qiagen, Germany). Specific sequences in the immunoprecipitates were quantified by real-time PCR using the primers listed below. The signal of each sample was normalized with the average signal obtained from the input of the same sample with each pair of primers used. Each experiment was done analysing two independent biological replicates.

Primers used:

GAPDH promoter F: 5’ AAAGGCACTCCTGGAAACCT

GAPDH promoter R: 5’ GGATGGAATGAAAGGCACAC

GAPDH negative control F: 5’ CTAGCCTCCCGGGTTTCTCT

GAPDH negative control R: 5’ ACAGTCAGCCGCATCTTCTT

TSS HIV1 +92 F: 5’ GCTTCAAGTAGTGTGTGCCC

TSS HIV1 +92 R: 5’ GCTTTCAAGTCCCTGTTCGG

---

**Supplementary Figure S1.**
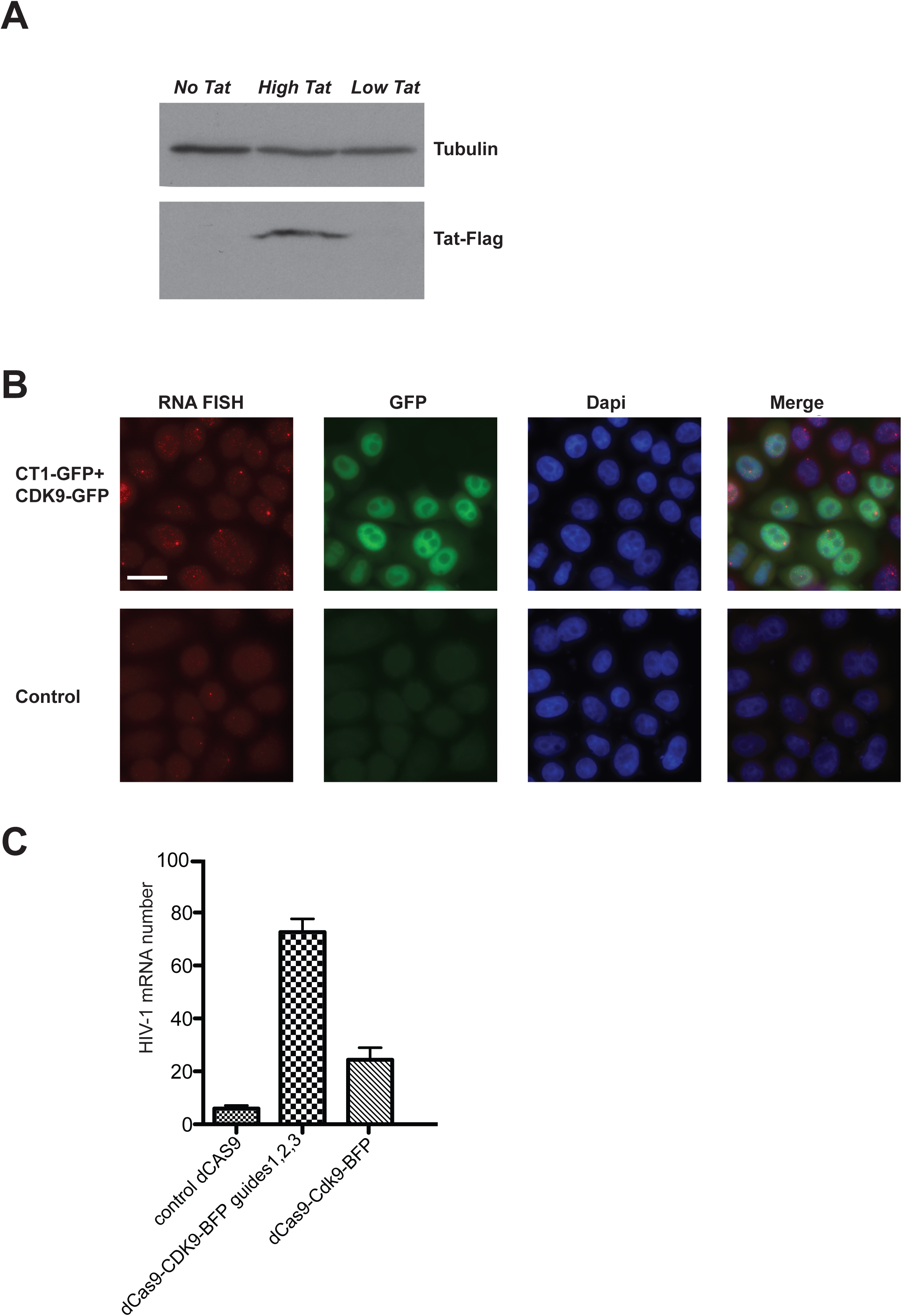
Transcriptional activation of HIV-1 128xMS2 reporter in Hela Flp-in cells is P-TEFb dependent. **A-** Western blot of the extracts of HIV-1 128xMS2 Hela cell lines with no, low and high Tat expression. Tat-Flag was detected with anti-Flag antibodies; loading control is tubulin. **B-** CDK9-GFP and cyclinT1-GFP activate transcription of the HIV-1 reporter. Fluorescent microscopy images of Hela Flp-in cells with the HIV-1 128xMS2 reporter, not expression Tat nor MCP-GFP, and co-transfected with plasmids encoding for CDK9-GFP and cyclinT1-GFP (24h after transfection). First row from the left: RNA of HIV-1 reporter detected by smFISH with Cy3 probes against 128xMS2 tag; second row: GFP signal corresponding to the cells transfected with CDK9-GFP and cyclinT1-GFP; third row: nuclear staining with dapi; last row: merge. Top panel: cells transfected with CDK9-GFP and cyclinT1-GFP. Bottom panel: control transfection with pBluescript. The scale bar is 10 μm. **C**-Tethering of CDK9 to the HIV-promoter using dCas9 leads to transcriptional activation. The histogram shows the results of mRNA counting on smFISH images 24h after transfection the Hela Flp-in HIV-1 128xMS2 no Tat cells (without MCP-GFP) with dCas9-CDK9-BFP fusion and 3 RNA guides targeting the CDK9 fusion specifically to the HIV-1 promoter (middle bar); dCas9-CDK9-BFP fusion without guides (right bar) or dCas9-BFP alone (left bar) were transfected in control experiments. On y axis is the mRNA number. Error bars are standard errors of the mean.

**Supplementary Figure S2.**
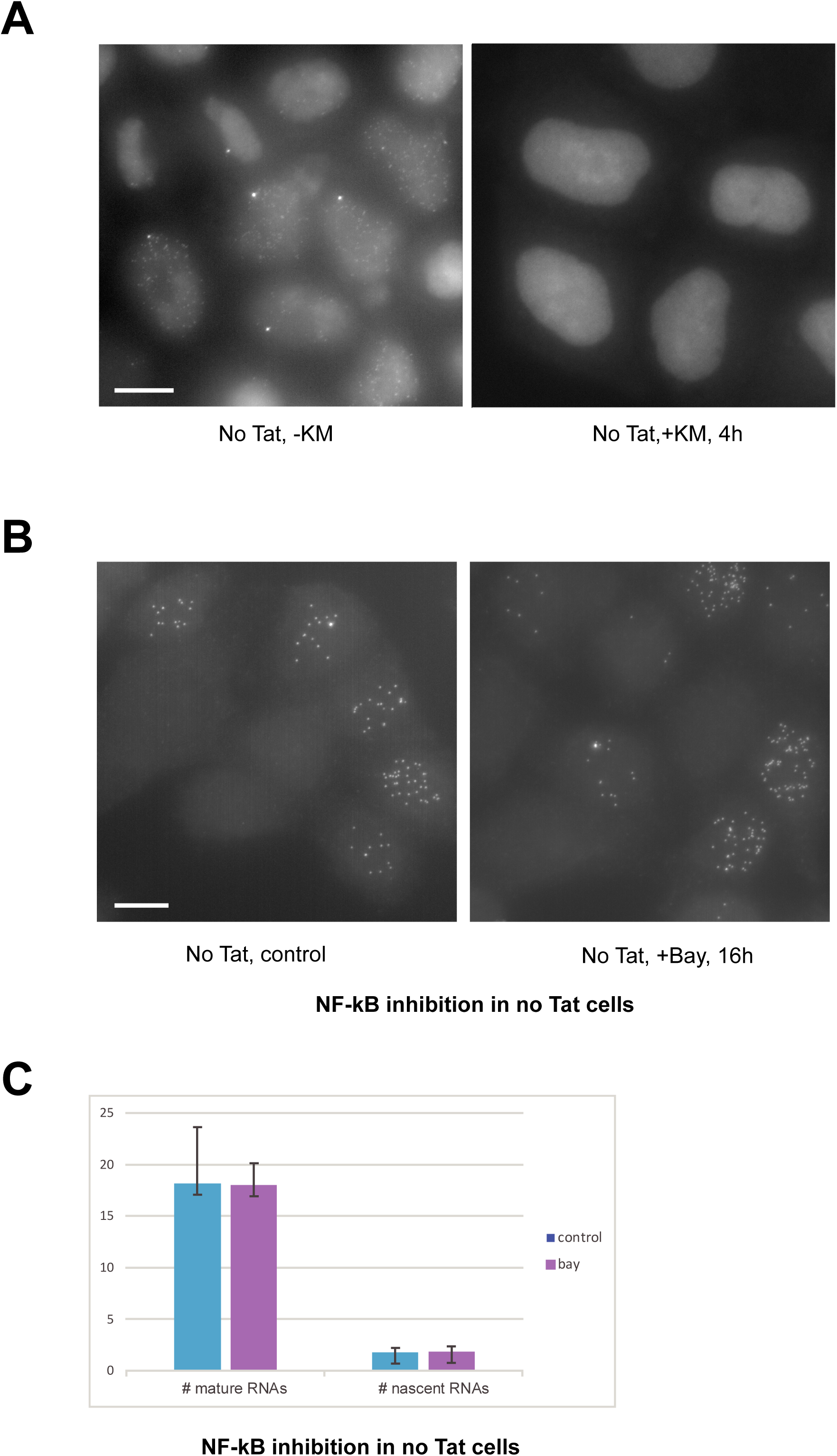
Transcriptional activation of HIV-1 reporter in the absence of Tat depends on enzymatic activity of CDK9 and is independent of NF-κB pathway. **A-** CDK9 inhibitor KM05283 inhibits HIV-1 transcription. Images of Hela Flp-in HIV-1 128xMS2 MCP-GFP no Tat cells treated with 100 μM KM05382 for 4 h, using GFP filter. Left – non-treated control; right – 4 hours of KM05382 treatment. The scale bar is 10 μM. **B-** NF-κB inhibitor BAY11-7082 does not affect HIV-1 reporter transcription. Left panel: Images of smFISH with Cy3 labeled probes of the cells Hela Flp-in HIV-1 128xMS2 MCP-GFP no Tat. Left - non-treated control; right -16h treatment with 2 μM BAY11-7082. **C-** Histogram showing the quantification of mature and nascent RNA number on the smFISH images after 16h inhibition of NF-kB with 2 μM BAY11-7082. On y axis is the RNA number. Error bars are standard deviations.

**Supplementary Figure S3.**
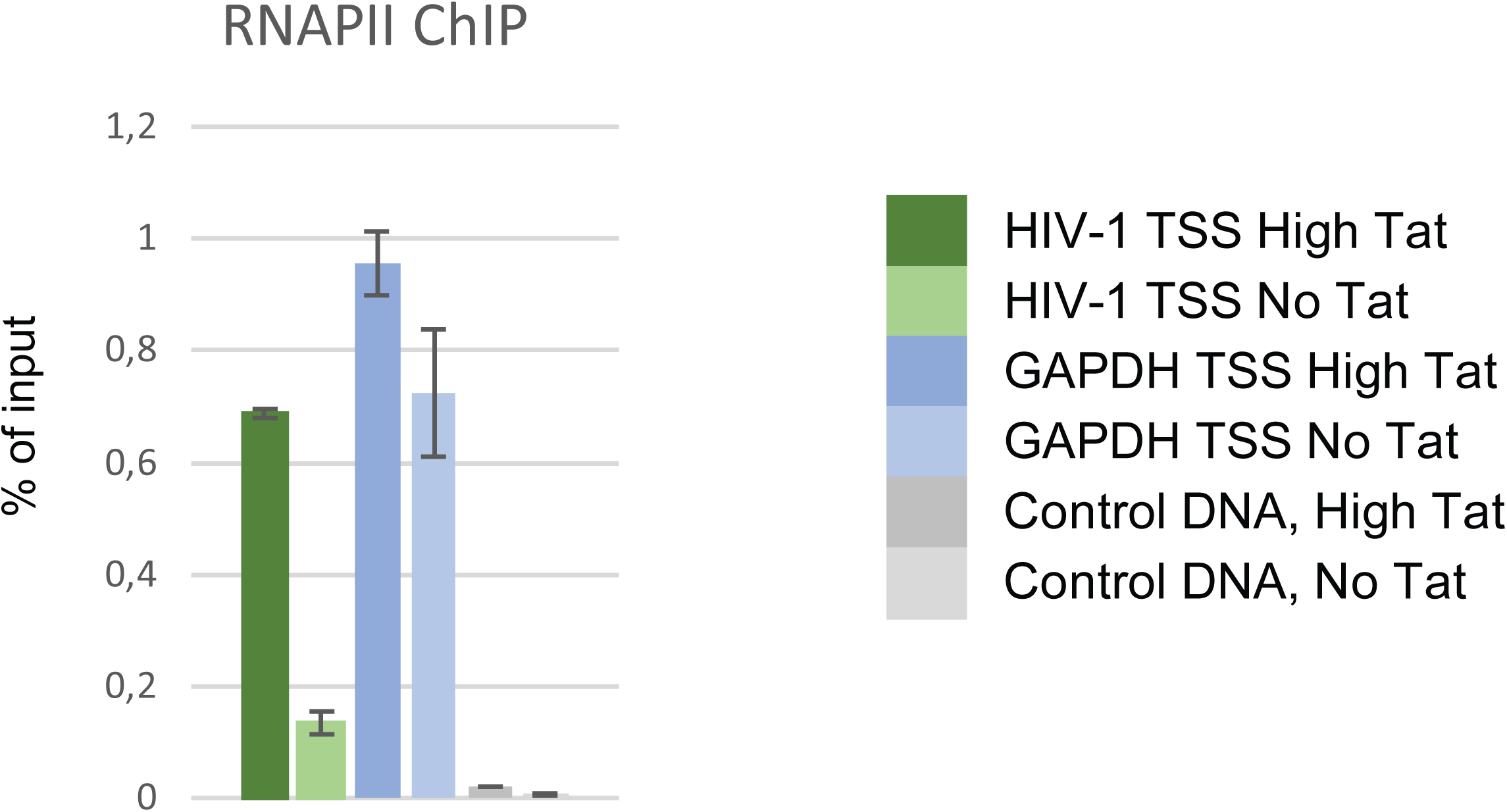
RNA Pol II ChIP in presence and absence of Tat. The graph depicts the RNA polymerase II ChIP signals at HIV-1 and GAPDH loci for the High Tat and No Tat cell lines. GAPDH TSS: transcription start site of the human GAPDH gene; HIV-1 TSS: transcription start site of the HIV-1 promoter; Control DNA: a non-transcribed genomic locus. ChIP signals were measure by qPCR and values are expressed as percent of input (y axis). The scale bar is 10 μM.

## Hybrid symbolic/numeric method for reverse engineering of transcriptional bursting processes

## Supplemental text to accompany

### 1 Introduction

#### 1.1 Summary of the method

We use machine learning to derive characteristics of single cell transcription activity from MS2 data. The output of the machine learning procedure is threefold. Using a **deconvolution method** and high resolution movies, we generate a **time map of transcription events** indicating, for each cell, the moments when different RNAP molecules start producing mRNA. This direct readout of transcriptional events in a cell population, represents a unique feature of our method, not available in other methods that fit directly a particular transcription model to the MS2 data such as methods based on the autocorrelation function [2, 5, 4], or maximal likelihood estimate [1] or on Bayesian inference [7, 6]. The map can be used for direct characterisation of transcription features, such as polymerase convoys and various statistics of inter-event times. A second output of the approach is a **multiscale cumulative distribution function** of the waiting time separating succesive transcription events (or the complementary function, called survival function). We provide both non-parametric Kaplan-Meyer and parametric multi-exponential estimates of the multiscale distribution function. This distribution, obtained by combining short, high-resolution and long, low-resolution movies, covers timescales from second to 10 hours. The dynamical range of our method supersedes those of other extant methods that are based on much smaller sampling rates and/or much shorter movie lengths. The waiting time distribution is model-free, but can be further used to identify various models of transcription dynamics. The third output of our method is the **model parameter identification**, simultaneously for several models that fit the data equally well. Although we focus on discrete transcription models based on Markovian transitions among hidden promoter states (for different number of states and for a rich collection of transition graph topologies), our method can be extended to the identification of more general models, including continuous or hybrid ones. Contrary to other methods that need separate fitting procedures for different models, in our method a single parametric fit of the multiscale waiting time distribution function is enough for identifying simultaneously a large collection of models that are all compatible with data and perform equally well. Another novelty with respect to other model fitting methods is the use of exact symbolic solutions, relating the parameters of the multiscale distributions to kinetic parameters of the model. For several models there is one-to-one relation between parameters of the distribution and kinetic parameters of the model. In this situation, the model kinetic parameters can be obtained analytically from the parameters of the multiscale distribution. Our method also leads to uncertainty estimates of the model parameters, based on optimal and close-to-optimal parametric fits of the multiscale distributions. As a matter of fact, models that fit equally well, can differ in their parametric uncertainties. Therefore, parametric uncertainty can be used as a model selection criterion that favor sure and reject uncertain models. The symbolic part of our method also identifies situations when parametric uncertainty results from redundancy, more precisely when there are manifolds of parameters that lead all to exactly the same goodness of fit. This is typically the situation when the relation between parameters of the multiscale distribution and the model parameters is one to many. Model and/or parameter uncertainty can be ultimately lifted by direct measurements of one or several kinetic parameters by alternative methods.

#### 1.2 Discrete Markovian models for transcription dynamics

A Markovian model of transcription dynamics includes stochastic transitions between several ON and OFF promoter states (Figure 1). Rather generally there is a ON state and several OFF states. The promoter transcribes only in the state ON when it can trigger several departures of RNAP molecules along DNA. The departure of one RNAP is when the model reaches the state EL. It is considered that immediately after departure the operator site becomes free (the transition from EL to ON is instantenous). The transitions define a continuous time Markov chain characterized by a set of positive parameters *k*_*ij*_ representing the transition probability per unit time (or equivalently the inverse mean transition time) from state *i* to state *j*. Given the number of states *N*, the structure of the Markov chain is defined by the directed graph *G* = {(*i, j*)|1 ≤ *i* ≤ *N*, 1 ≤ *j* ≤ *N, k*_*i,j*_ ≠ 0}; several possible structures with *N* = 3 are shown in Figure 1. We show here how the parameters *k*_*ij*_ of a model can be adjusted to reproduce the transcriptional bursting and RNA synthesis observed in the live cells experiments. The parameter estimates are performed simultaneously for several possible model structures.

**Figure 1:**
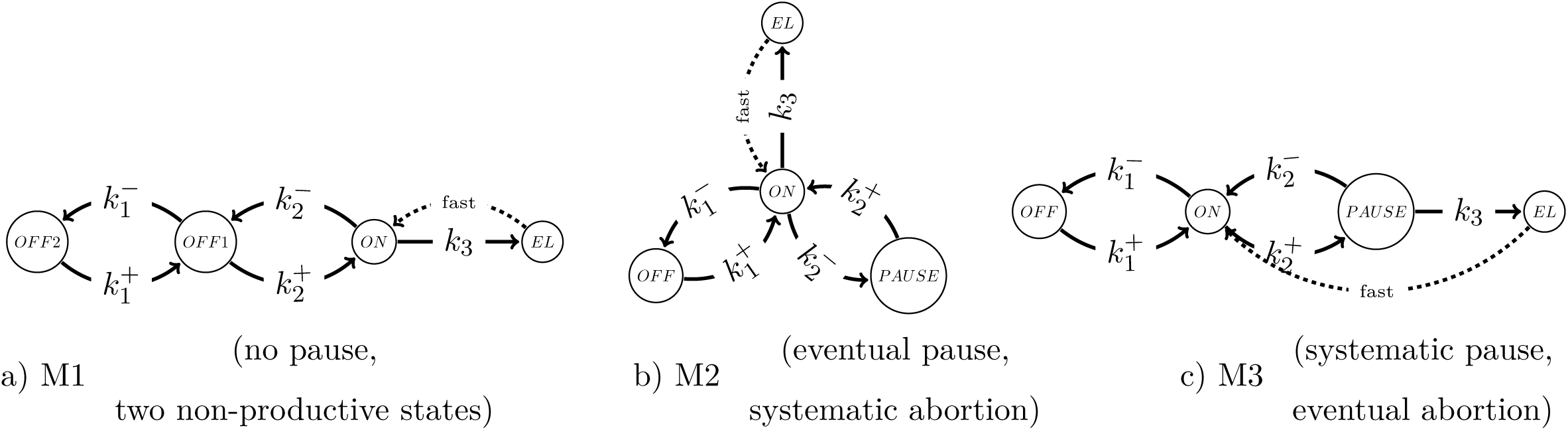
Three state, three exponential models of transcription dynamics. Transcription dynamics is represented as transitions between two OFF and one ON promoter states. ON represents the productive state. EL represents the elongation state which immediately liberates the promoter (from EL there is always fast return to ON). ON may not lead systematically to EL, for instance if there is transcription pausing. In these models, the pausing state is represented as one of the OFF state. If the pause leads systematically to transcription abortion the pause state leads to ON (model M2); otherwise it leads both to ON and to EL, with different probabilities (model M3). In model M1 the inactive state OFF1 can lead to another inactive state OFF2. The represented topologies differ by the connections between different states. The constants *k*_*i*_ are inverses of transition times, as such a) M1: *k*_3_ is the initiation rate; b) M2: *k*_3_ is the initiation rate, 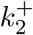 is the abortion rate, 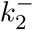 is the pause enter rate; c) M3: *k*_3_ is the pause exit rate and 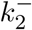 is the abortion rate. All the represented models have 3 states (excepting the final elongation state), 5 kinetic parameters and their stochastic transcription activity can be described by a three exponential survival function.

#### 1.3 The machine learning procedure

This procedure was initially designed for MS2 data obtained from human cell cultures but it has also been applied to in vivo study of Drosophila embryo development [3].

The machine learning procedure has several steps:

a. The first step is the **numerical deconvolution of the signal and Pol II Positioning**. The signal from each cell is a convolution between the contribution of a single polymerase and the point process (set of time points) describing all the start transcription events. For each cell, we reconstruct the start events by a least square optimization method performed by a genetic algorithm. We prefer optimization to Fourier transform based deconvolution in order to avoid the Gibbs phenomenon (the signal produced by a single polymerase is discontinuous).
b. The second step is the **non-parametric estimate of the survival function**. The data resulting at a) (positions of transcription start events) is used to estimate the survival function which is the complementary cumulative distribution function of the inter-event (waiting) times. Further complexity is brought in at this step by the utilization of two types of movies with short and long time resolution. Only short movies data undergoes deconvolution, the long movies are used to obtain long waiting times directly. This results into two distribution functions that are joined together (by affine transformations corresponding to the law of total probability) to cover many decades of timescales (second to ten hours). In certain applications the hours scale is not reachable because of biological constraints. For instance, in developmental biology, the studied developmental stage may be too short. In this case, we use only short movies and the joining step is not needed.
c. The third step is the **multi-exponential regression of the survival function**, performed by gradient optimization with random starting guesses, uniformly distributed in logarithmic scale (this choice is dictated by the multiscale nature of the signal). Usually, two or three exponentials (i.e. two or three time scales) are enough to describe our data. Chosing more than three exponentials is justified when this improves the fit without increasing parameter uncertainty. Conversely, choosing less exponentials is justified if this does not diminish the fit while decreasing parameter uncertainty. The first three steps of our procedure are model-free because they make no assumption about the dynamics of the transcription regulation.
d. The last step of the procedure is the **symbolic reverse engineering of transcription models** from the survival function. We consider that the transcription machinery has several discrete states among which only one is productive. Then, the waiting time between successive transcription start events is the first return time to the productive state. The distribution of this waiting time satisfies a system of ODEs whose solution can be expressed as a sum of exponentials. The inverse problem consists in computing model’s kinetic parameters from the parameters of the multi-exponential regression. We have developed a symbolic solution to perform this step. Our symbolic solution also tackles the ill-posed character of the inverse problem. Indeed, although the same distribution function can be produced by several models with different structures, the significance and the value of each parameter are different in different models. Moreover, we know precisely how to pass from one model to another by changing the parameter values. It is therefore enough to perform a direct independent experimental measurement of a single parameter in order to discriminate between different models. In the case of redundant parameters (parameters not influencing independently the observed distribution function) and parameter uncertainty, some parameters may remain independent and can be used for model discrimination.

### 2 Numerical deconvolution of short movies

#### 2.1 Description of the problem

The experimental data obtained from short movies is shown in the Figure 2 for the HIV-1 promoter. The signal intensity from the mRNA MS2 reporter is represented as a function of time for each active transcription site. We are interested in reconstructing from this signal the sequence of waiting times between successive transcription start events (see Figure 3), for each transcription site.

Transcription events can not be straightforwardly detected from local features of the intensity signal because at a given time and for the same transcription site, more than one polymerase transcribe simultaneously. Furthermore, the signal from one polymerase does not appear immediately after initiation (see below).

One should thus consider that experimental data is a a convolution between the sequence of start events {*t*_*i*_, 1 ≤ *i* ≤ *N*_*pol*_} and the signal *h*(*t*) from a polymerase molecule:

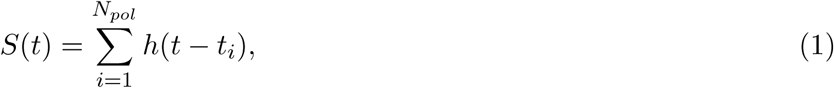

where *N*_*pol*_ is the number of polymerases contributing to the signal. *N*_*pol*_ is not known and will be determined by the optimization procedure (see below). The parameters *t*_*i*_ are the initial polymerase positions on the DNA, indicating the transcription start events.

The polymerase signal *h*(*t*) can be described as follows (see Figure 4):

i. Transcription begins when the RNA polymerase II leaves the promoter. However, no signal will be generated yet.
ii. The fluorescence signal is generated as soon as the polymerase reaches the MS2 sequence. During the transcription of the MS2 sequence the signal can be represented as a linear ramp-up.
iii. The signal will stay constant from the end of the MS2 sequence until when the polymerase leaves the transcription site, when the signal falls abruptly.

**Figure 2:**
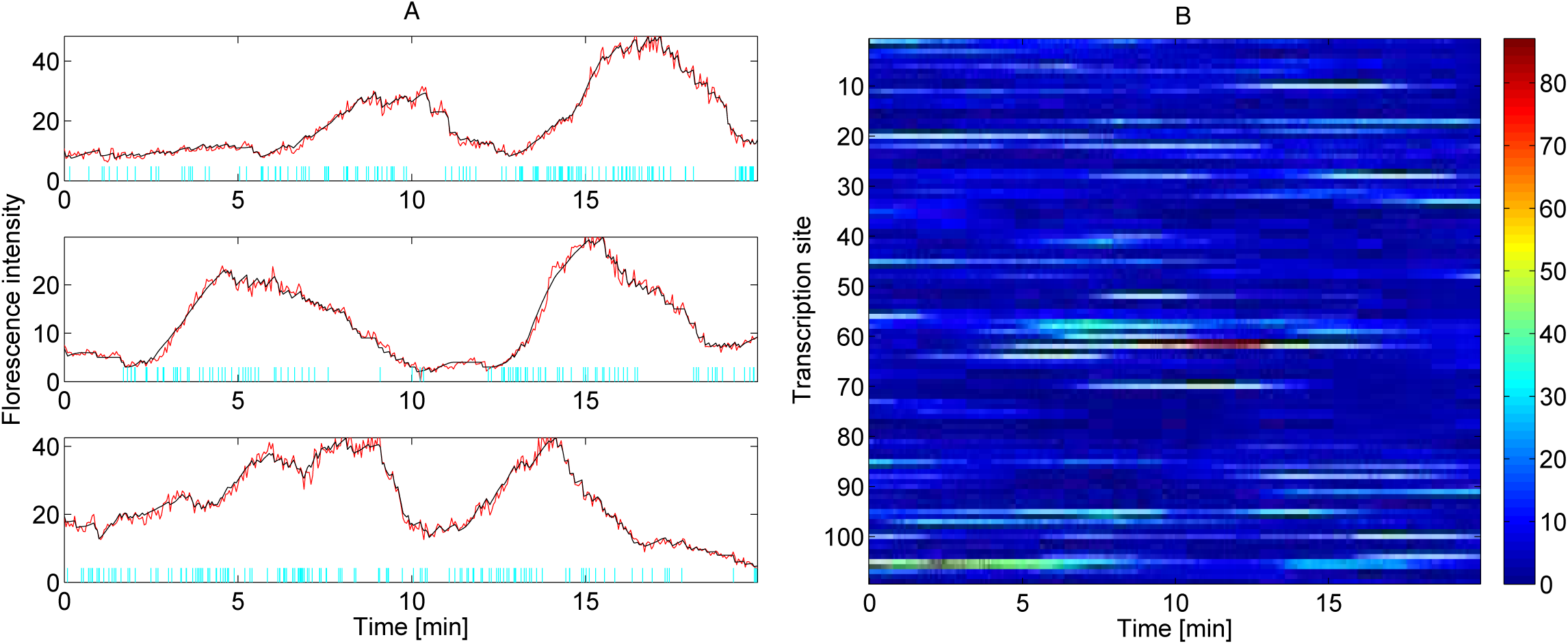
Short movie data for a HIV-1 promoter (no tat condition). A) fluorescence intensity vs. time for several transcription sites (red line); the reconstructed polymerase positions and signal are indicated as vertical cyan bars and black line, respectively. B) colormap of intensity for all transcription sites in a short movie.

**Figure 3:**
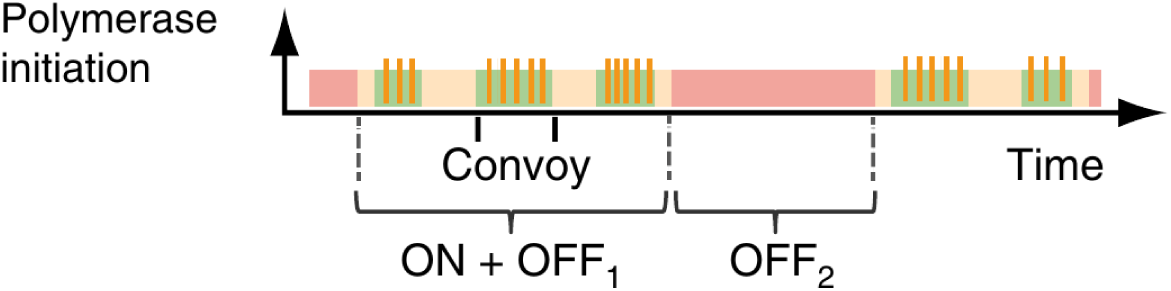
Dynamics of the promoter changing states. Start events are represented as red bars. During a ON period several polymerases start, forming a convoy. For this signal, two types of non-productive states, short (OFF1) and long (OFF2) can be obseved.

In order to compute the times corresponding to the three stages we use the length (expressed in base pairs) of the three sequences PRE, SEQ and POST (before MS2, MS2 and post MS2). These lengths depend on the MS2 construction (the values in our HIV-1 experiments are PRE=700bp, SEQ=2900bp, POST=1600bp). The sequence lengths are divided by the polymerase speed *V*_*pol*_ to be transformed into times. For our HIV-1 promoter we have used *V*_*pol*_ = 67*bp/s* (see main text). An extra time *P*_*poly*_ = 100*s* is added to POST, corresponding to the polyadenylation signal (during this time the polymerase has finished transcription and waits on the transcription site).

**Figure 4:**
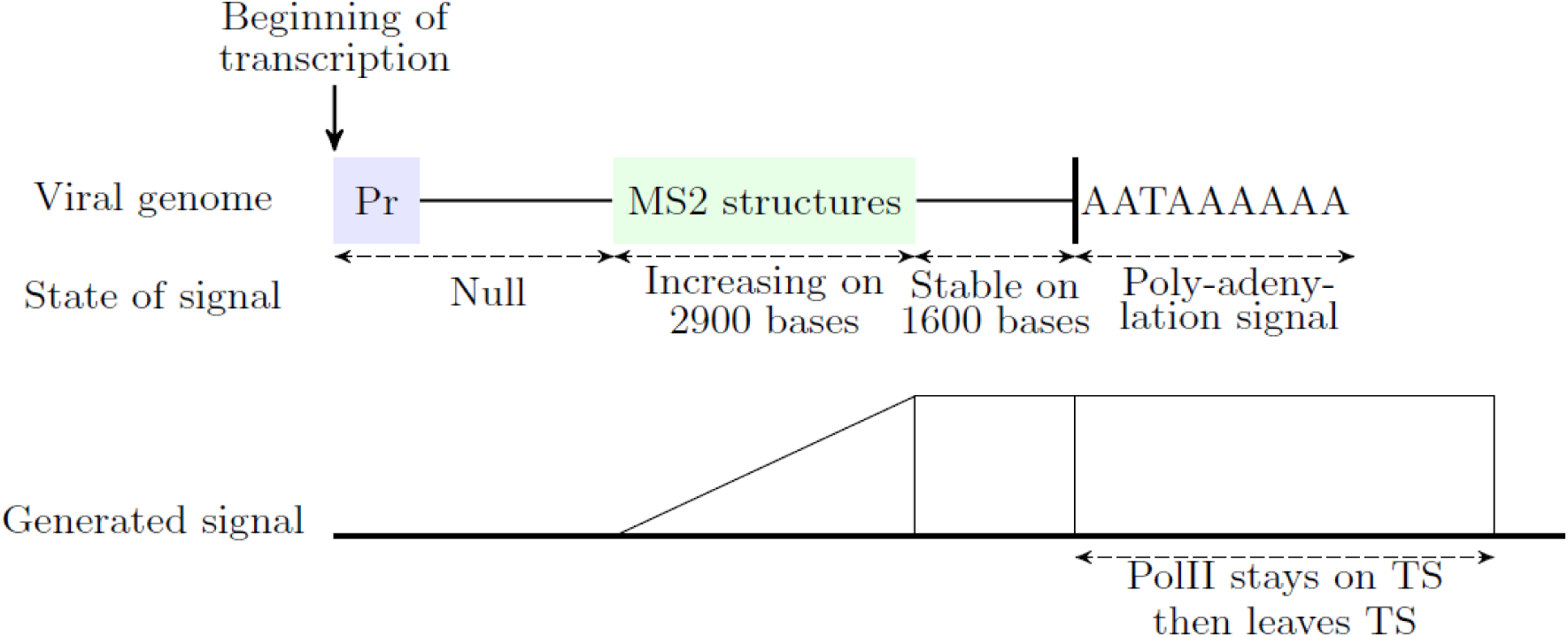
Representation of the signal from one polymerase for the HIV1-promoter. The parameters are indicative and can change for other applications.

**Figure 5:**
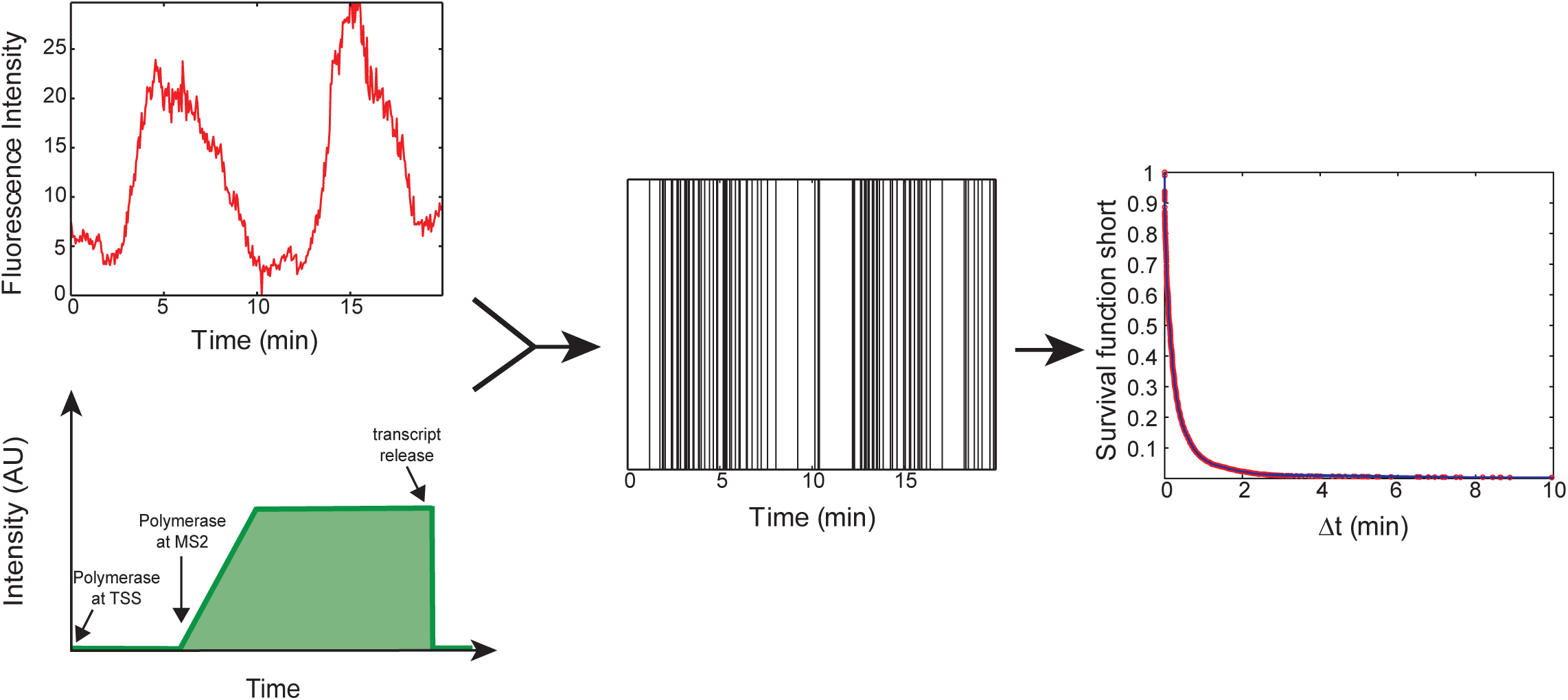
Distribution of transcription initiation events can be reconstructed by deconvolution.

**Figure 6:**
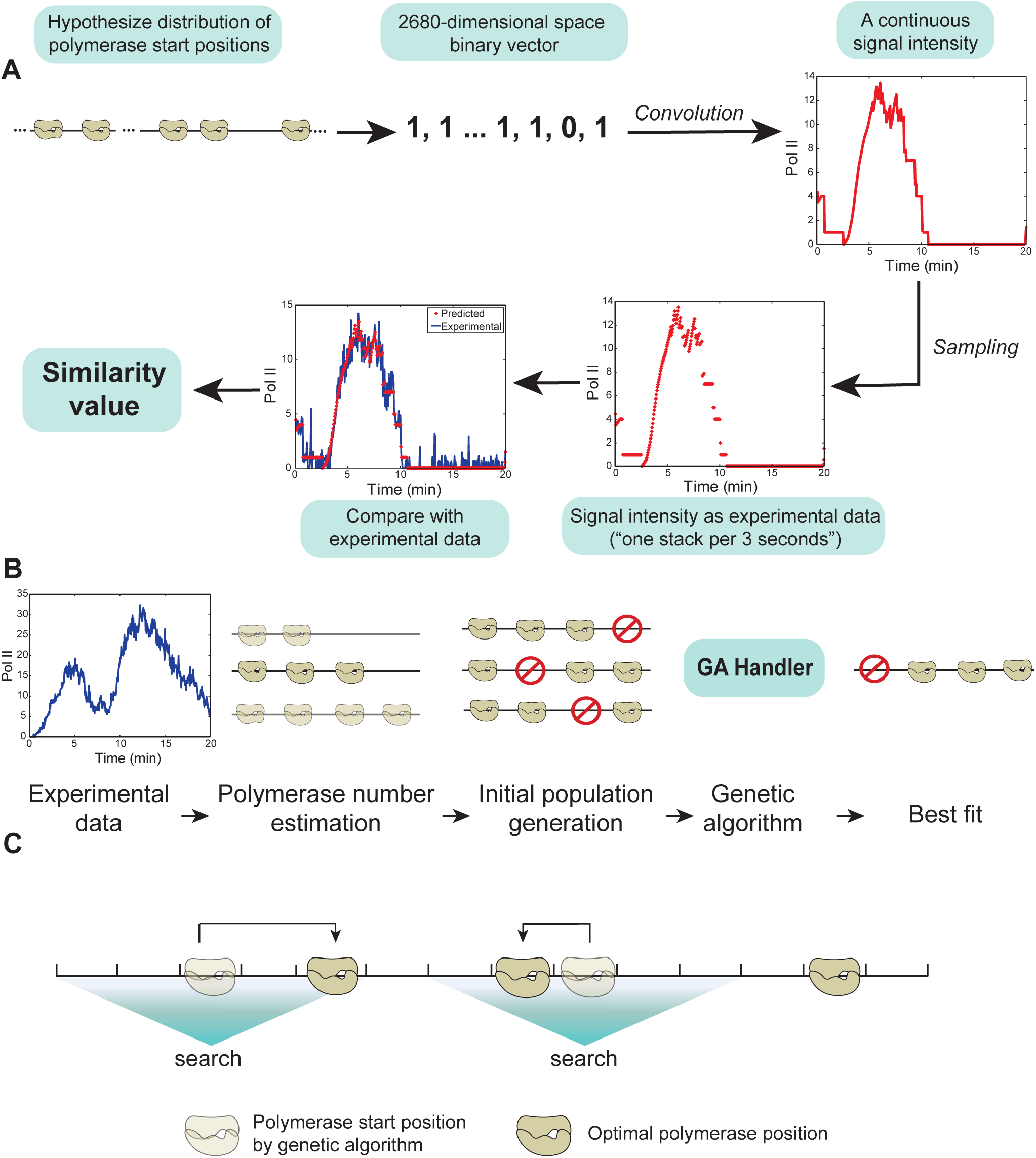
A. Flowchart of the numerical deconvolution method. B. Genetic algorithm step. C. Local optimization step.

After signal calibration the unit of fluorescence represents the amplitude of the signal from one polymerase. Using this model we want to reconstruct the sequence of initiation events by deconvolution (see Figure 5).

More precisely, we will determine *N*_*pol*_ and *t*_*i*_, 1 ≤ *i* ≤ *N*_*pol*_ that minimize the following objective function:

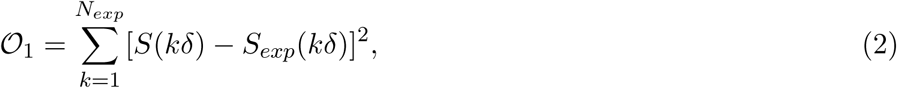

where *δ* is the time step (inverse frame rate), *N*_*exp*_ is the number of frames, *S* is described by (1) and *S*_*exp*_ is the experimental signal. For a short movie, the frame rate is 1*/*3*s*^−1^, thus *δ* = 3*s* and *N*_*exp*_ = 400 for a movie length of *P*_*max*_ = 20min.

#### 2.2 Discretization of the optimization problem

It is useful to use a dual representation of the polymerase positions *t*_*i*_ in terms of seconds and base pairs on the DNA sequence. Although the polymerase positions are in principle continuous variables, for computation reasons we discretize them. In this dual representation, it is natural to consider that possible polymerase positions are multiples of the minimum distance *d*_*min*_ between two polymerases (*d*_*min*_ = 30bp in our program). The precise value of *d*_*min*_ is not needed. Generally, *d*_*min*_ should be chosen as small as possible to guarantee precision of the polymerase positions. It should be smaller than the real minimum distance between polymerases and larger than a value dictated by the computation costs (the computation costs increase when *d*_*min*_ decreases).

Using this discretization, polymerase positions are coded as a binary vector. Every possible polymerase position will have either 1 or 0 value, which represents if there is a polymerase in the current position or not. Considering that the polymerase speed is constant, the polymerase positions are all we need to determine the signal. For a movie of length *P*_*max*_ = 20min and for *V*_*poly*_ = 67bp/s, the polymerase positions are represented as a binary vector of length *N* = *P*_*max*_*V*_*poly*_*/d*_*min*_ = 2680. Considering discretization steps larger than *d*_*min*_ is also possible. In this case binary vectors are shorter and computation is faster, however the precision may be reduced.

#### 2.3 Solve the deconvolution problem by a genetic algorithm followed by local optimization

Every binary vector of dimension *N* represent the polymerase start positions and determine a value of the objective function (2). The opposite of the objective function is the *fitness*. The deconvolution problem represents finding the minimum of this objective function (maximum of fitness) in the *N* dimensional binary space (Figure 6). In order to solve this hard combinatorial problem, we apply first a global optimization genetic algorithm (GA).

As shown in Figure 6 B, GA follows three steps: estimating the amount of polymerases, generating an initial population and applying genetic algorithm. We estimate the number of polymerases *N*_*pol*_ from the signal integral intensity, as the ratio of integral intensities of the experimental signal and of the single polymerase signal. The resulting amount is not an accurate number, and it is a rough estimation which can be used to accelerate next steps. Then we prepare an initial population according to the estimation of polymerase amount. Starting with a vector with *N* ‘0’s, we randomly pick *N*_*pol*_ positions and change them into ‘1’s. After the preparation of initial population, we use the genetic algorithm implemented in the GA solver provided by Matlab global optimization toolbox. Mutation, crossover and selection are processed by the MATLAB built-in function ga (MATLAB, version (R2013b), Natick, Massachusetts: The MathWorks Inc.). At each step, the genetic algorithm solver selects individuals at random from the current population to be parents and uses them to produce the children for the next generation. Over successive generations, the population “evolves” toward an optimal solution.

In order to verify this method, we implemented a test using an artificial experimental signal. We deconvolved the artificial signal, for which we know exactly the polymerase start positions. The simulation of genetic algorithm, as in the example of Figure 7, shows that the genetic algorithm can approximately reconstruct the signal. However, the global minimum is not precisely reached and the polymerase start positions of simulation are not exactly the same as the artificial ones (Figure 7).

There are various reasons why polymerases were not exactly placed into right positions as follows: the limitation of the maximum number of iterations, the limitation of population size, the initial error in the estimated number of polymerases, the noise generated by the algorithm, etc.. Although GA can not give a precise result (or it is time consuming to get a precise result), it provides a solution not far from the optimal result.

With this in mind, we use a local exhaustive search to accomplish local optimization. The idea is to “move” a polymerase left or right relative to the GA found position to see if this improves the fitness function (Figure 6 C). For every polymerase we find the best position which has the highest fitness value and we update the best positions for all of them. The local optimization result is shown in Figure 7. By this method, practically all the polymerases were arranged into the correct positions. The local optimisation method has limitations, for instance it does not allow correction of the total number of polymerases; we suppose that this number has already been found by the GA.

**Figure 7:**
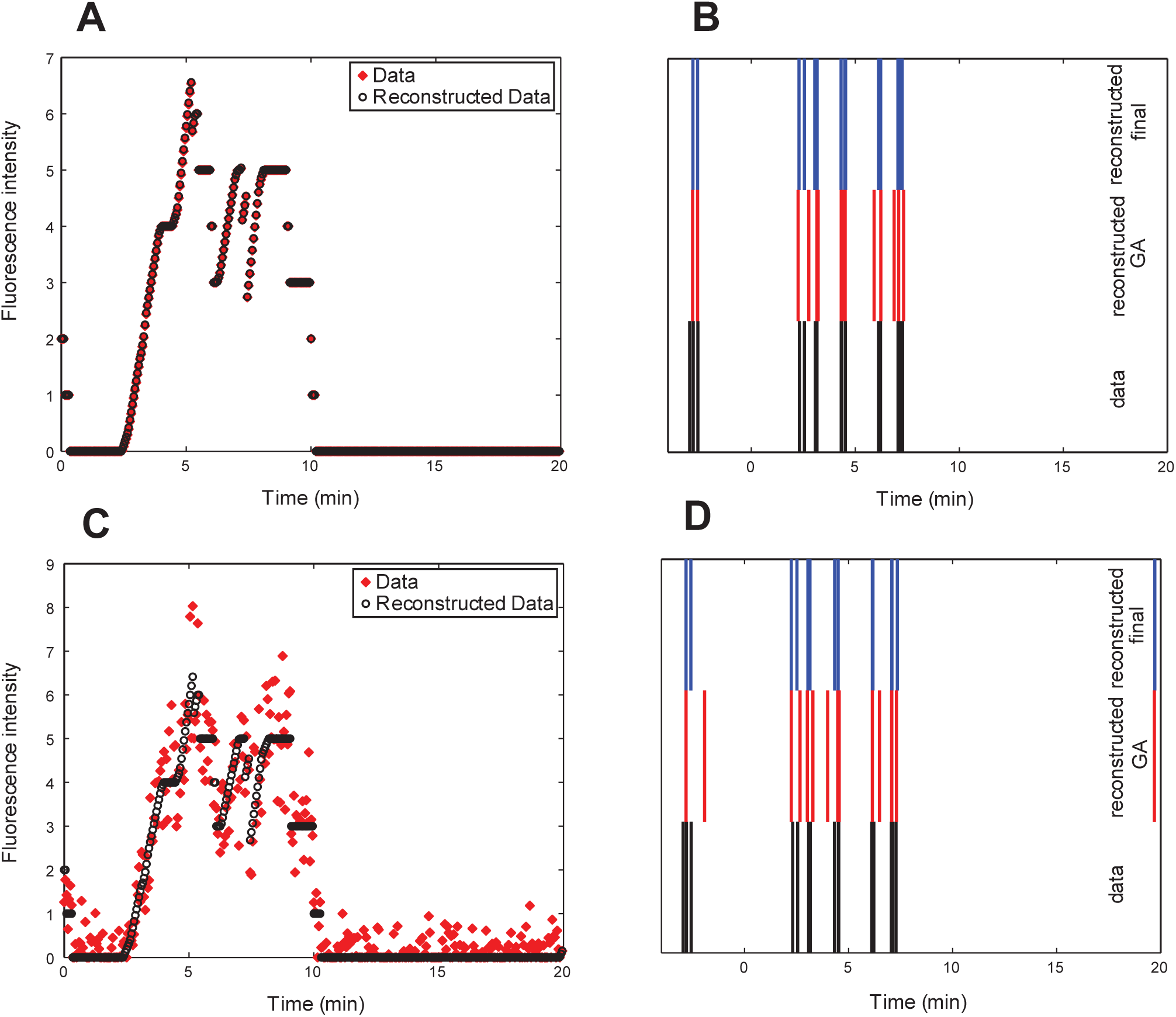
Result of the deconvolution step on artifical data. A) Signal generated artificially (no noise). B) Original polymerase positions compared to positions resulting from the genetic algorithm step and to the final positions corrected by local optimisation (no noise). C) Signal generated artificially (noise added according to the procedure described in Section 8). D) Original polymerase positions compared to positions resulting from the genetic algorithm step and to the final positions corrected by local optimisation (noise added).

### 3 Multi-exponential regression of the distribution function

From the numerical deconvolution step, we obtain the series of intial polymerase positions, for each active transcription site detected in the short movies.

From each transcription site we compute waiting times defined as time interval between successive positions. When the position is the last one in the movie, the waiting time is defined as the distance to the end of the movie. Considering that all transcription sites are statistically equivalent, we gather the waiting times from all sites that are active in the same short movie.

Long movies also provide waiting times by a different method, without numerical deconvolution (see below).

We consider that the transcription events form a renewal process with independent, identically distributed waiting times *Δ*. This property is valid at stationarity, but not only. For instance, in Markovian models, the property is true if after every transcription event the system allways returns to the same state. Given that non-Markovian models can be made Markovian by adding hidden states, we believe that the property is quite general. All the models from Figure 1 satisfy this property, because from the *EL* state one can only go to the *ON* state.

We want to estimate the *complementary cumulative distribution function* (also called *survival function* in survival analysis) of the waiting times defined as:

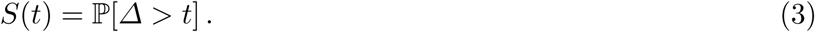

#### 3.1 Waiting times from short movies: *Δ*_*s*_

##### Outliers handling

Several observed transcription sites had abnormal behaviour (too many or too few events). The decision was made to take them off the data set as follows

1. Compute the amount of events of transcription that happened during the movie (Ev).
2. Compute the 1st and the 3rd quartile (respectively Q1 and Q3) of the distribution of Ev.
3. Only consider the transcription sites where

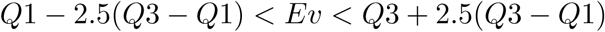

##### Affine transformation, parameter *p*_*s*_

The *Δ*_*s*_ are the waiting times deduced from the short movies, therefore they all satisfy the condition *Δ*_*s*_ < *P*_*max*_, where *P*_*max*_ is the movie length. Therefore, this data does not reconstruct the full survival function, but the conditional survival function *S*_<*P*_*max* (*t*) = ℙ[*Δ* > *t*|*Δ* < *P*_*max*_].

In order to compute the relation between *S*(*t*) and *S*_<*P*_*max* (*t*) we use the total probability theorem:

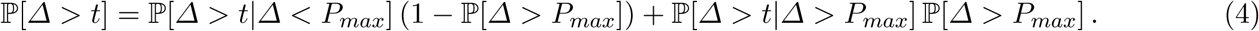

Let us note that for *t* < *P*_*max*_, one has ℙ[*Δ* > *t*|*Δ* > *P*_*max*_] = 1. Hence, from (4) it follows that

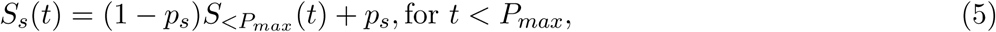

where *p*_*s*_ = ℙ[*Δ* > *P*_*max*_] is the probability that the waiting time is longer than the length of the short movie.

In other words, for short movies, the survival function is obtained from the conditional survival function by an affine transformation.

#### 3.2 Waiting times from long movies: *Δ*_*l*_

##### Active and inactive periods, threshold parameter

The frame rate for long movies is 1*/*3*min*^−1^ and the typical length is 9*h*. Let us notice that the deconvolution procedure is not possible for long movies, because the number of polymerases is too large. Therefore, long waiting times are obtained directly from the signal. For long movies, there is no need to calibrate the fluorescence intensity, nor to deconvolve the signal. An intensity threshold is defined and a given transcription site is considered active in a given frame if its intensity is larger than the threshold, inactive if not, see Figure 8.

##### Outliers handling

We define the fraction of inactivity (FI) as the ratio of cumulative total inactivity time to the total cumulative time in the long movie and for all the transcription sites.

Some transcription sites in long movies data set also show unusual behaviours being active (FI=0) or inactive (FI=1) during the entire movie. We exclude these outliers as we did it for the short movies, but based on the fraction of inactivity for each transcription site.

We will only consider the transcription sites from the long movies where

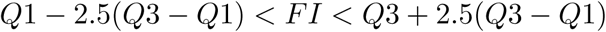

##### Corrected waiting times, parameter *Δ*_0_

In the long movies the waiting times *Δ*_*l*_ between successive transcription initiations correspond roughly to the inactive periods *Δ*_*I*_. As a matter of fact, these waiting times can be longer than the inactive periods by a time varying between 0 and 6min (because the signal needs about 3min to vanish and starts about 3min before it is detected). This unknown time is a parameter of the method and its values are discretized to *Δ*_0_ = 0, 3, 6. All waiting times are computed as *Δ*_*l*_ = *Δ*_*I*_ + *Δ*_0_.

**Figure 8:**
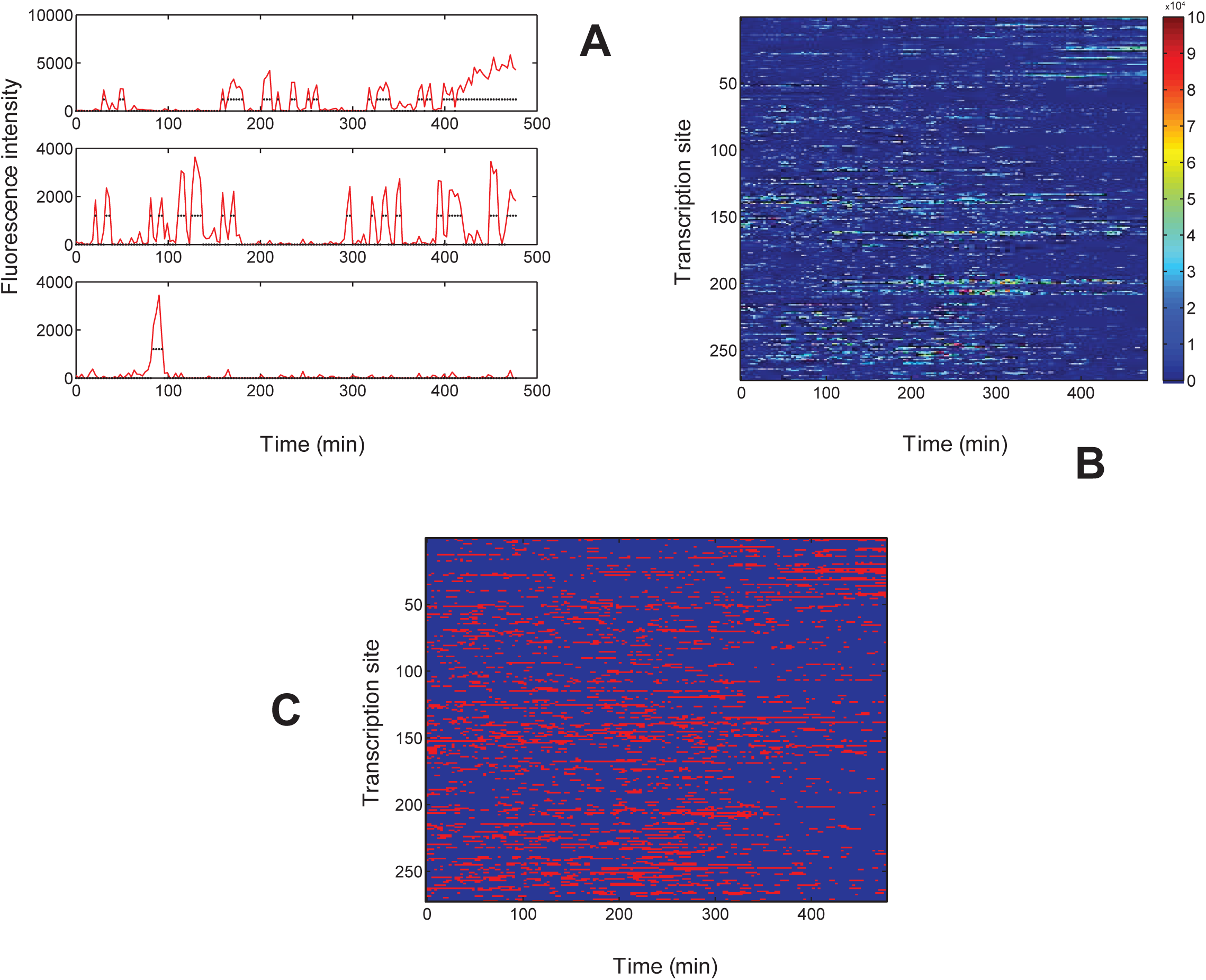
Long movie data for a HIV-1 promoter (no tat condition). A) intensity of fluorescence vs. time for several transcription sites; the real valued intensity are transformed into a binary signal (dots) by thresholding (here the threshold value is 1200). B) intensities for all transcription sites. C) Binary valued intensities for all transcription sites.

##### Affine transformation, parameter *p*_*l*_

The signal from a single polymerase lasts roughly 3min (see Figure 4). In this case, waiting times shorter than *P*_*min*_ where *P*_*min*_ is roughly 3min can not be observed. The observed conditional survival function is now 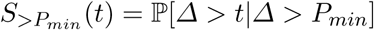.

We can note that for *t* > *P*_*min*_, one has ℙ[*Δ* < *t*|*Δ* < *P*_*min*_] = 1.

Once again, from the total probability theorem (4), it follows:

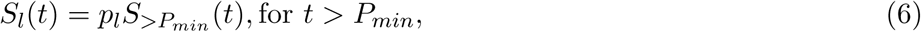

where *p*_*l*_ = ℙ[*Δ* > *P*_*min*_] is the probability that the waiting time is longer than *P*_*min*_, *S*_*l*_ is the survival function for long movies.

##### Estimate of parameter *p*_*l*_

The probability *p*_*l*_ is estimated by combining information extracted from the long and the short movies. *p*_*l*_ is precisely the probability that waiting times are observed as inactive periods in the long movie. Let *N*_*inactive*_ and *N*_*active*_ be the number of waiting times observed as inactive periods, and hidden within active periods of the long movie, respectively. *N*_*inactive*_ can be determined directly from the long movie, it represents the number of inactive periods. *N*_*active*_ is obtained as the ratio *P*_*active*_*/*𝔼[*Δ*|*Δ* < *P*_*min*_] where *P*_*active*_ is the cumulative time of all active periods in the long movie and 𝔼[*Δ*|*Δ* < *P*_*min*_] is the conditional expectancy of the waiting time provided that this is smaller than *P*_*min*_ therefore undetectable by the long movie. By definition one has

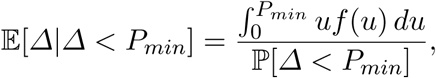

where *f* is the probability density function of *Δ*. Taking the derivative of 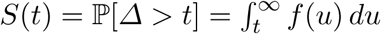 we get *f*(*t*) = −*S*′(*t*). Using the integral by parts formula we find

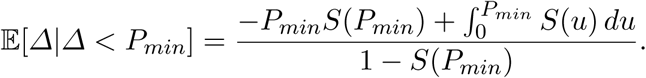

Summarizing, we find

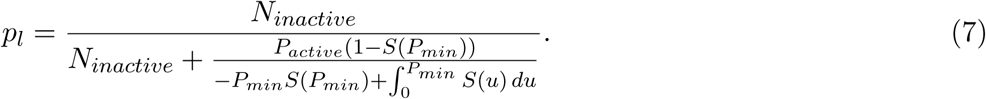

Both *S*(*t*) and the integral above are computed using the survival function of the short movie. *N*_*inactive*_ and *P*_*active*_ are determined from the long movie data.

##### Estimate of parameter *p*_*s*_

*p*_*s*_ is estimated by optimization. We look for the value of *p*_*s*_ that minimizes the square distance between the solutions (5) and (6) on the overlap interval [*P*_*min*_, *P*_*max*_].

The survival functions after calculation of *p*_*l*_, *p*_*s*_ and affine transformations are shown in Figure 9.

**Figure 9:**
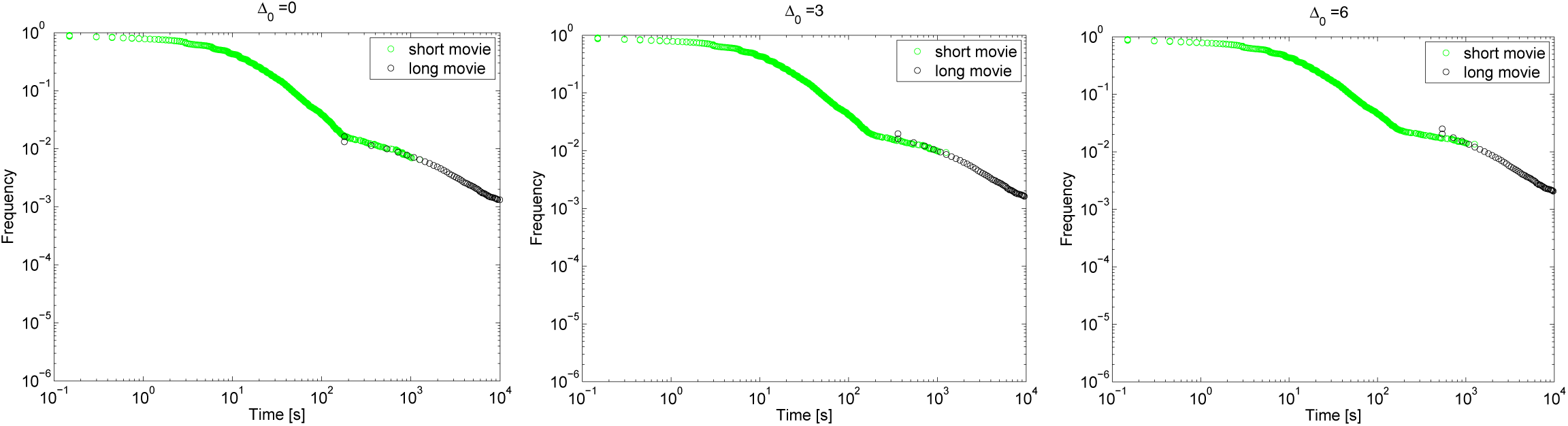
Survival functions of the waiting time after affine transformations for several values of the shift *Δ*_0_ (HIV promoter, no tat condition, see text).

**Figure 10:**
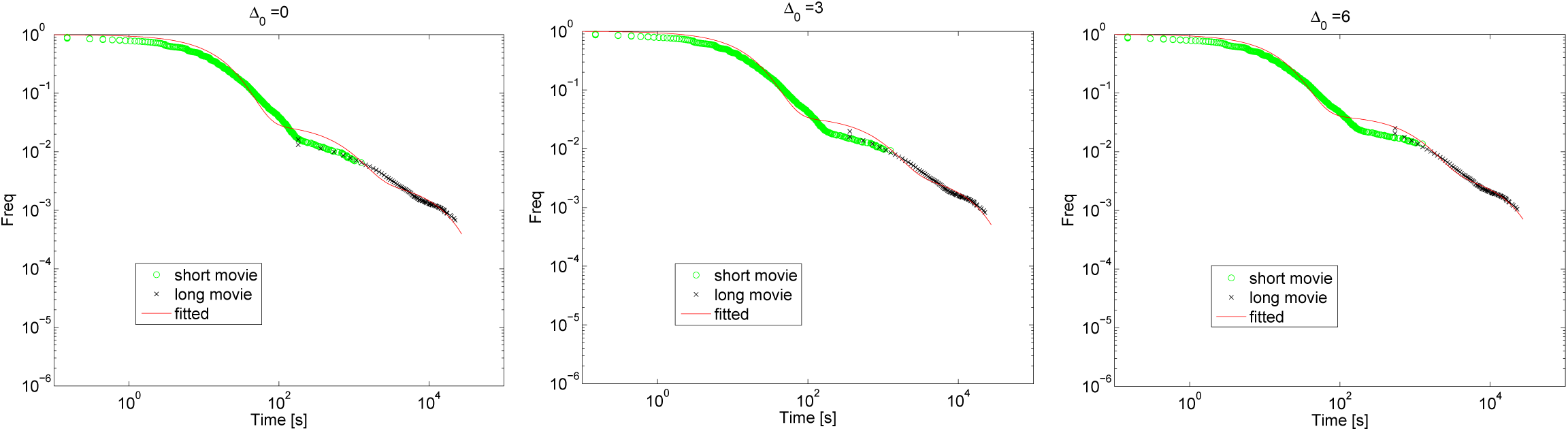
Multi-exponential regression (*N* = 3) for several values of the shift *Δ*_0_ (HIV promoter, no tat condition, see text).

#### 3.3 Multiexponential regression

The previously determined survival function function (5), (6) is modelled by a multiexponential function:

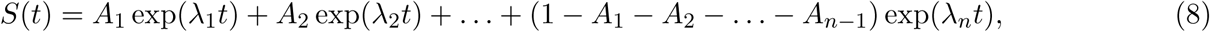

where *A*_1_, …, *A*_*n*−1_, *λ*_1_, …, *λ*_*n*_ are 2*n* − 1 parameters.

Because lim_*t*→∞_ *S*(*t*) = 0, these parameters must satisfy the constraints *λ*_*i*_ < 0, 1 ≤ *i* ≤ *n*. For practical reasons we can consider that all *λ*_*i*_ are distinct. Degenerate cases, when two or more *λ*_*i*_ are equal can be uniformly approximated by formula (8) with distinct *λ*_*i*_ (see the Section 4.2). Up to relabelling we can consider that |*λ*_1_| > |*λ*_2_| > … |*λ*_*n*_|. Furthermore, because the complementary distribution function is always decreasing we have

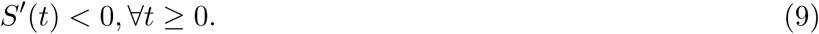

The condition (76) implies that 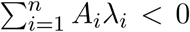, where 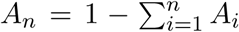 (follows from *S*′(0) < 0) and that *A*_*n*_ > 0 (this follows from lim_*t*→∞_ *S*′(*t*)*exp*(−*λ*_*n*_*t*) = *A*_*n*_*λ*_*n*_ < 0 and *λ*_*n*_ < 0). The hyperplanes 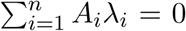, *A*_*n*_ = 0 together with other manifolds delineate the domain of valid parameters *A*_*i*_. This domain depends on the exponents *λ*_*i*_ as illustrated for *n* = 3 in Figure 11.

**Figure 11:**
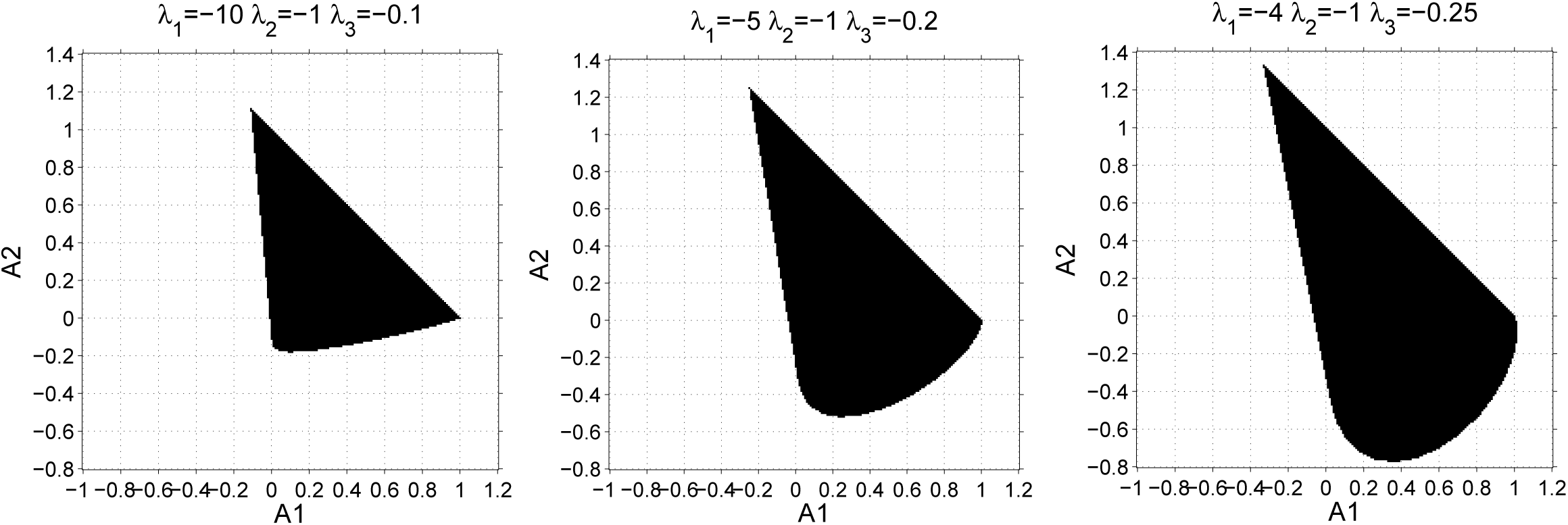
Permitted values of *A*_*i*_, 1 ≤ *i* ≤ *n* for *n* = 3 are represented in black for various *λ*_*i*_. These parameters values are defined by the condition 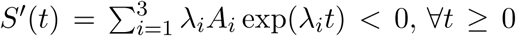, where *A*_3_ = 1 − *A*_1_ − *A*_2_. The permitted values are limited at the top right by the line *A*_1_ + *A*_2_ = 1 and at the left by the line *λ*_1_*A*_1_ + *λ*_2_*A*_2_ + *λ*_3_(1 − *A*_1_ − *A*_2_) = 0.

Like usually in machine learning, the choice of *n* is guided by a parcimony principle. One can start with *n* = 2 and progressively increase *n* until the goodness of fit stops improving (at equal goodness of fit, one favors the model with lowest complexity, lowest *n* and/or with lowest parameter uncertainty).

The objective function is defined as follows:

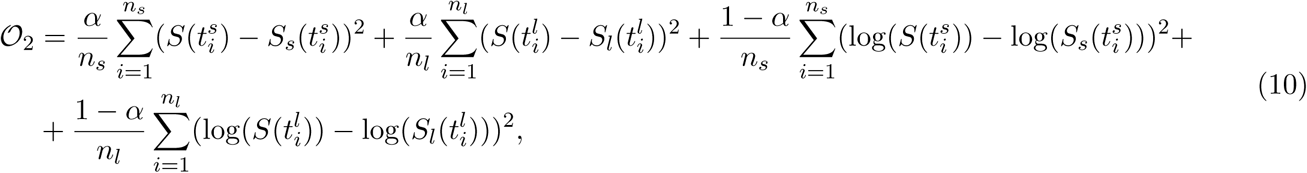

where *S*(*t*) is defined by (8); *S*_*s*_ and *S*_*l*_ are computed by (5),(6), respectively; 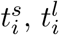 are sampling times for short and long movies, respectively; *α* is a positive weight representing the relative importance of the linear scale compared to the logarithmic scale in the representation of the survival function.

We minimize (10) by local optimization (Levenberg-Marquardt algorithm implemented in the Matlab function *lsqnonlin*) starting with *N*_*p*_ (in our program *N*_*p*_ = 100) random values of the regression parameters *A*_1_, …, *A*_*n*−1_, *λ*_1_, …, *λ*_*n*_. The initial parameters *A*_1_, …, *A*_*n*−1_ are chosen uniformly distributed in the cube [−*M, M*]^*n*−1^ (we used *M* = 2), whereas the initial parameters *λ*_1_, …, *λ*_*n*_ are all negative and log-uniformly distributed in absolute value. More precisely, log(|*λ*_*i*_|) are uniform in a cube (*l*_1_, *l*_2_, …, *l*_*n*_) + [−*K, K*]^*n*^, where *l*_1_ < *l*_2_ < … < *l*_*n*_.

The optimization is repeated for all values of *Δ*_0_ and each time repeated *N*_*p*_ times with different initial parameters (Figure8). We keep the lowest value 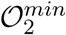 of (10) as well as sub-optimal solutions with 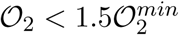. The suboptimal parameters are utilized to estimate the parameter uncertainty. For each parameter we compute an uncertainty interval defined by the minimum and the maximum values over the set of all optimal and suboptimal parameters. Uncertain parameters have large uncertainty intervals.

An example of multi-exponential fit is given in Figure 12.

**Figure 12:**
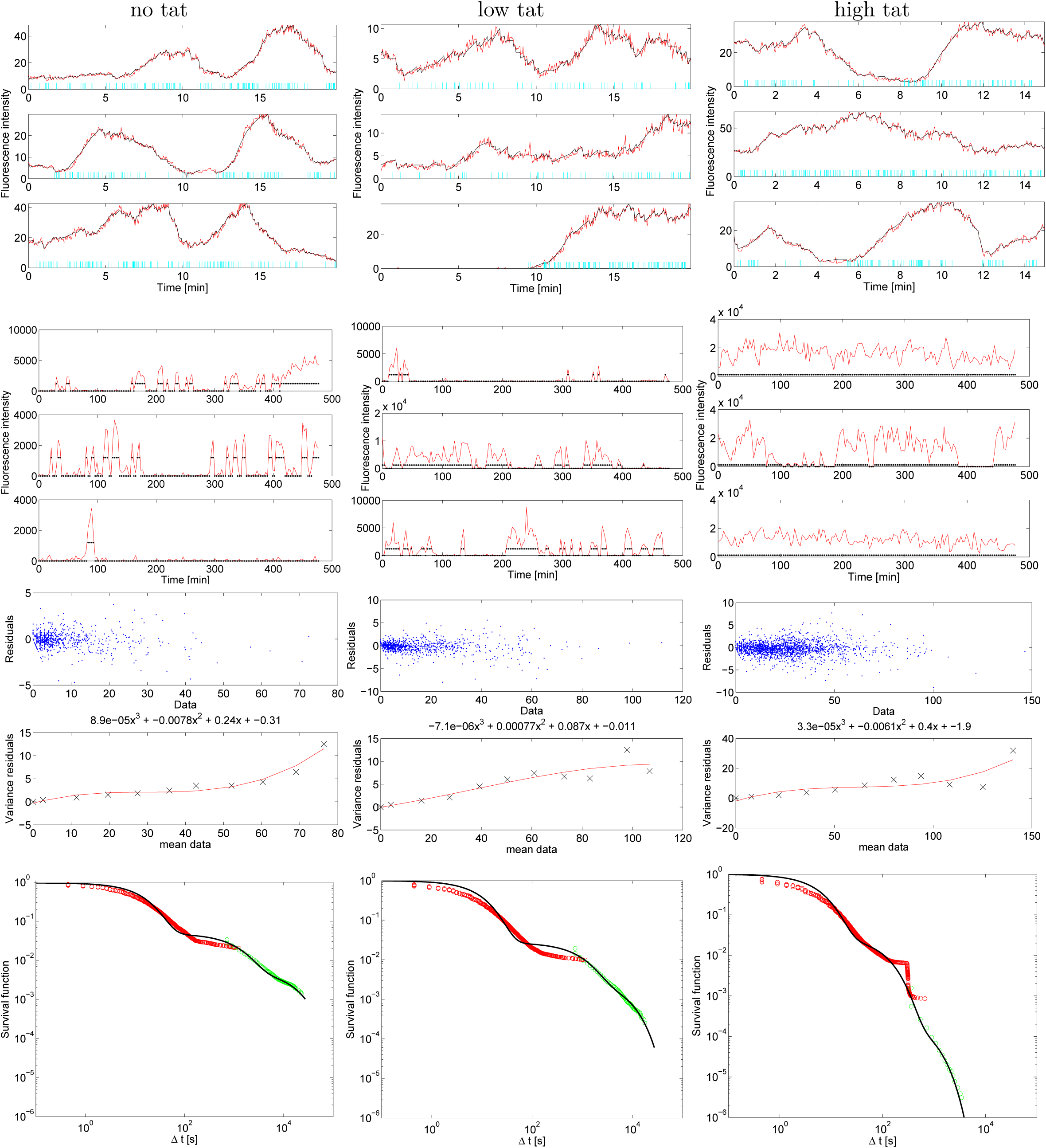
Results of the unconstrained three-exponential fit. First row: short movie data with reconstructed polymerase positions. Second row: long movie data. Third row: noise interpolation. Fourth row: most optimal fit for *α* = 0.30.

### 4 Symbolic solution to the inverse problem

#### 4.1 General model and waiting time distribution

We consider a continuous time Markov chain promoter model with N states *P*_*i*_, *i* ∈ [1, *N*]. One of these *N* states, *P*_*o*_, is the “ON” state from which polymerase can start transcription, and all the other states are “OFF” states (non-processive). A supplementary state *P*_*N*+1_, designates the start of processive elongation. From *P*_*N*+1_, there is systematic return to *P*_*o*_. The models have parameters *k*_*i,j*_, 1 ≤ *i, j* ≤ *N* + 1 indicating the transition rates from the promoter state *i* to the promoter state *j*. We consider that processive elongation immediately frees the operator and the promoter returns to the “ON” state. In mathematical terms

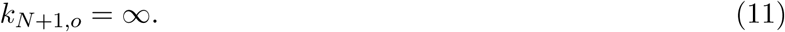

We also consider that only one state, denoted *X*_*N*_, can lead to processive elongation *X*_*N*+1_: *k*_*N,N*+1_ ≠ 0, *k*_*i,N*+1_ = 0, for 1 ≤ *i* ≤ *N* − 1. *X*_*N*_ is not necessarily *X*_*o*_, for instance it can be a paused transcription state.

Because the movies are always started when transcription sites are in the active state and supposing that after each transcription initiation there is return to the active state, we model the experimental waiting time as the first time when the promoter reaches the state *P*_*N*+1_ starting from *P*_*o*_. This is a first hitting time (or first passage time) problem. Because the lifetime of the state *P*_*N*+1_ is zero and *P*_*N*+1_ is always followed by *P*_*o*_ (see (11)), the same waiting time is also the first return time to *P*_*o*_.

In order to compute the distribution of the first hitting time we use the following standard method.

Let *M* (*t*) be the state of the Markov chain at the time *t*. For the purposes of this calculation, we can consider that *M* (*t*) stops when it reaches *P*_*N*+1_. Let *X*_*i*_ = ℙ[*M* (*t*) = *P*_*i*_|*M* (0) = *P*_*o*_]. Because *M* (*t*) is stopped in *P*_*N*+1_, one has *X*_*N*+1_ = ℙ[*M* (*t*) = *P*_*N*+1_|*M* (0) = *P*_*o*_] = ℙ[*Δ* ≤ *t*]. Thus, *X*_*N*+1_ is the cumulative distribution function of the waiting time *Δ* to reach *P*_*N*+1_ from *P*_*o*_. The survival function of *Δ* is *S*(*t*) = 1 − *X*_*N*+1_(*t*).

The variables *X*_*i*_(*t*), 1 ≤ *i* ≤ *N* + 1, satisfy the following system of linear differential equations (the master equation):

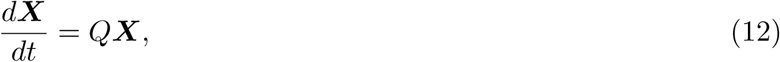

with the initial conditions *X*_*i*_(0) = *δ*_*i,o*_, where *δ* is the Kronecker symbol; *Q* is the transpose transition rate matrix whose elements are defined by *Q*_*j,i*_ = *k*_*i,j*_, *Q*_*i,i*_ = − _*j*…*i*_ *k*_*i,j*_.

Because *M* (*t*) is stopped in *P*_*N*+1_, the last column of the matrix *Q* is zero, namely *Q*_*i,N*+1_ = 0.

Let 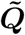 the *N* × *N* matrix obtained by eliminating the last line and the last column of the (*N* + 1) × (*N* + 1) matrix ***Q***.

Then 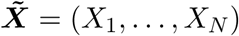 is the solution of the reduced equation

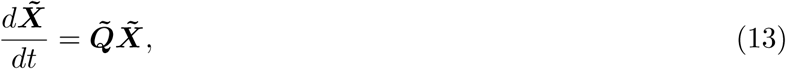

with initial conditions *X*_*i*_ = *δ*_*i,o*_ and reads

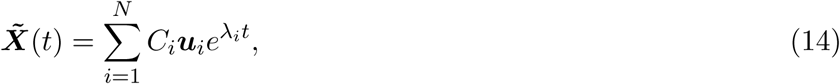

where *λ*_*i*_ and ***u***_*i*_, *i* ∈ [1, *N*] are eigenvalues and eigenvectors of 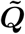, respectively.

Although (14) is written with the non-degenerate case in mind, when *λ*_1_ ≠ *λ*_2_ ≠ … ≠ *λ*_*N*_ (in general (14) is valid when 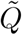 is diagonalizable), our final results, relating parameters of the survival function and kinetic parameters, can be extended to the degenerate case by continuous extension (see Section 4.2).

Furthermore, *X*_*N*+1_ can be obtained from the equation

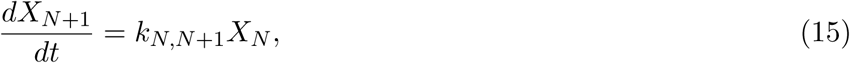

with the initial condition *X*_*N*+1_(0) = 0.

Without restricting generality, all eigenvectors ***u***_*i*_ can be chosen such that their o-th coordinate is 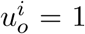.

Therefore, from (14), it follows

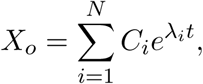

and from (15) it follows

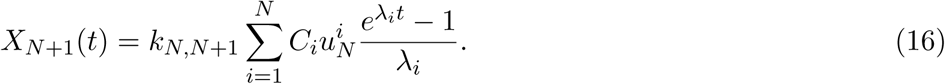

From lim_*t*→∞_ *X* (*t*) = 1 and (16) we get 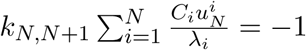. Using again (16) we find

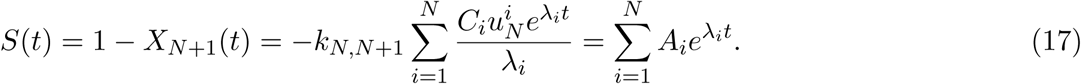

Hence

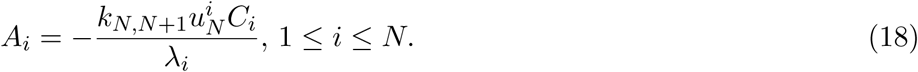

In particular, when *N* = *o*

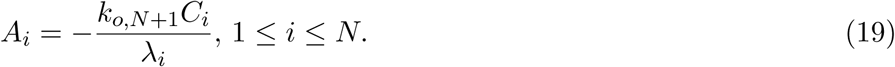

Eq(17) implies that in the non-degenerate case the survival function is a combination of exponential functions, implying that the waiting time has a mixed exponential distribution. However, although the mixture coefficients satisfy 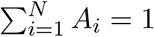, they are not guaranteed positive in general (see Figure 11).

#### 4.2 Degenerate case

The matrix 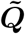 is a linear function in the model parameters *k*_*ij*_. According to classical results (see Kato), the eigenvalues of this matrix are branches of analytic functions in the parameters with only algebraic singularities. Moreover, the number of distinct eigenvalues is constant with the exception of a zero measure set of parameter values where this number is different. Excluding the permanently degenerate case when the matrix 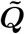 has a number of distinct eigenvalues smaller than *N* almost everywhere, we may consider that 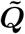 has *N* distinct eigenvalues except in a finite number of parameter values where it is degenerate, i.e. where *λ*_*i*_ = *λ*_*j*_ for at least two distinct indices *i* ≠ *j*.

Each degenerate case is arbitrarily close in the parameter space to a non-degenerate case. The solutions of the linear differential system (12) are continuous in the transition rates parameters *k*_*i,j*_, therefore the survival function computed in a degenerate case can be approximated by survival functions computed for non-degenerate cases. Because all the survival functions are monotone, by Dini’s theorem, this approximation can be made uniform for *t* ∈ [0, *T*], for any *T*. Using the inequality 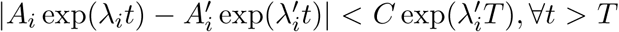, where 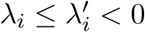, we can show that the uniform approximation is valid for all times.

Let us now compute the survival function in the degenerate case.

When, in spite of having degenerate eigenvalues (there are *N* independent eignevectors), the matrix 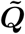 is diagonalizable, then Eqs.(14),(16),(17) hold. Therefore, in the diagonalizable case with degenerate eignevalues the survival function is a sum of less than *N* exponentials.

When the matrix 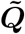 is not diagonalizable (there are less than *N* independent eigenvectors), (14) no longer holds.

Let *g*_*i*_ ≤ *n*_*i*_ be the geometric multiplicity (number of independent eigenvectors) of the eigenvalue *λ*_*i*_. Here *n*_*i*_ is the algebraic multiplicity of the eigenvalue *λ*_*i*_, representing the number of times this eigenvalue occurs as a root of the characteristic polynomial 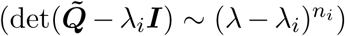 and one has 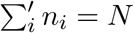, where the sum is over all distinct eigenvalues. Let us consider that *g*_*i*_ < *n*_*i*_ for at least one *i*. In this situation, 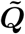 is not diagonalizable but can be reduced to a Jordan normal form. For each eigenvalue, there are *g*_*i*_ Jordan blocks. After reindexing the eigenvalues and Jordan blocks we have 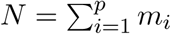, where *p* is the total number of Jordan blocks 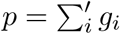 (the sum is over distinct *λ*_*i*_) and *m*_*i*_ is the dimension of a block *i*.

Let us remind that a generalized eigenvector ***v*** is any vector from the kernel 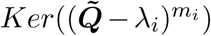. The subspace 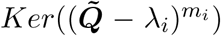 corresponds to a Jordan block and is generated by a chain of generalized eigenvectors 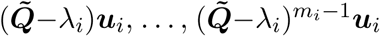, where ***u***_*i*_ is a generalized vector that satisfies 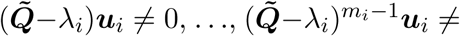. Furthermore, the solution of (13) starting from any generalized vector ***v*** reads:

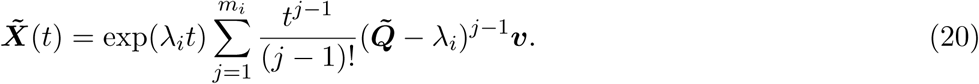

Let us consider that

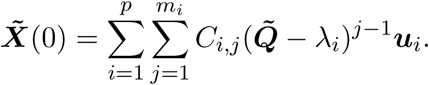

By the definition of the generalized eignvectors, 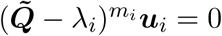.

Therefore, in the non-diagonalizable case, (14) must be replaced by:

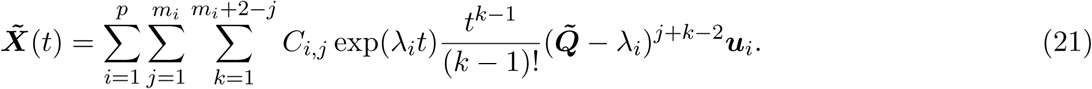

Then, (16) should be replaced by

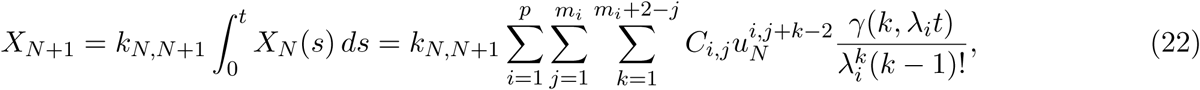

where 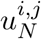 is the *N* ^*th*^ coordinate of 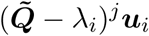, and *γ* is the incomplete gamma functions. It follows that (17) should be replaced by

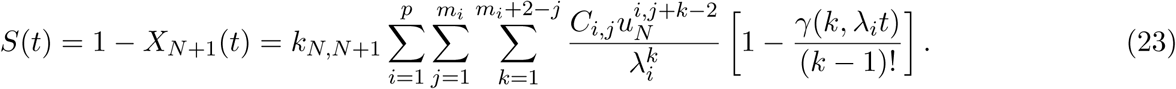

Eq. (23) implies that in the non-diagonalizable case, the survival function is a combination of gamma functions, implying that the waiting time has a mixed gamma distribution.

As an example illustrating this case let us consider the irreversible chain 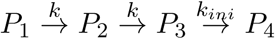 where *P*_3_ is the *ON* state and *P*_4_ is the *EL* state. in this case we have

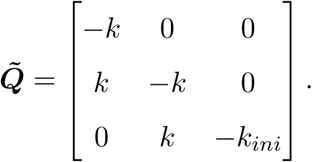

There are two distinct eigenvalues *λ*_1_ = −*k* and *λ*_2_ = −*k*_*ini*_. Each eigenvalue contributes with one Jordan block of dimensions 2 and 1, respectively. The chains of generalized eigenvectors are

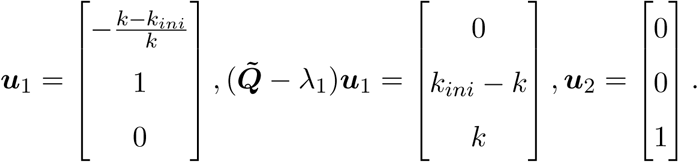

Suppose we want to compute the distribution of the waiting time to reach *P*_4_ starting from *P*_1_. Then

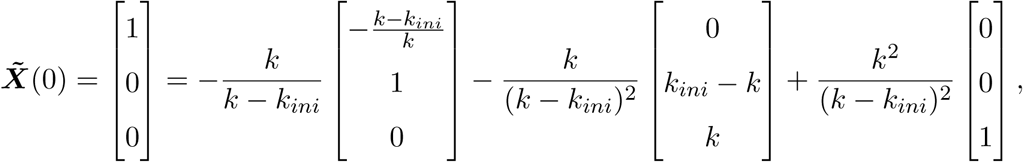

and

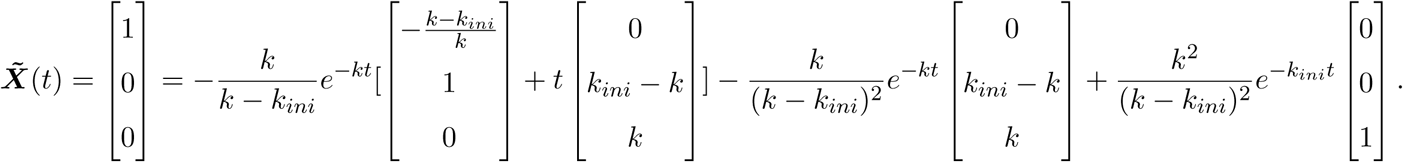

The survival function reads

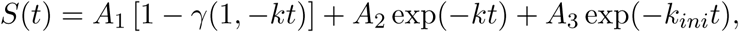

where *A*_1_ = −*k*_*ini*_*/*(*k* − *k*_*ini*_), *A*_2_ = −*k*_*ini*_*k/*(*k* − *k*_*ini*_)^2^, *A*_3_ = *k*^2^*/*(*k* − *k*_*ini*_)^2^ satisfy *A*_1_ + *A*_2_ + *A*_3_ = 1.

The waiting time is distributed according to a mixture of gamma and exponential distributions. If *k*_*ini*_ ≫*k*, then *A*_1_ ≈ 1, *A*_2_, *A*_3_ ≈ 0, meaning that the waiting time is distributed according to a gamma distribution of shape parameter 2 and scale parameter 1*/k*.

If in the previous model we make *k*_*ini*_ = *k*,

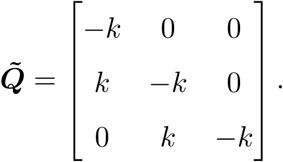

Then, 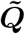 has only one eigenvalue *λ* = −*k* and one Jordan block of dimension 3. The chain of generalized eigenvectors is

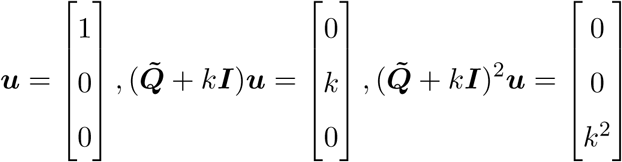

The survival function reads

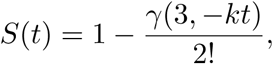

meaning that the waiting time is distributed according to a gamma distribution with scale parameter 1*/k* and shape parameter 3. This result is obvious from the structure of the model. If *k* = *k*_*ini*_, in order to reach *P*_4_ from *P*_1_ one needs three exponentially distributed steps of equal mean time 1*/k*; a sum of three independent equally distributed exponential variables is a gamma distribution of shape parameter 3.

In general, if the chain contains *n* limiting steps of constant *k*, the waiting time to reach the end of the chain starting from the beginning is distributed approximately according to a gamma distribution with shape parameter *n* and scale parameter 1*/k*.

#### 4.3 Inverse problem

In order to formulate a well posed inverse problem, we have to choose a structure of the model. The structure is defined by the directed graph *G* whose vertices are the promoter states and such that there is an edge from *i* to *j* if and only if *k*_*i,j*_ ≠ 0. Thus the model structure specifies which transitions are allowed between the promoter states. We also need to specify which one of the promoter states is ON.

Given a model structure, the inverse problem consists in computing the kinetic constants *k*_*i,j*_, 1 ≤ *i, j* ≤ *N* and *k*_*N,N*+1_ from the 2*N* −1 parameters of the survival function. This is possible only if there are at most 2*N* −1, kinetic constants. Uniqueness and thus well-posedness of the solution is possible only if there are exactly 2*N* − 1 parameters. However, not all models with 2*N* − 1 parameters have unique solutions of the inverse problem (an example is the model M3, see Figure 1 and Section 4.6).

In order to solve the inverse problem, we must write down the equations relating the parameters *k*_*i,j*_, *A*_*i*_ and *λ*_*i*_.

Let us consider that all the nonzero kinetic parameters are the 2*N* − 1 elements of a vector ***k*** ∈ ℝ^2*N* −1^.

##### Vieta’s formulas

Let us introduce the elementary symmetric polynomials of eigenvalues

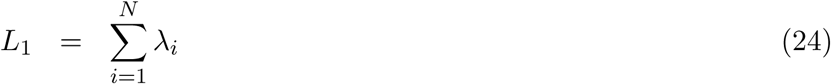

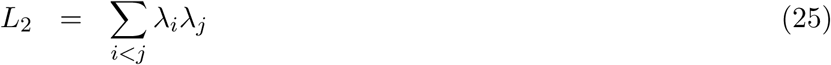

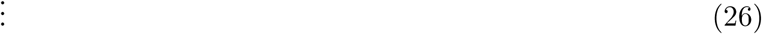

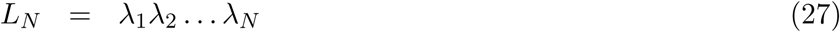

The characteristic polynomial of 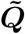 is

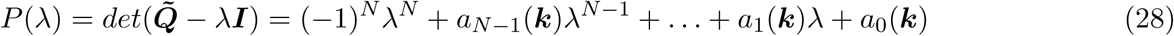

where the coefficients *a*_*i*_ are multivariate polynomial functions of the kinetic constants.

The coefficients of the characteristic polynomial are related to the symmetric polynomials of eigenvalues by the so-called Vieta’s formulas. We have the following *N* equations for the kinetic constants:

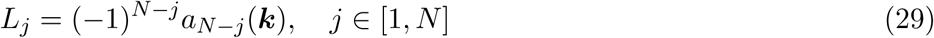

##### Eigenvectors

The eigenvectors of 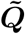 are solutions of the system of linear equations 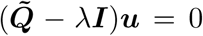 − *λ****I***)***u*** = 0 and are chosen of the form ***u*** = (*u*_1_(*λ*, ***k***), …, *u*_*o*−1_(*λ*, ***k***), 1, *u*_*o*+1_(*λ*, ***k***), …, *u*_*N*_ (*λ*, ***k***)), where *u*_*n*_(*λ*, ***k***), *n* ∈ [1, *N* − 1] are rational functions (ratios of polynomials) of *λ* and ***k***.

The initial conditions satisfied by the variables *X*_*i*_ provide a linear system of equations for the constants *C*_*i*_:

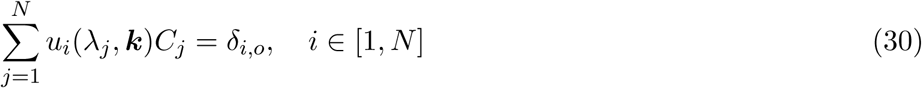

Let *C*_*i*_(***λ, k***), *i* ∈ [1, *N*] be the unique solution of (30).

From (18),(19) we get *N* − 1 equations for the kinetic constants ***k***:

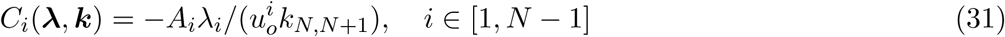

##### Inverse problem

The solution of the inverse problem is the solution of the system of 2*N* − 1 equations (29) and (31). In the next sections we solve this system symbolically. When a solution of the inverse problem exists, the kinetic parameters *k*_*i,j*_ can be expressed as functions in *λ*_*i*_ and *A*_*i*_. These functions are symmetric in the pairs (*λ*_*i*_, *A*_*i*_) and homogeneous of degree −1 in *λ*_*i*_. These functions are not always rational. For instance, they can have branching singularities, allowing, eventually, to pass from one solution to another, equivalent one. In general, multiple solutions are equivalent with respect to symmetries of the model. For instance, the model M2 in the Figure 1 is symmetric with respect to the permutation of the two lateral chains. In this case there are two solutions of the inverse problem, one solution being obtained from the other by permuting the parameters 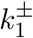 with 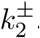. The general solutions will be presented elsewhere. In the sequel we provide full solutions for some models with *N* ≤ 4.

##### Recursion relations for eigenvectors

The eigenvector components *u*_*i*_(*λ*_*j*_, ***k***) can be obtained by recursion along the structure digraph.

We consider models such that any state of the promoter is connected to the ON state, in both directions, by directed paths on the structure digraph.

In the sequel, we discuss two representative cases.

The type I (*single chain*) model is a reversible chain ending with the *P*_*N*_ state:

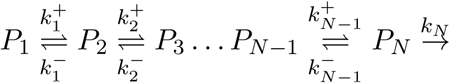

For this model, an eigenvector (*b*_1_, *b*_2_, …, *b*_*N*_) satisfies the equations

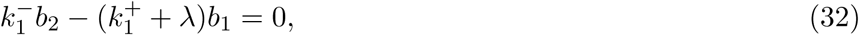

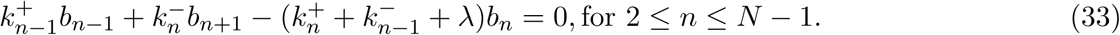

We can choose *b*_1_ = 1 and then from (32) 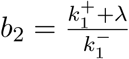. Threfore, *b*_*n*_ satisfy the recursion

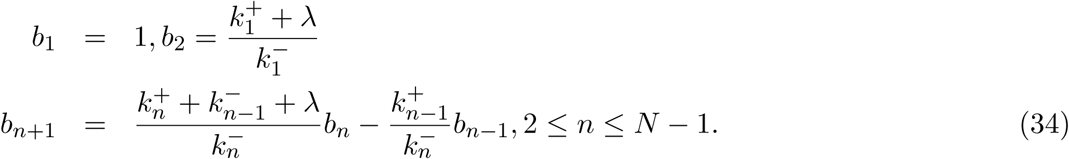

In order to have 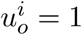 for all 1 ≤ *i* ≤ *N*, we define

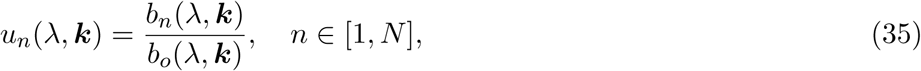

where *b*_*n*_ are rational functions of *λ* and ***k*** computed with the recursion (34).

The type II model is a reversible chain with the *P*_*N*_ state inside the chain:

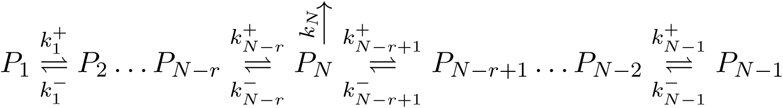

The type II model can be also described as two reversible chains branching from the *P*_*N*_ state. We can call this type *two chain model*.

For this model, an eigenvector (*b*_1_, *b*_2_, …, *b*_*N*_) satisfies the recursion

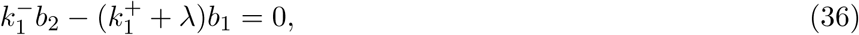

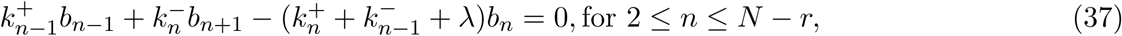

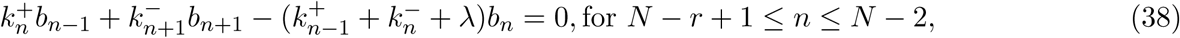

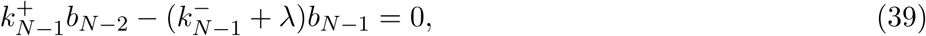

The recursion (36),(37),(38),(39) can be solved in the following way:

i. Choose *b*_1_ = 1 and compute *b*_2_ from (36).
ii. Use (37) to compute *b*_*n*_, 3 ≤ *n* ≤ *N* − *r* and *b*_*N*_.
iii. Use (39) and (38) to compute *b*_*n*_, *N* − *r* + 1 ≤ *n* ≤ *N* − 1 and *b*_*N*_ as multiples of *b*_*N*−1_.
iv. Determine *b*_*N*−1_ from *b*_*N*_, already computed at step ii).

Below we study several examples of type I and type II models.

#### 4.4 Symbolic solution to the inverse problem for the M1 model (*N* = 3)

This model is described by the transitions 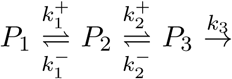. It is a type I model. In this case *P*_3_ is the ON state. The matrix of kinetic rates reads

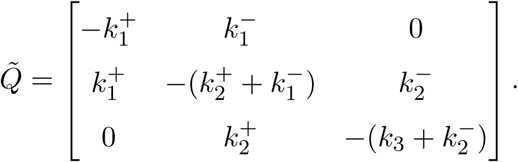

The characteristic polynomial of 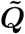 is 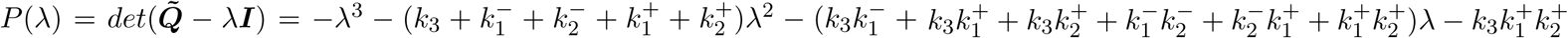.

The Vieta formulas read

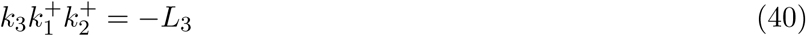

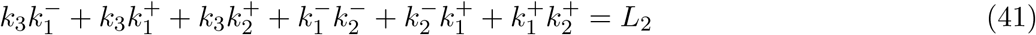

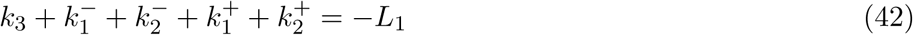

The solution of the recursion (34) is

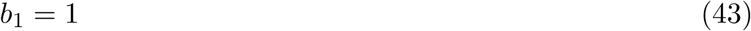

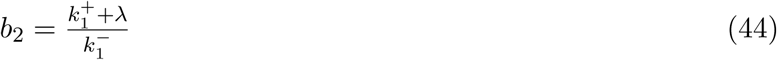

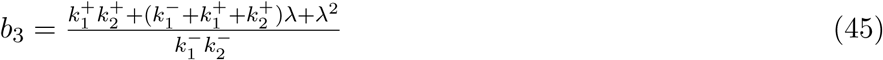

The system (30) has the solution

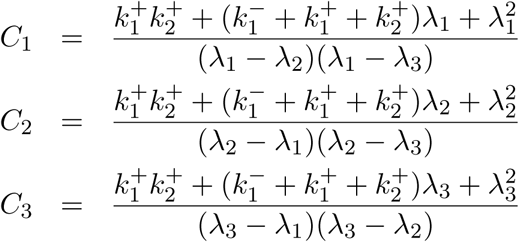

The unique solution of (29) and (31) is

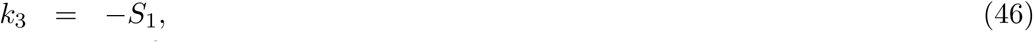

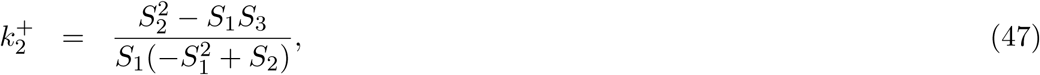

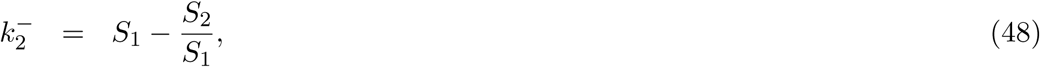

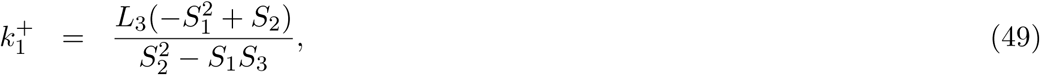

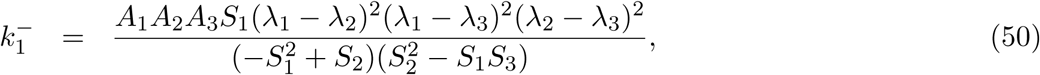

where

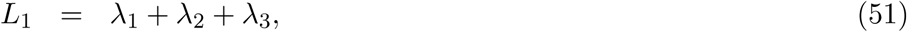

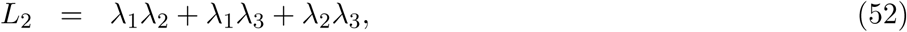

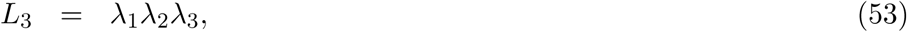

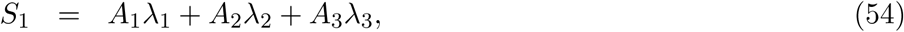

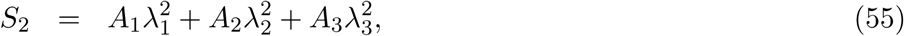

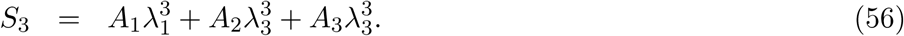

#### 4.5 Symbolic solution to the inverse problem for the M2 model (*N* = 3)

This model is described by the transitions 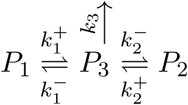. In this case *P*_3_ is the *ON* state. Model M2 is a type II model. It has a matrix of kinetic rates

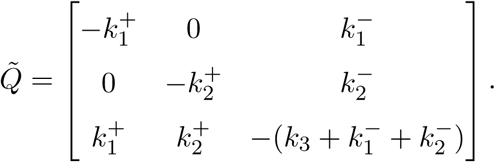

The characteristic polynomial of 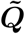 is 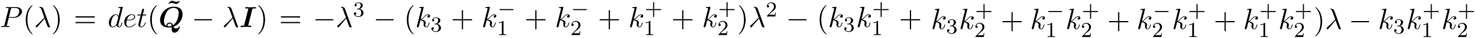.

The Vieta formulas read

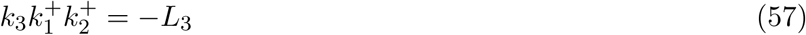

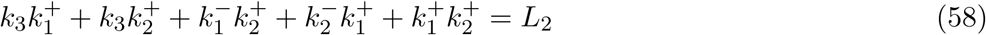

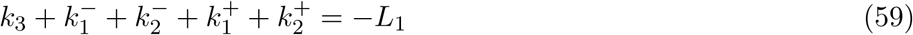

The solution of the recursion (36),(37),(38),(39) reads

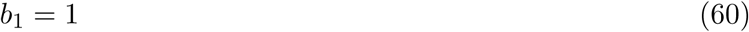

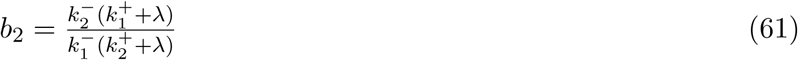

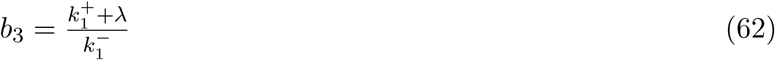

The system (30) has the solution

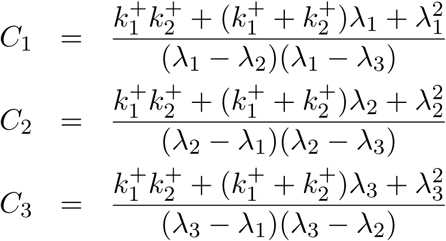

Up to the permution symmetry *P*_1_ ↔ *P*_2_ the solution of (29) and (31) is unique and described by

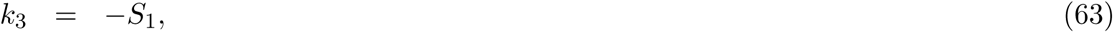

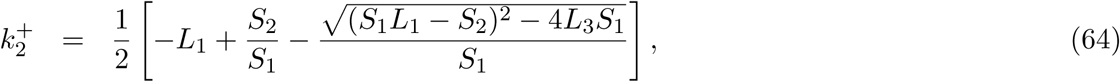

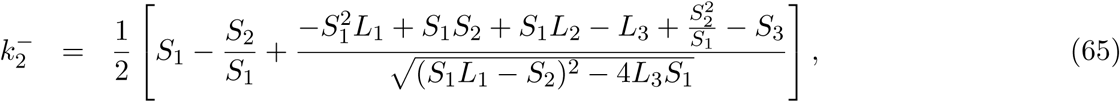

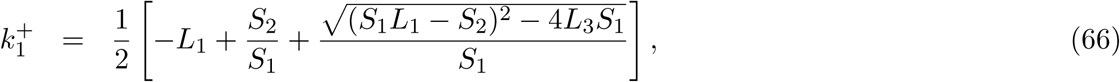

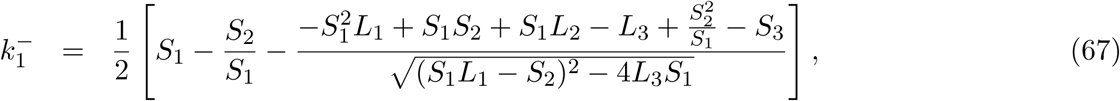

#### 4.6 Symbolic solution to the inverse problem for the M3 model (*N* = 3)

The chain without *P*_4_ is the same as the model M1. Like M1, M3 is a type I model. The difference is the position of the ON state which is in the middle of the chain (*P*_2_ is the ON state).

The matrix 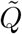 and its characteristic polynomial are the same as in the section 4.4. In particular, the Vieta relations remain the same:

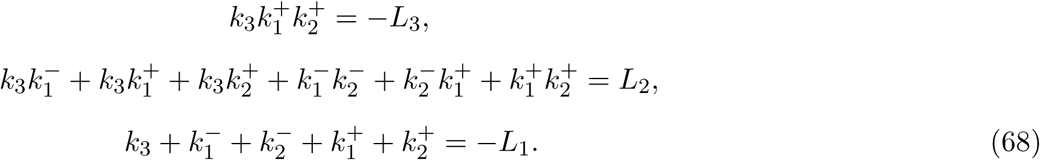

However, instead of computing the waiting time for reaching *P*_4_ starting from *P*_3_, we compute the waiting time for reaching *P*_4_ starting from *P*_2_. In this model, the significance of the states *P*_2_ and *P*_3_ is ON and PAUSE, respectively. The observed waiting time is from ON to EL, therefore from *P*_2_ to *P*_4_.

We look for solutions of the master equation (13) with initial conditions ***X***(0) = (0, 1, 0, 0).

Like in section in order to compute solutions of (13) we need the eigenvectors of 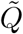. For the new initial conditions it is convenient to impose the normalization condition *u*_2_ = 1, where *u*_*i*_, 1 ≤ *i* ≤ 3 are the components of the eigenvector ***u***. We get

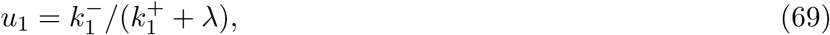

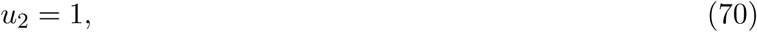

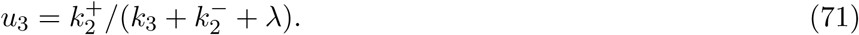

A solution of the (13) reads ***X***(*t*) = *C*_1_***u***_1_ exp(*λ*_1_*t*)+ *C*_2_***u***_2_ exp(*λ*_2_*t*)+ *C*_3_***u***_3_ exp(*λ*_3_*t*). From the initial conditions, it follows

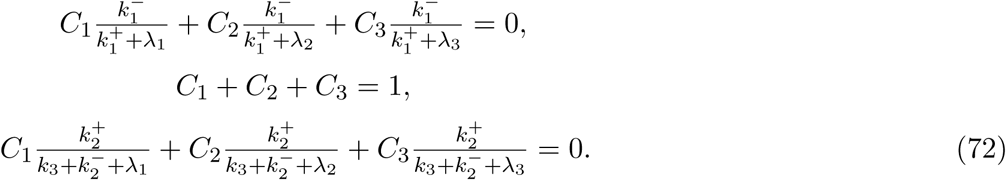

The system (72) has the solution

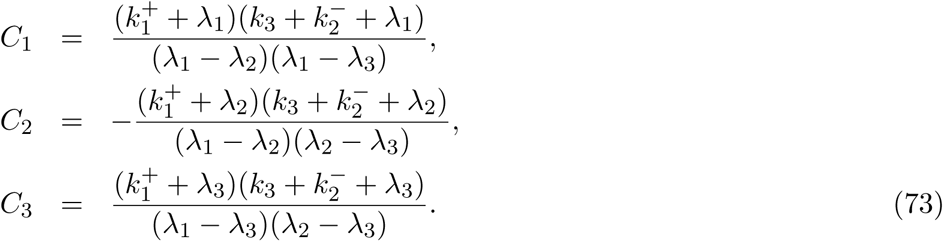

*X*_4_ obeys 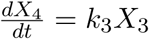 and the survival function is

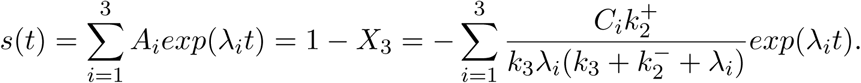

The relation between *C*_*i*_ and *A*_*i*_ reads:

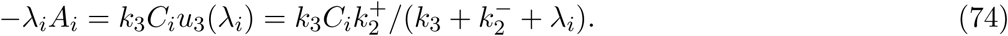

By definition *s*(0) = 1, therefore

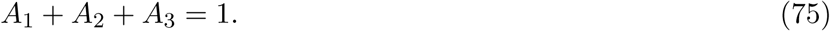

Using (74) and (29) we can show that

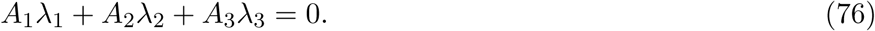

Eq.76 is very important. It implies that in this case, instead of 5 independent parameters, the survival function has only 4 independent parameters *A*_1_, *λ*_1_, *λ*_2_, *λ*_3_. Using (76) and (75) we can compute the remaining parameters as

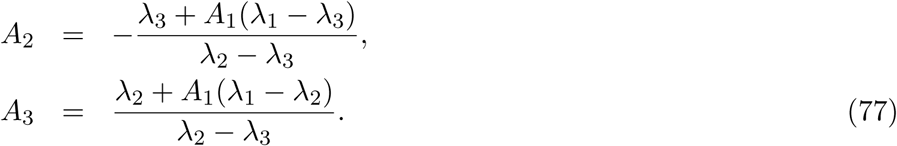

If the condition (76) is not satisfied, then the system formed from eqs. (68) and (74) (with *i* = 1, 2) is incompatible.

If the condition (76) is satisfied, then the system (68), (74) is indeterminate and has an infinity of solutions. In this case, all the solutions can be expressed as functions of a free parameter. In the sequel we will choose *k*_3_ as free parameter. This choice leads to the following symmetric expressions:

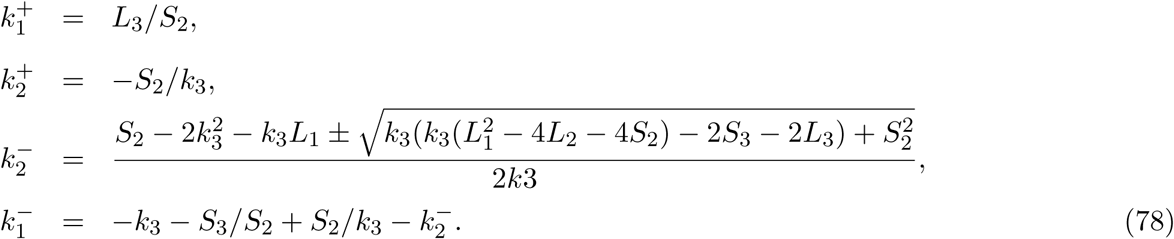

#### 4.7 Symbolic solution to the inverse problem for the two state ON-OFF model (*N* = 2)

The two states ON-OFF model (telegraph model) reads 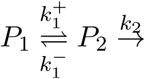.

In order to identify this model we use a two exponential fit of the survival function *S*(*t*) = *A*_1_ exp(*λ*_1_*t*) + *A*_2_ exp(*λ*_2_*t*). Without restricting the generality, we can consider that *λ*_1_ < *λ*_2_ < 0. Then, from *S*′(*t*) ≤ 0 it 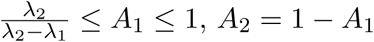.

From the parameters of the survival function we can compute the model parameters as follows

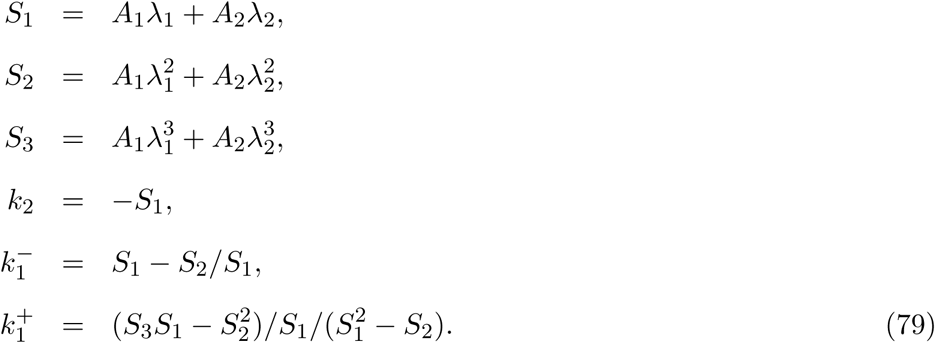

#### 4.8 Symbolic solution to the inverse problem for the four state chain model (*N* = 4)

This model is described by the transitions

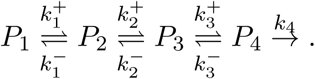

In this case *P*_4_ is the *ON* state. The model is of type I.

We have

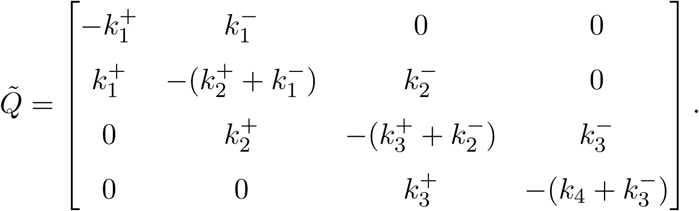

The Vieta formulas read

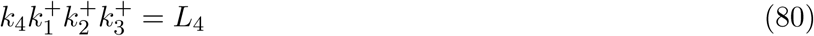

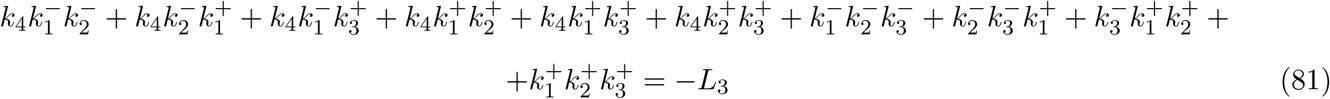

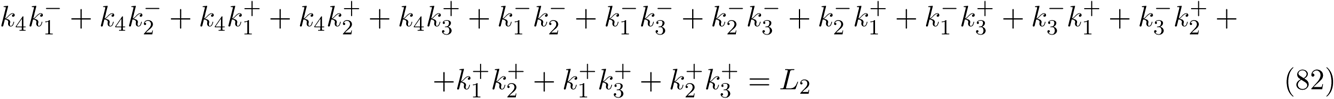

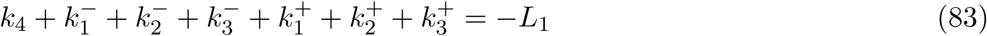

The solution of the recursion (34) is

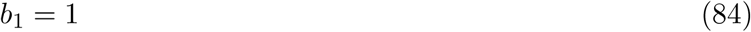

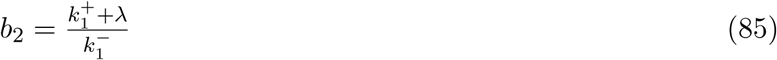

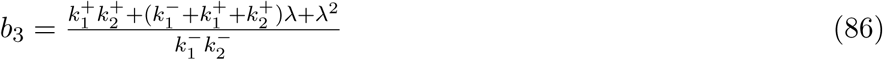

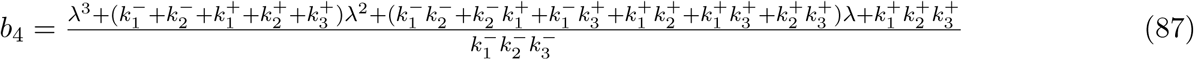

The system (30) has the solution

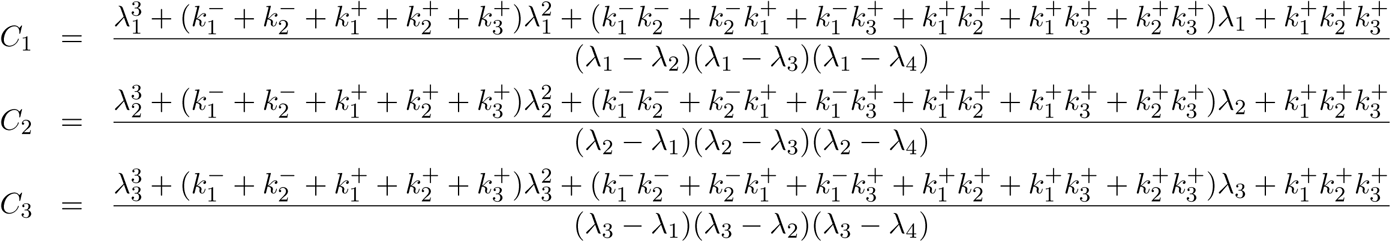

The eqs. (29) and (31) provide

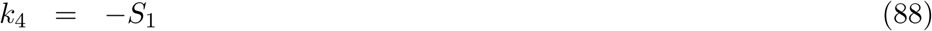

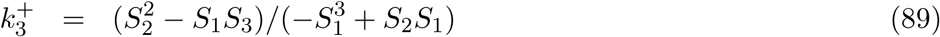

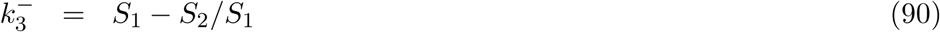

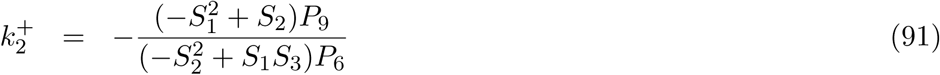

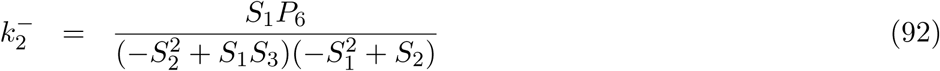

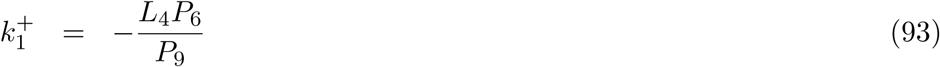

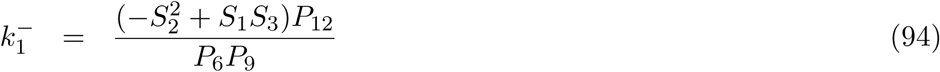

where 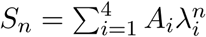,

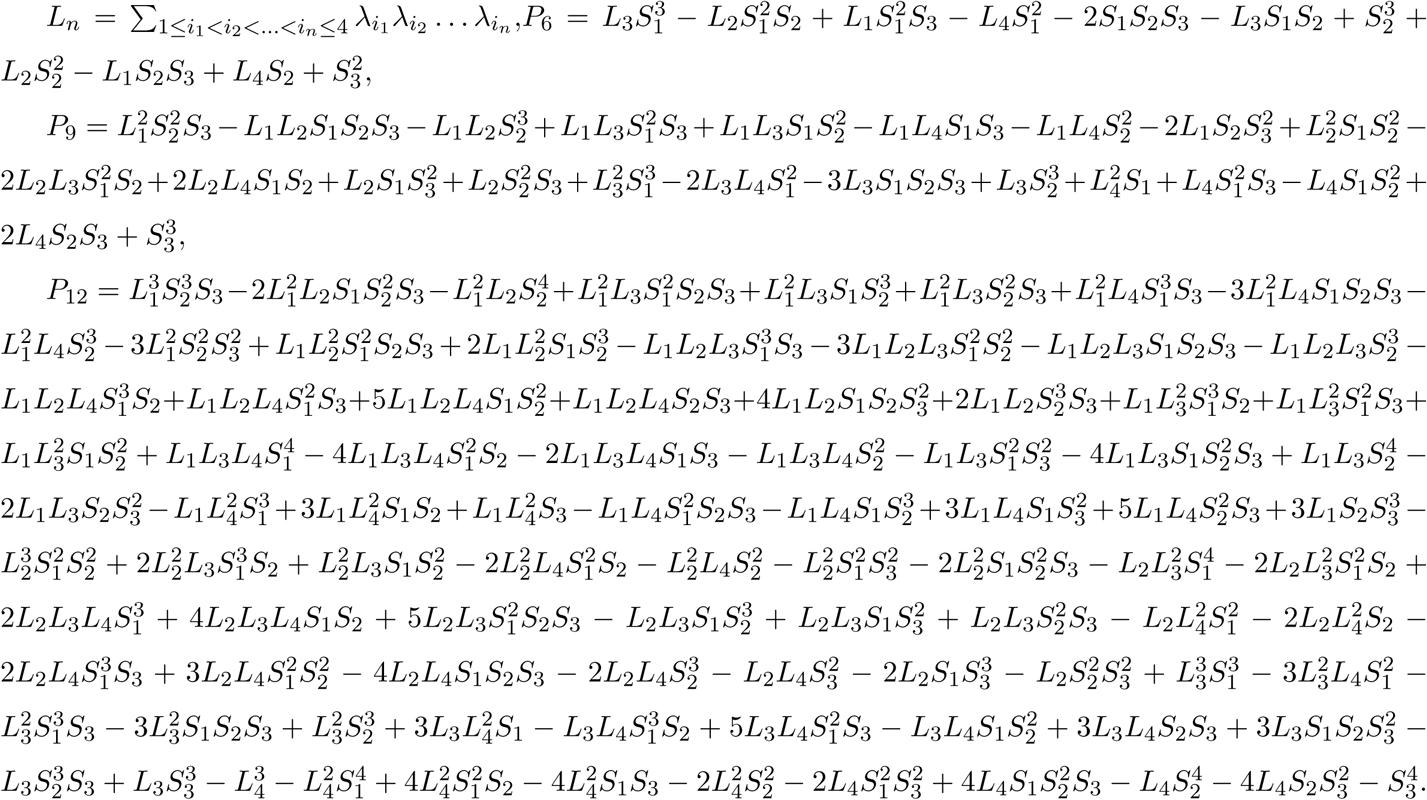

Using the relation *A*_1_ + *A*_2_ + *A*_3_ + *A*_4_ = 1, the above expressions can be simplified to

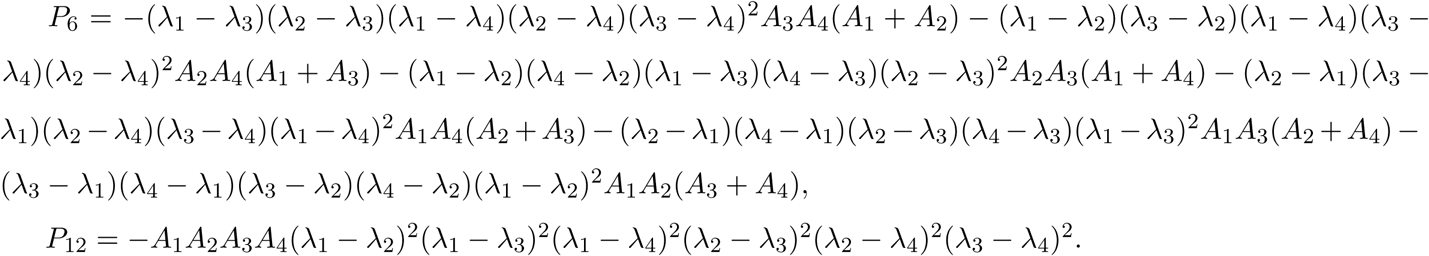

#### 4.9 Symbolic solution to the inverse problem for the four state model with branching (*N* = 4)

This model is described by the transitions 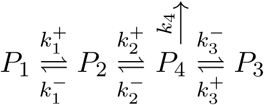. In this case *P*_4_ is the *ON* state. The model is of type II.

We have

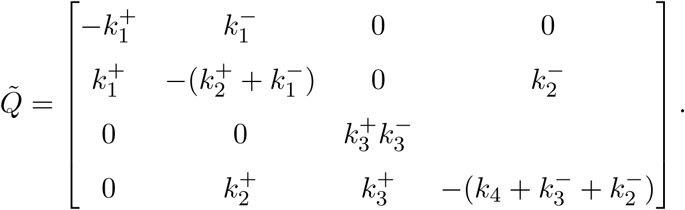

The Vieta formulas read

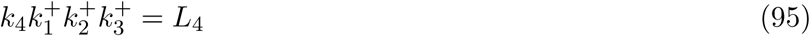

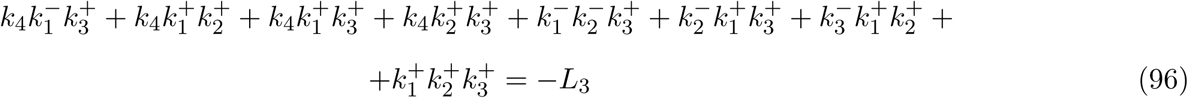

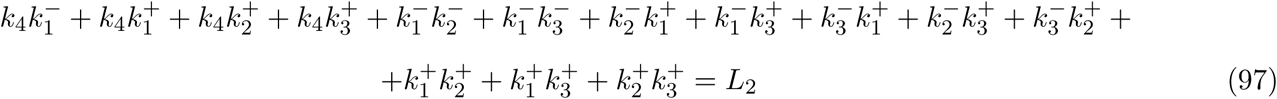

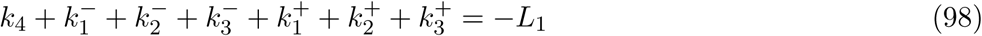

The eigenvectors of 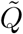 are

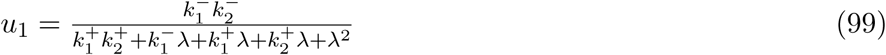

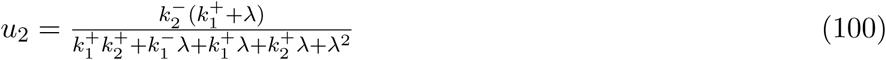

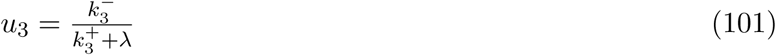

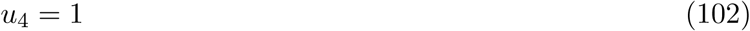

The system (30) has the solution

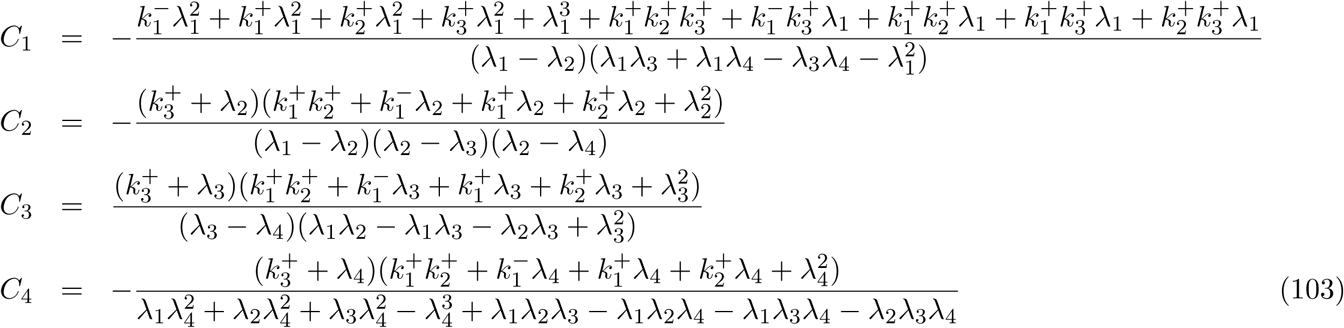

The eqs. (29) and (31) provide

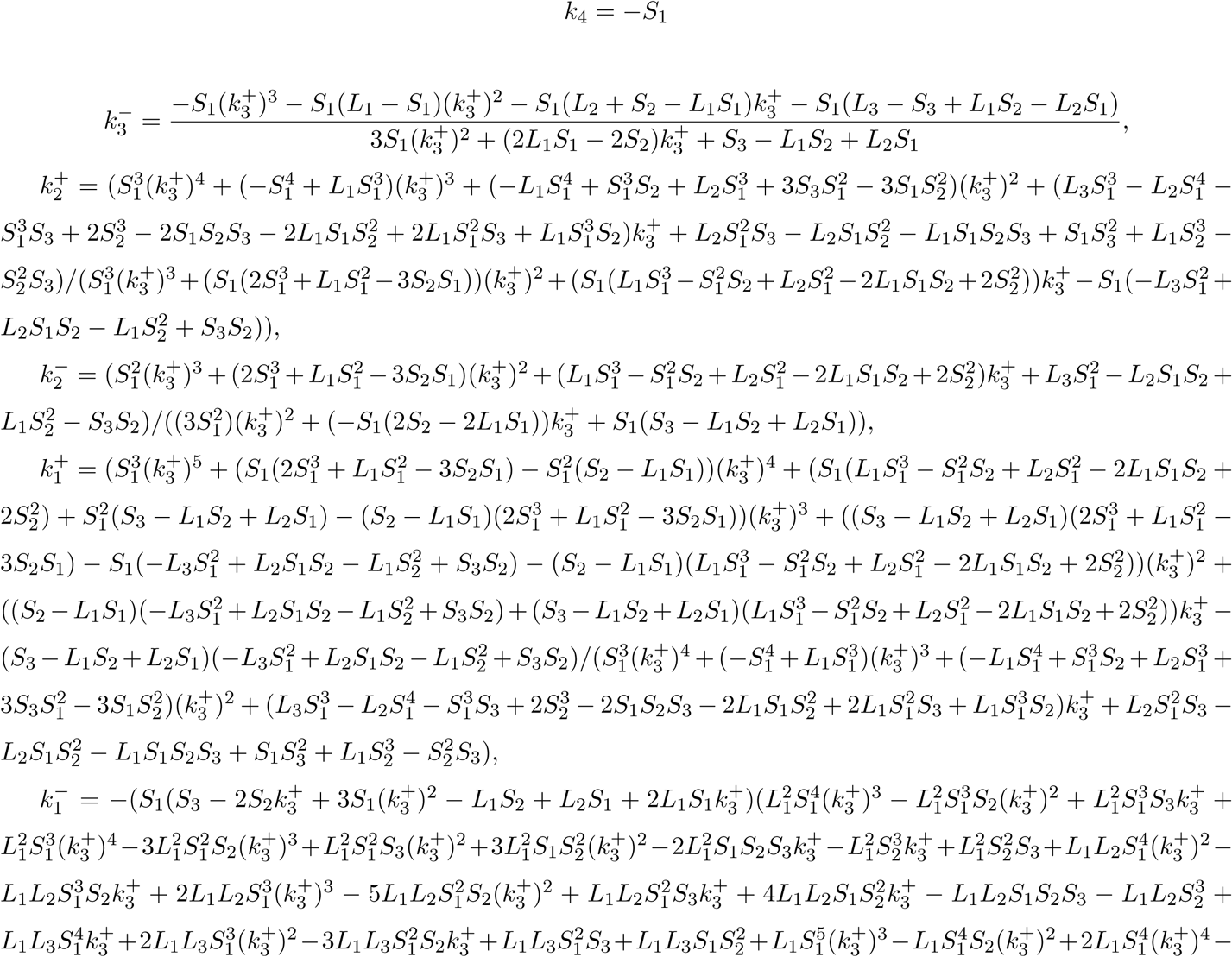

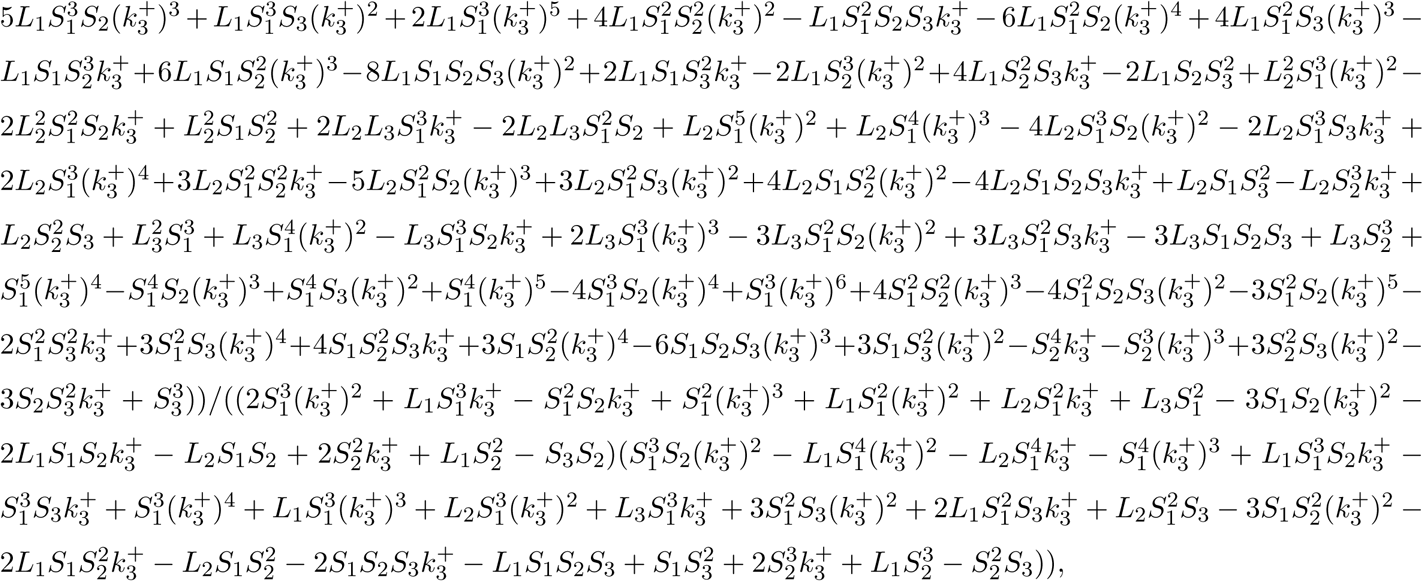

where 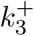 is the solution of the cubic equation

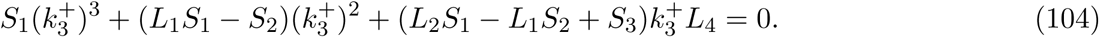

The equation (104) has the discriminant

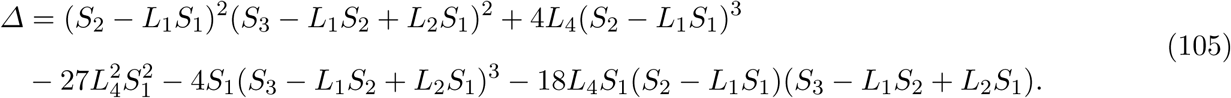

When *Δ* < 0, there is an unique real solution

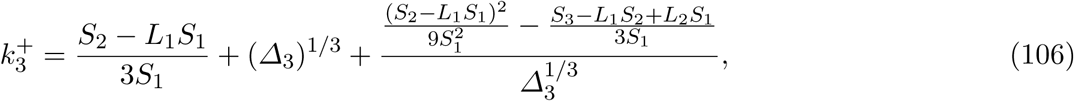

where 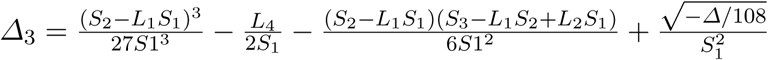.

#### 4.10 Symbolic solution to the inverse problem for four state chain with return in state *P*_3_ (*N* = 4)

This model has exactly the same transitions as the 4 state chain model described in the Section 4.8 with the difference that the ON state is *P*_3_.

Using the same methods as in Section 4.6 we show that in this case

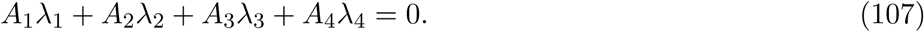

Using (107) and *A*_1_ + *A*_2_ + *A*_3_ + *A*_4_ = 1 we can compute the remaining parameters as

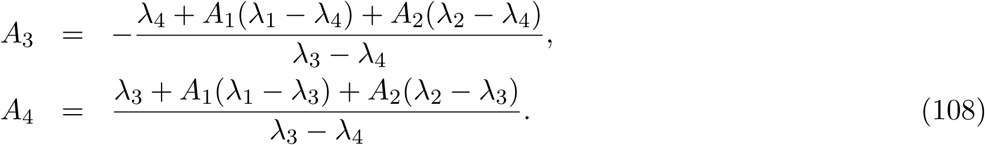

If the condition (107) is not satisfied, then there is no solution to the inverse problem.

If the condition (107) is satisfied, then the inverse problem is not well posed and has an infinity of solutions. In this case, all the solutions can be expressed as functions of a free parameter. In the sequel we will choose *k*_4_ as free parameter. Although we were able to obtain analytic solutions, these are too long to be displayed.

The following, simple relations are useful for the anaysis of this model:

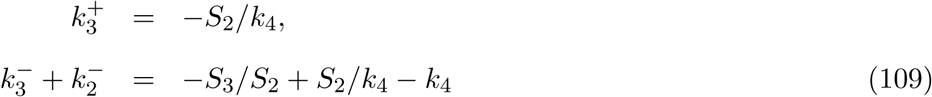

### 5 Uncertainty estimation for the model parameters

In Section 3.3 we have used optimization with multiple initial parameters to estimate confidence intervals for each parameter of the multi-exponential survival function as lower and upper bounds of optimal and sub-optimal parameters. These intervals are presented as 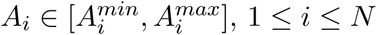 and 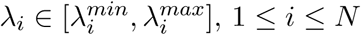.

In the sections above we have shown how to compute symbolically the kinetic parameters of various models from the parameters *A*_*i*_, *λ*_*i*_, 1 ≤ *i* ≤ *N* of the multi-exponential survival function. By applying the symbolic mapping to the confidence intervals 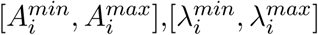 one can get the confidence intervals of the kinetic parameters. However, finding intervals that bound the kinetic parameters from the confidence intervals of the survival function parameters is a non-convex optimization problem with constraints which may prove difficult. Therefore, in the current implementation of our software we decided to apply the symbolic mapping directly to the entire set of optimal and sub-optimal survival function parameters obtained in Section 3.3 and compute the lower and upper bounds of the resulting kinetic parameters.

### 6 Computing the mean mRNA at the steady state

The statistics of the waiting time between two successive transcription initiations can be used to compute the statistics of the number of mRNA molecules. Each elongating polymerase will generate one molecule of mRNA that will survive in the average a time *T* ≈ 45 min. Therefore the mean mRNA number at the steady state is simply:

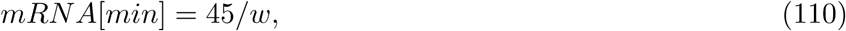

where *w* is the average waiting time.

The straightforward calculation

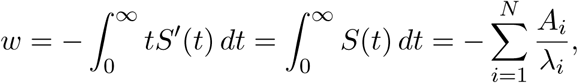

leads to two equivalent ways to compute the mean mRNA number, from the area under curve, or from the parameters of the survival function

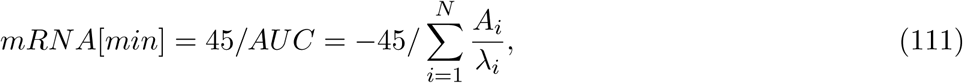

where AUC is the area under curve of the survival function.

### 7 Computing the probability of each state at stationarity

Although the computation of the distribution of waiting times does not require stationarity conditions (successive waiting times form a renewal process even without stationarity, as soon and as long as the model parameters are constant in time) it is usefull to have estimates for the stationary probabilities of being in each of the model’s state. The sojourn time in the state *P*_*N*+1_ being nil the probability of being in this state is also nil. The remaining *N* probabilities *p*_*i*_ = ℙ[*M* (*t*) = *P*_*i*_], 1 ≤ *i* ≤ *N* satisfy *p*_1_ + *p*_2_ + … + *p*_*N*_ = 1 and the following homogeneous system of linear equations:

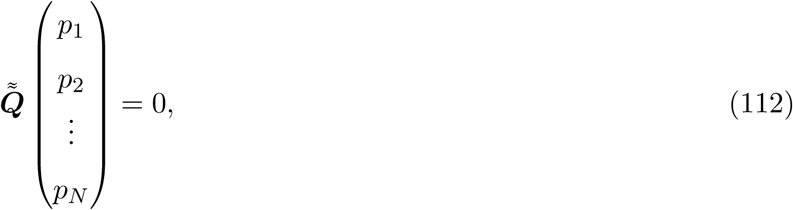

where 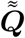 is obtained from 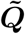 by setting *k*_*N,N*+1_ = 0.

A few examples follow.

For the model M1,

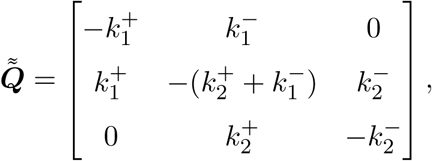

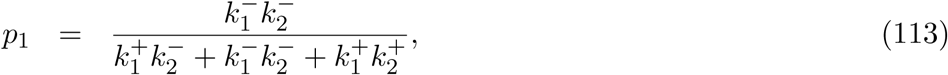

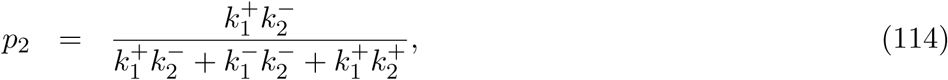

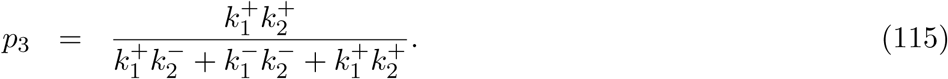

For the model M2,

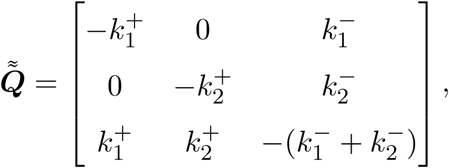

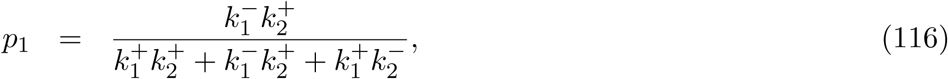

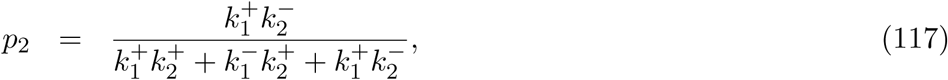

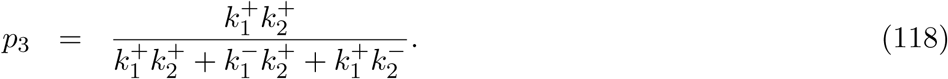

For the four states model,

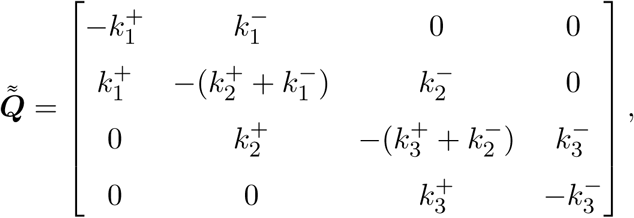

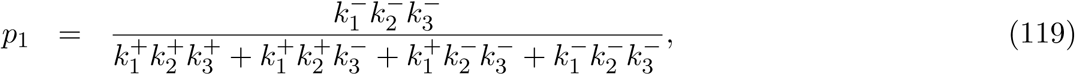

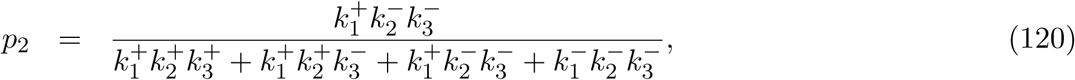

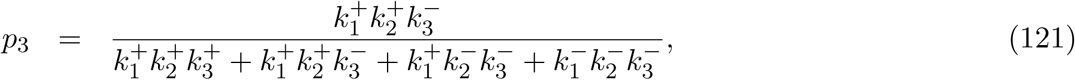

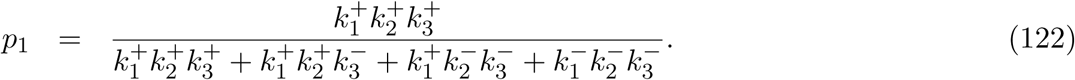

For the two states (ON-OFF) model

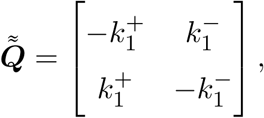

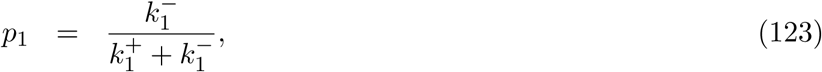

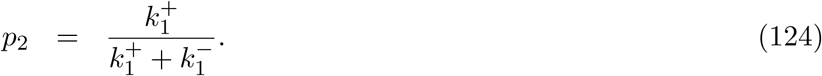

### 8 Testing the robustness of the method using artificial data

The numerical method is based on the assumption that the instrumental noise and other sources of noise are averaged out by the algorithm and therefore can be neglected. In this subsection we use artificial data to test the consequences of releasing this assumption. Furthermore, the optimization algorithm is stochastic and include approximate steps such as the estimation of the parameters *p*_*s*_ and *p*_*l*_, and errors resulting from the analog to digital conversion of the long movie signals. Artificially generated data with well know parameters will also allow us to test the fidelity of the parameter identification in our method.

Artificial data was generated by simulating the model M1 using the Gillespie algorithm. We use three parameter sets, similar to those identified from data in the three experimental conditions (previous subsection). The simulations generate artificial polymerase positions from which we first compute a noiseless signal using Eq. (1).

In a second step we add to the signal a centered Gaussian noise, whose variance is similar to the one in data, as follows

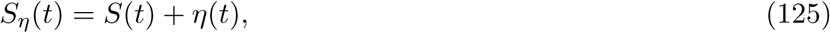

where *S*(*t*) is the noiseless signal and *η*(*t*) is the noise.

The noise estimate is obtained from the short movies data. It is defined as the difference between the raw signal and the signal reconstructed by deconvolution (computed using Eq. (1)). We found that the noise variance is an increasing function of the signal amplitude. By using cubic polynomial interpolation we have derived analytic formulas for the variance in the three experimental conditions:

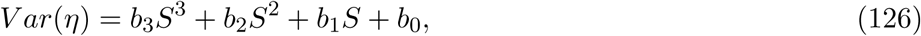

where *b*_*i*_, 0 ≤ *i* ≤ 3 are parameters whose values can be found in the Table 1.

**Table 1:** Noise parameters for various experimental conditions in the study of the HIV-1 promoter.

| Data set | $b_0$ | $b_1$ | $b_2$ | $b_3$ |
| --- | --- | --- | --- | --- |
| Low tat | 0.27 | 0.026 | 0.0022 | -1.5e-5 |
| No tat | -0.97 | 0.23 | -0.0021 | 9.8e-6 |
| High tat | 0.27 | 0.026 | 0.0022 | -1.5e-5 |

We applied our algorithm to a raw signal described by (125) and obtained estimates of the kinetic parameters. *η* is defined by (126) and Table 1. Together with *η* we have also tested the double 2*η* and four times 4*η* noise amplitude. These estimates were compared to the know values of the parameters that were used for simulating the artificial data. The result of the comparison is shown in Fig.13.

The method faithfully retrieves the parameter values, at least for noise amplitudes comparable to the ones determined from the data used in this study. For larger noise amplitudes some parameters may not be faithfully retrieved. As expected, some large kinetic parameters, corresponding to small time scales are not faithfully retrieved. However, the small parameters, corresponding to large time scales are faithfully retrieved even for large noise amplitudes. This proves the robustness of the method with respect to noise.

**Figure 13:**
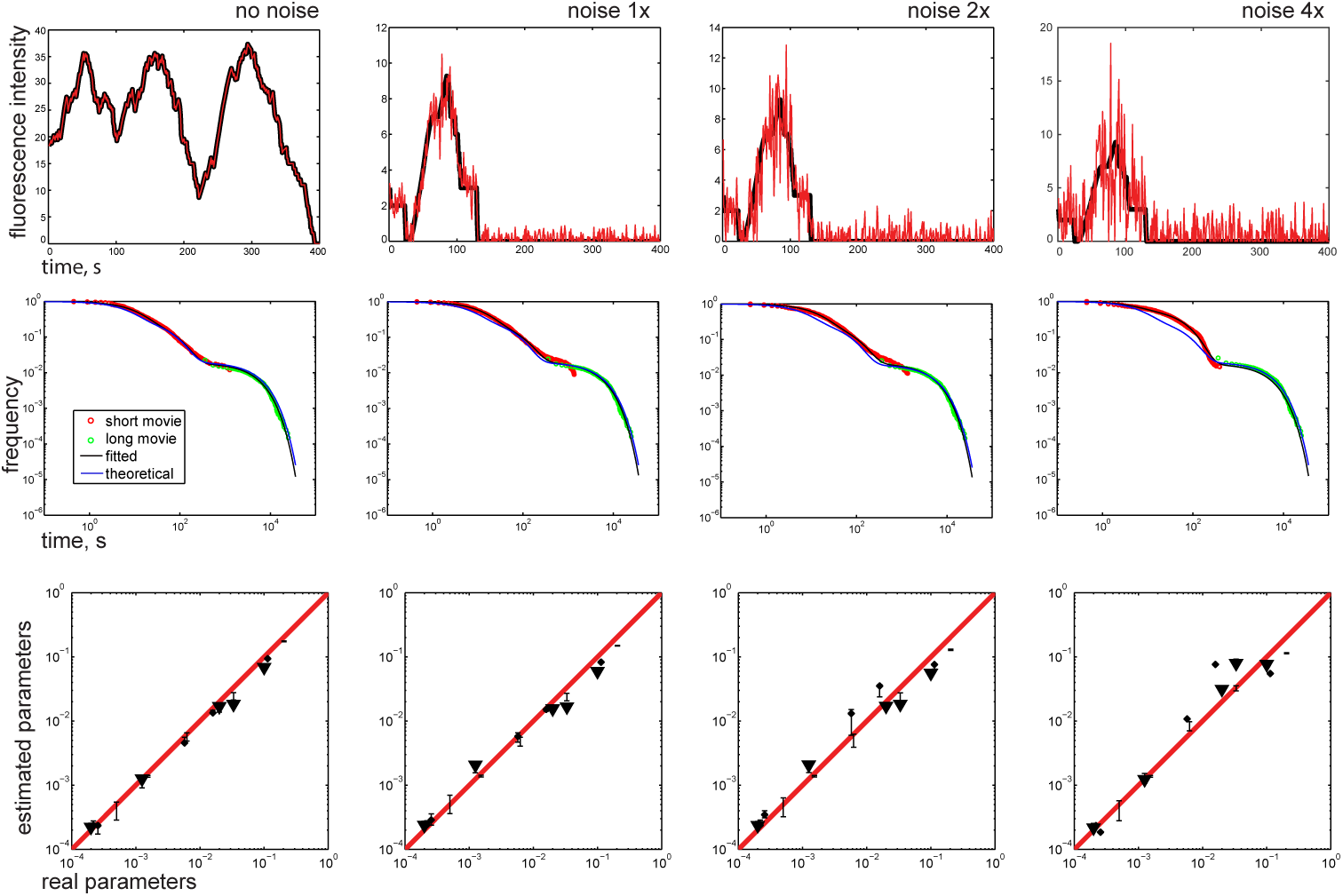
Testing the algorithm with artificial data for various noise amplitude. ×1 represents artificial data with the same amplitude of noise as the real data. ×0 is the noiseless artificial data. For each comparison we consider 3 sets of 5 parameters corresponding to the three experimental conditions in the HIV-1 data: no tat, low tat and high tat. The survival functions (middle row) and the artificial signal (upper row) are shown only for the no-tat conditions.

### 9 Results

#### 9.1 Identifying the parameters of the model M2

Model M2 corresponds to the stochastic, facultative pausing (Figure 1). The model parameters can be identified from a unconstrained three-exponential fit (Figure 12).

The results of the fit are presented in the Table 2 and in the Figure 14.

**Figure 14:**
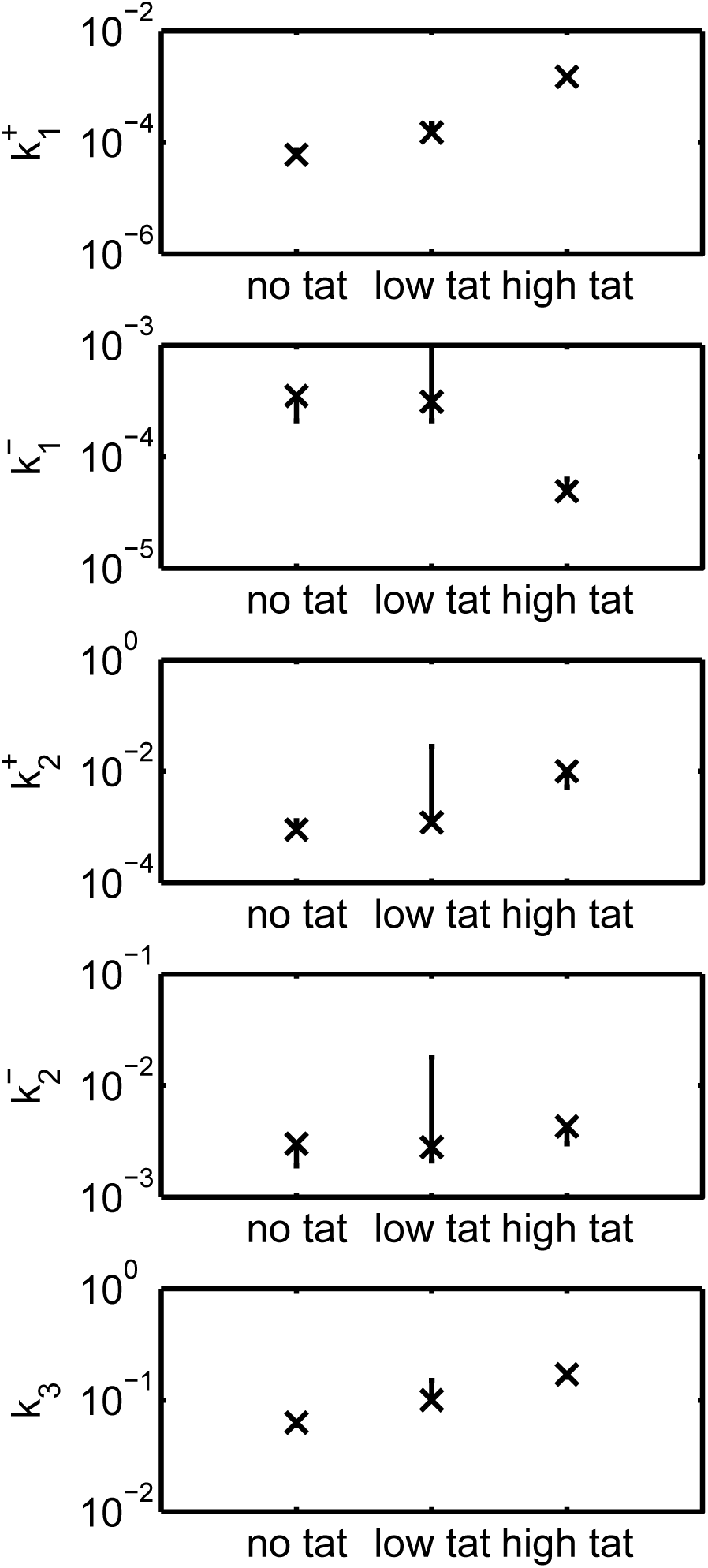
Results of the unconstrained three-exponential fit of the model M2. Parameter dependence on the experimental conditions for *α* = 0.30. The vertical bars are uncertainty intervals.

**Table 2:**
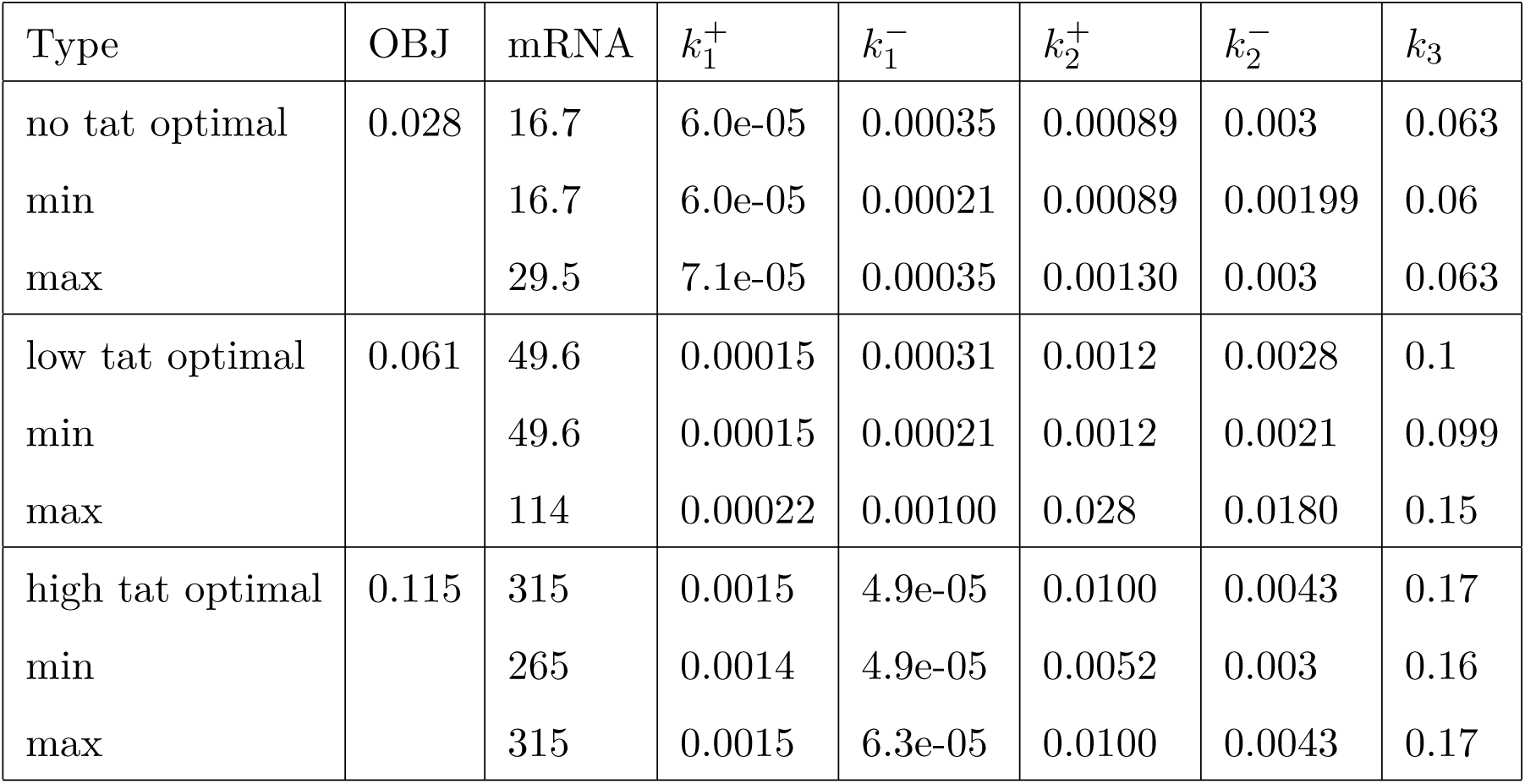
Results of the unconstrained three-exponential fit of the model M2. *α* = 0.30

#### 9.2 Identifying the parameters of the two states ON-OFF model

In order to identify this model we use a two exponential fit of the survival function *S*(*t*) = *A*_1_ exp(*λ*_1_*t*) + *A*_2_ exp(*λ*_2_*t*). The model parameters are computed from the survival function parameters according to the Section 4.7.

The result of the fit is given in the Table 3 and in the Figure 15. The large values of the objective function suggest that this model is not suitable for our data.

**Table 3:** Results of the two-exponential fit, *α* = 0.30. The objective function has large values compared to the three state model *M*_2_, for the same value of *α*.

| Type | OBJ | $\lambda_1$ | $\lambda_2$ | $A_1$ | $A_2$ | $k_2$ | $k_1^-$ | $k_1^+$ | mRNA |
| --- | --- | --- | --- | --- | --- | --- | --- | --- | --- |
| no tat optimal | 0.22046 | -0.0478 | -0.000141 | 0.984 | 0.0159 | 0.047 | 0.000755 | 0.000144 | 20.3 |
| min |  | -0.0478 | -0.000141 | 0.984 | 0.00646 | 0.0444 | 0.000287 | 0.000136 |  |
| max |  | -0.0446 | -0.000135 | 0.994 | 0.0159 | 0.047 | 0.000755 | 0.000144 |  |
| low tat optimal | 0.273 | -0.0782 | -0.00025 | 0.991 | 0.00943 | 0.0774 | 0.000733 | 0.000252 | 53.5 |
| min |  | -0.0782 | -0.00025 | 0.991 | 0.00368 | 0.0719 | 0.000264 | 0.000244 |  |
| max |  | -0.0721 | -0.000243 | 0.996 | 0.00943 | 0.0774 | 0.000733 | 0.000252 |  |
| high tat optimal | 0.64799 | -0.12 | -0.00222 | 0.996 | 0.00422 | 0.12 | 0.000488 | 0.00223 | 264.8 |
| min |  | -0.12 | -0.00222 | 0.00278 | 0.00113 | 0.097 | 0.000105 | 0.00218 |  |
| max |  | -0.0971 | -0.00218 | 0.999 | 0.997 | 0.12 | 0.000488 | 0.00223 |  |

**Figure 15:**
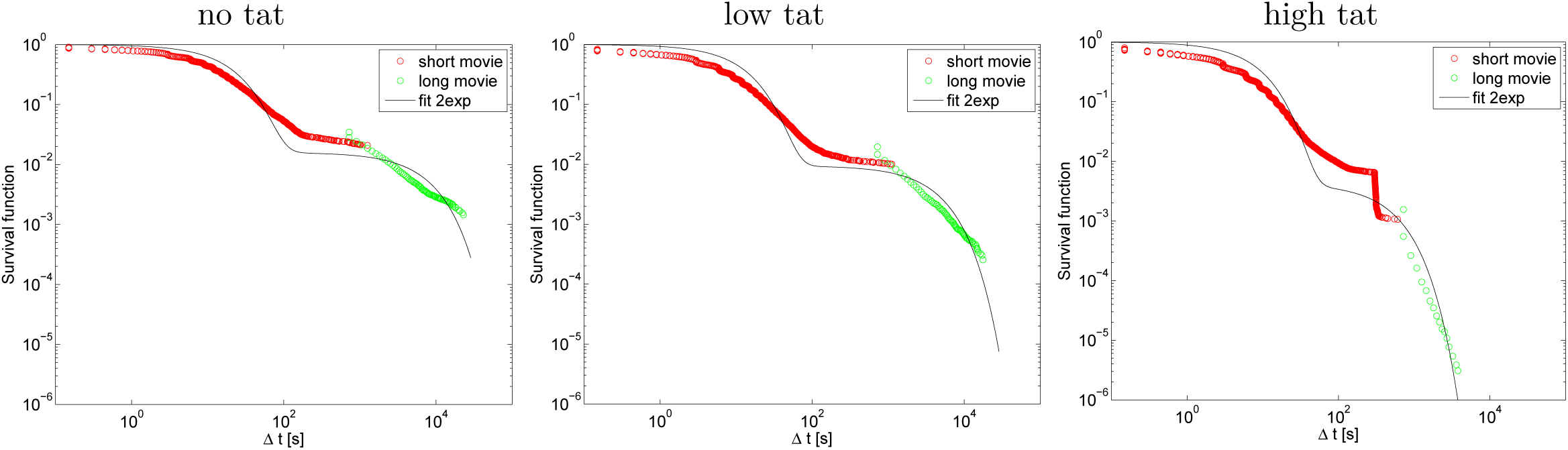
Results of the two-exponential fit: most optimal fit for *α* = 0.30.

#### 9.3 Identifying the parameters of the model M3

Model M3 corresponds to obligatory pausing (see Figure 1) and is identified using the constrained three-exponential fit described in the Section 4.6. However, even if (up to errors) the three-exponential fit provides a single best fit, the set of corresponding parameters of model M3 is a curve in the 5D space of parameters. The inverse problem for model M3 is not well posed as the relation between the parameters of the model M3 and the parameters of the three-exponential fit is many to one. The result of the constrained three-exponential fit is given in the Table 4.

**Table 4:**
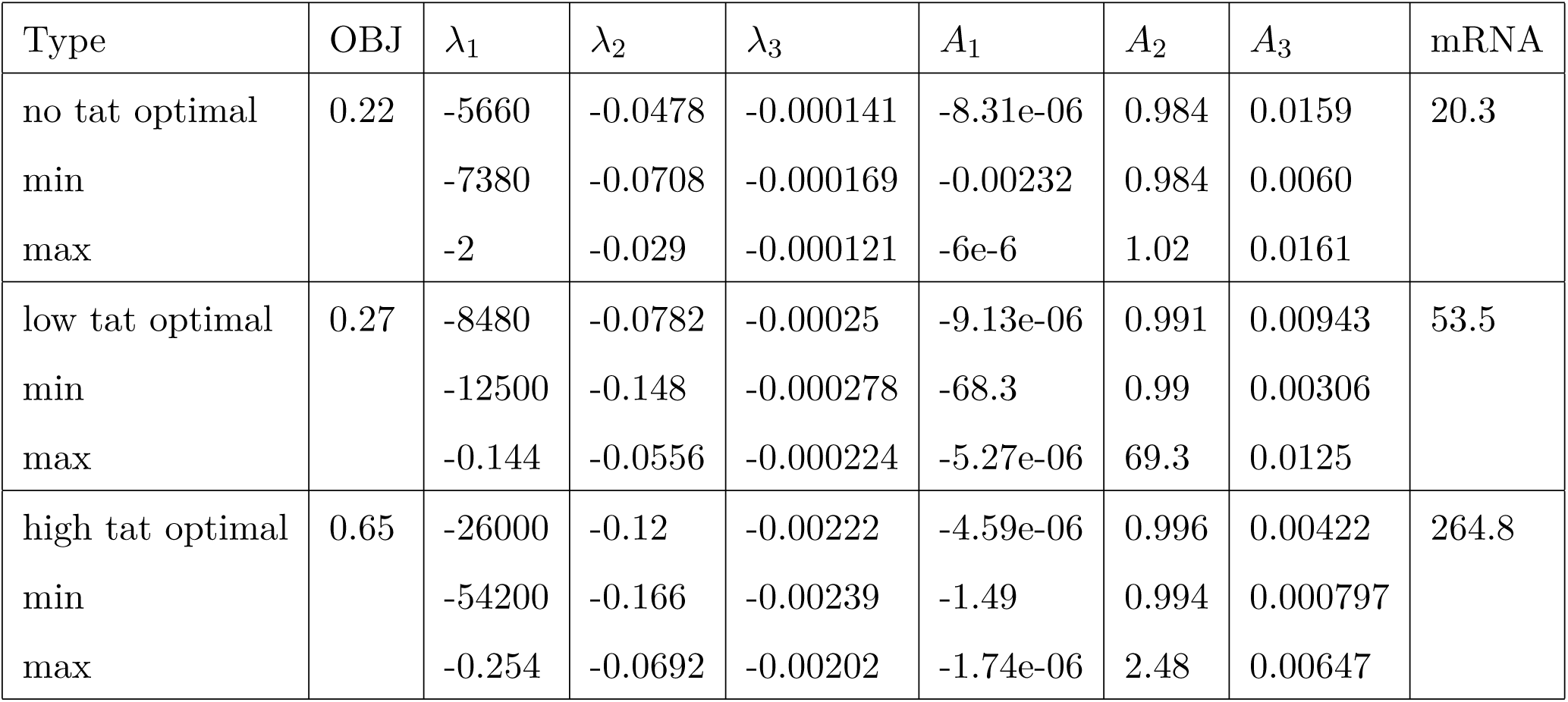
Results of the constrained three-exponential fit of the model M3, *α* = 0.30. The objective function has large values (compared to different models and for the same *α*) and the fitted parameters are very uncertain.

| Type | OBJ | $\lambda_1$ | $\lambda_2$ | $\lambda_3$ | $A_1$ | $A_2$ | $A_3$ | mRNA |
| --- | --- | --- | --- | --- | --- | --- | --- | --- |
| no tat optimal | 0.22 | -5660 | -0.0478 | -0.000141 | -8.31e-06 | 0.984 | 0.0159 | 20.3 |
| min |  | -7380 | -0.0708 | -0.000169 | -0.00232 | 0.984 | 0.0060 |  |
| max |  | -2 | -0.029 | -0.000121 | -6e-6 | 1.02 | 0.0161 |  |
| low tat optimal | 0.27 | -8480 | -0.0782 | -0.00025 | -9.13e-06 | 0.991 | 0.00943 | 53.5 |
| min |  | -12500 | -0.148 | -0.000278 | -68.3 | 0.99 | 0.00306 |  |
| max |  | -0.144 | -0.0556 | -0.000224 | -5.27e-06 | 69.3 | 0.0125 |  |
| high tat optimal | 0.65 | -26000 | -0.12 | -0.00222 | -4.59e-06 | 0.996 | 0.00422 | 264.8 |
| min |  | -54200 | -0.166 | -0.00239 | -1.49 | 0.994 | 0.000797 |  |
| max |  | -0.254 | -0.0692 | -0.00202 | -1.74e-06 | 2.48 | 0.00647 |  |

**Table 5:** Results of the constrained four-exponential fit of the model M4, *α* = 0.3. The objective function shows that the fit is not better than the one of the model M2, for the same *α* and the fitted parameters are very uncertain.

| Type | OBJ | $\lambda_1$ | $\lambda_2$ | $\lambda_3$ | $\lambda_4$ | $A_1$ | $A_2$ | $A_3$ | $A_4$ |
| --- | --- | --- | --- | --- | --- | --- | --- | --- | --- |
| no tat optimal | 0.036875 | -8430 | -0.0636 | -0.00103 | -6.6e-05 | -7.23e-06 | 0.958 | 0.0373 | 0.0043 |
| min |  | -19700 | -0.087 | -0.0121 | -0.00011 | -0.0698 | 0.869 | 0.0261 | 0.00306 |
| max |  | -1.02 | -0.0597 | -0.00103 | -6.6e-05 | -3.06e-06 | 1.02 | 0.126 | 0.00477 |
| low tat optimal | 0.078907 | -5550 | -0.102 | -0.0018 | -0.000161 | -1.79e-05 | 0.976 | 0.0221 | 0.00226 |
| min |  | -36600 | -1.74 | -0.0487 | -0.000228 | -335 | 0.716 | 0.0198 | 0.00212 |
| max |  | -0.202 | -0.0975 | -0.00157 | -0.000156 | -2.97e-06 | 336 | 0.453 | 0.00525 |
| high tat optimal | 0.11601 | -9330 | -0.174 | -0.0101 | -0.00148 | -1.82e-05 | 0.972 | 0.0281 | 0.000288 |
| min |  | -85200 | -0.178 | -0.0126 | -0.00149 | -0.0442 | 0.968 | 0.0218 | 0.000258 |
| max |  | -4.1 | -0.166 | -0.00681 | -0.00144 | -1.91e-06 | 1.01 | 0.0323 | 0.000349 |

The dependence of the parameters of the model M3 on the undetermined parameter *k*_3_ is shown in the Figure 17. The parameters 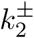 have very large values compared to all other parameters. The model *M* 3 is in this case equivalent to the two states ON-OFF model and inherits the difficulty of this model to fit the data.

**Figure 16:**
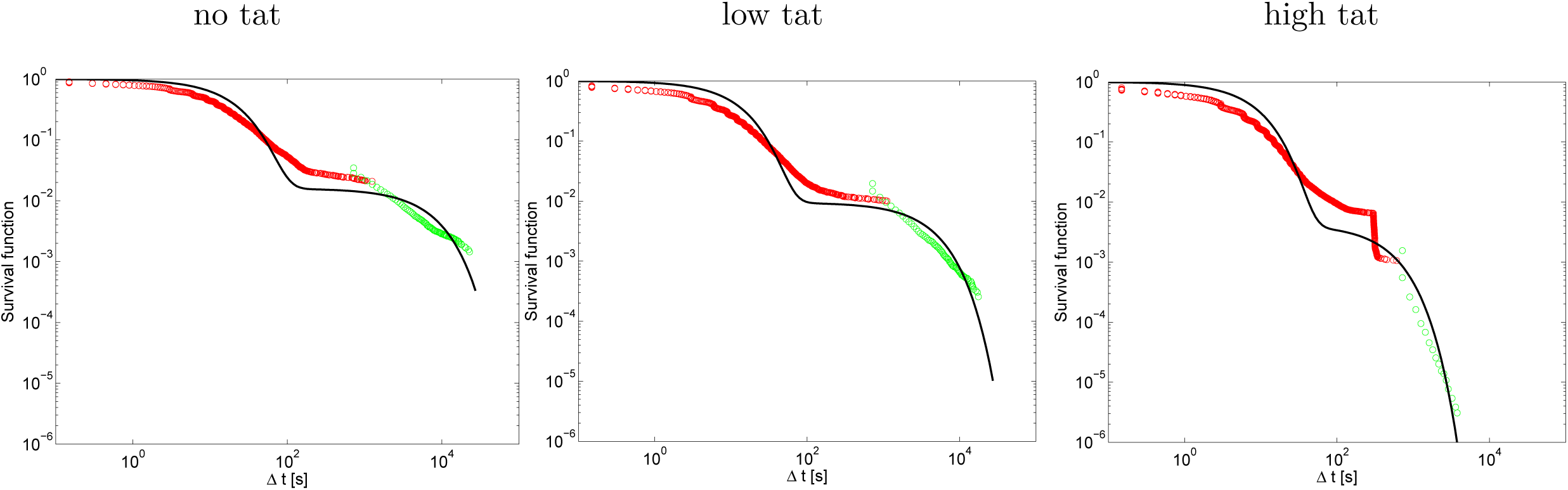
Results of the constrained three-exponential fit: most optimal fit for *α* = 0.30.

**Figure 17:**
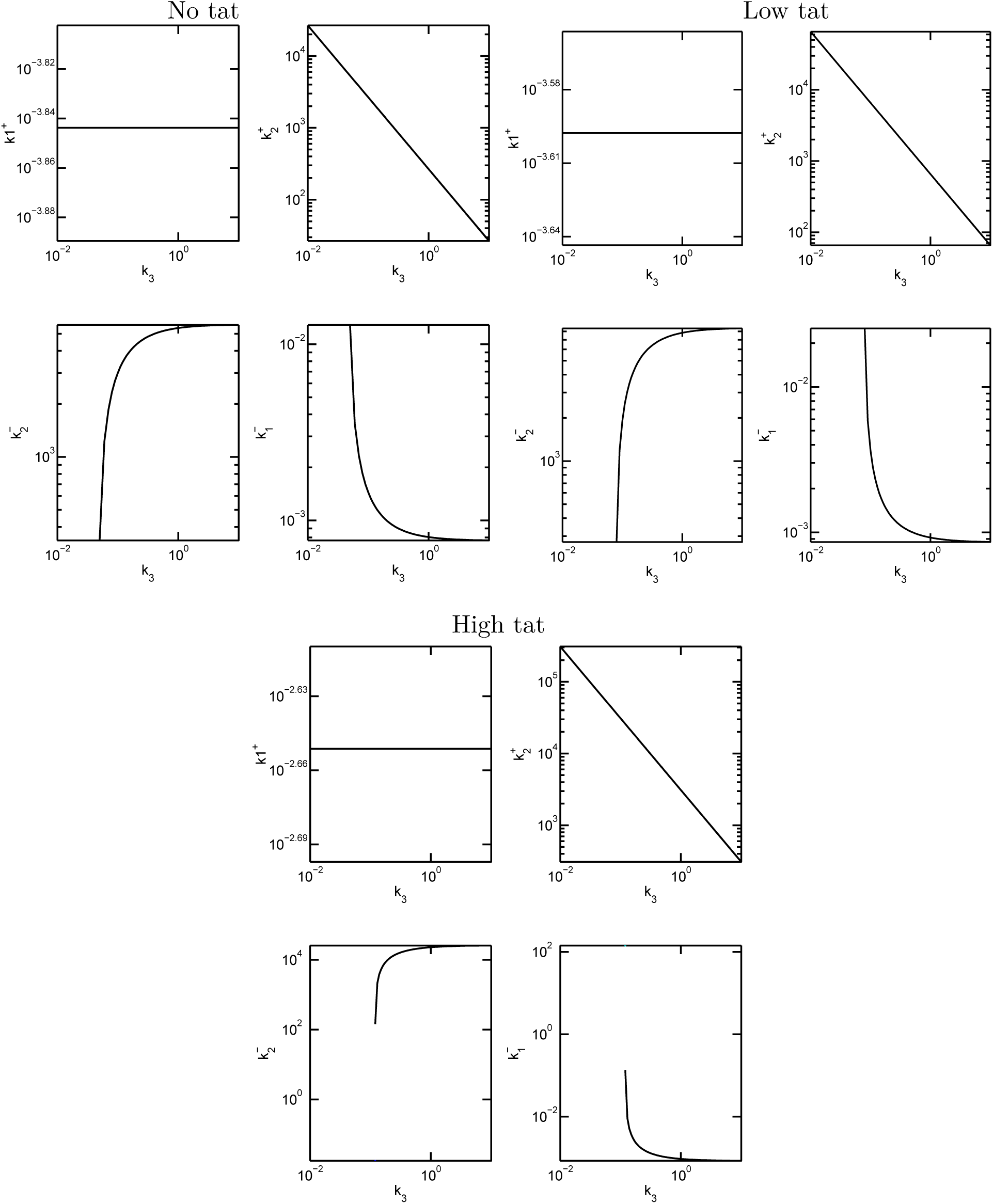
Results of the constrained three-exponential fit of the model M3. Parameter dependence on the undetermined parameter *k*_3_ (pause exit rate) for *α* = 0.30. The parameters 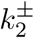 have very large values compared to all other parameters and correspond to very fast processes (timescales smaller than 0.01*s*). For such parameters the model *M* 3 is equivalent to a two states ON-OFF model (the states ON and PAUSE can be pooled with no information loss in the model M3). In order to ensure positivity of kinetic parameters, one needs *k*_3_ > 0.1*s*^−1^.

#### 9.4 Identifying the parameters of a four states model with pausing

In order to identify four states models we use a four exponential fit of the survival function *S*(*t*) = *A*_1_ exp(*λ*_1_*t*)+ *A*_2_ exp(*λ*_2_*t*)+*A*_3_ exp(*λ*_3_*t*)+*A*_4_ exp(*λ*_4_*t*), where *A*_1_+*A*_2_+*A*_3_+*A*_4_ = 1. Let us consider that *λ*_1_ < *λ*_2_ < *λ*_3_ < *λ*_4_ < 0. From *S*′(*t*) ≤ 0 it follows *λ*_1_*A*_1_ + *λ*_2_*A*_2_ + *λ*_3_*A*_3_ + *λ*_4_*A*_4_ ≤ 0, *A*_4_ ≥ 0.

The model *M*_4_ is obtained by adding one more OFF state to the model *M*_3_ (see Figure 18). It corresponds to the theoretical model described in the Section 4.10. The parameters of this model can be obtained from a constrained four exponential fit with six free parameters *λ*_1_,*λ*_2_,*λ*_3_,*λ*_4_,*A*_1_,*A*_2_ (see Eq.(108)). Although this model has more free parameters than the model *M*_2_, the fit quality is lower.

**Figure 18:**
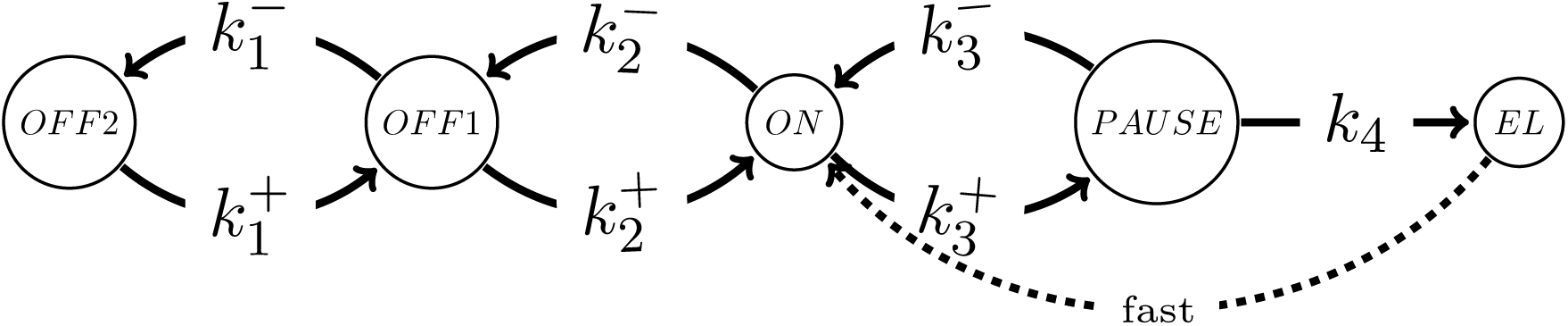
Model M4 with two OFF states and obligatory pausing. *k*_4_ is the pause exit rate, 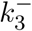 is the transcription abortion rate.

**Figure 19:**
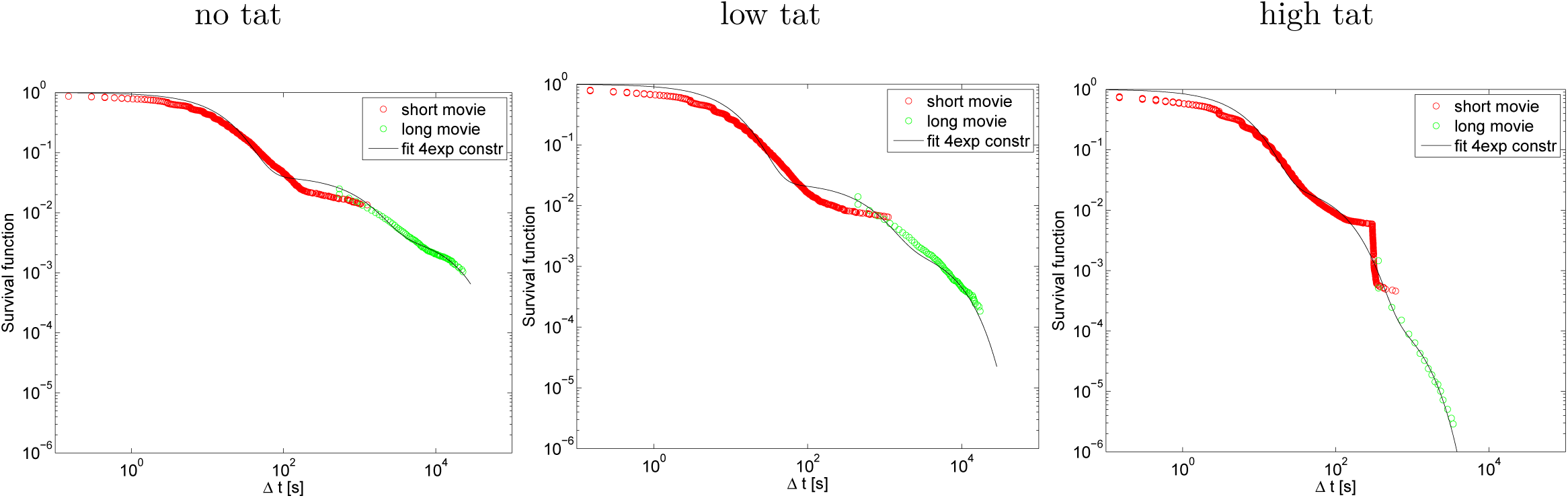
Results of the constrained four-exponential fit of the model M4: most optimal fit for *α* = 0.30.

**Figure 20:**
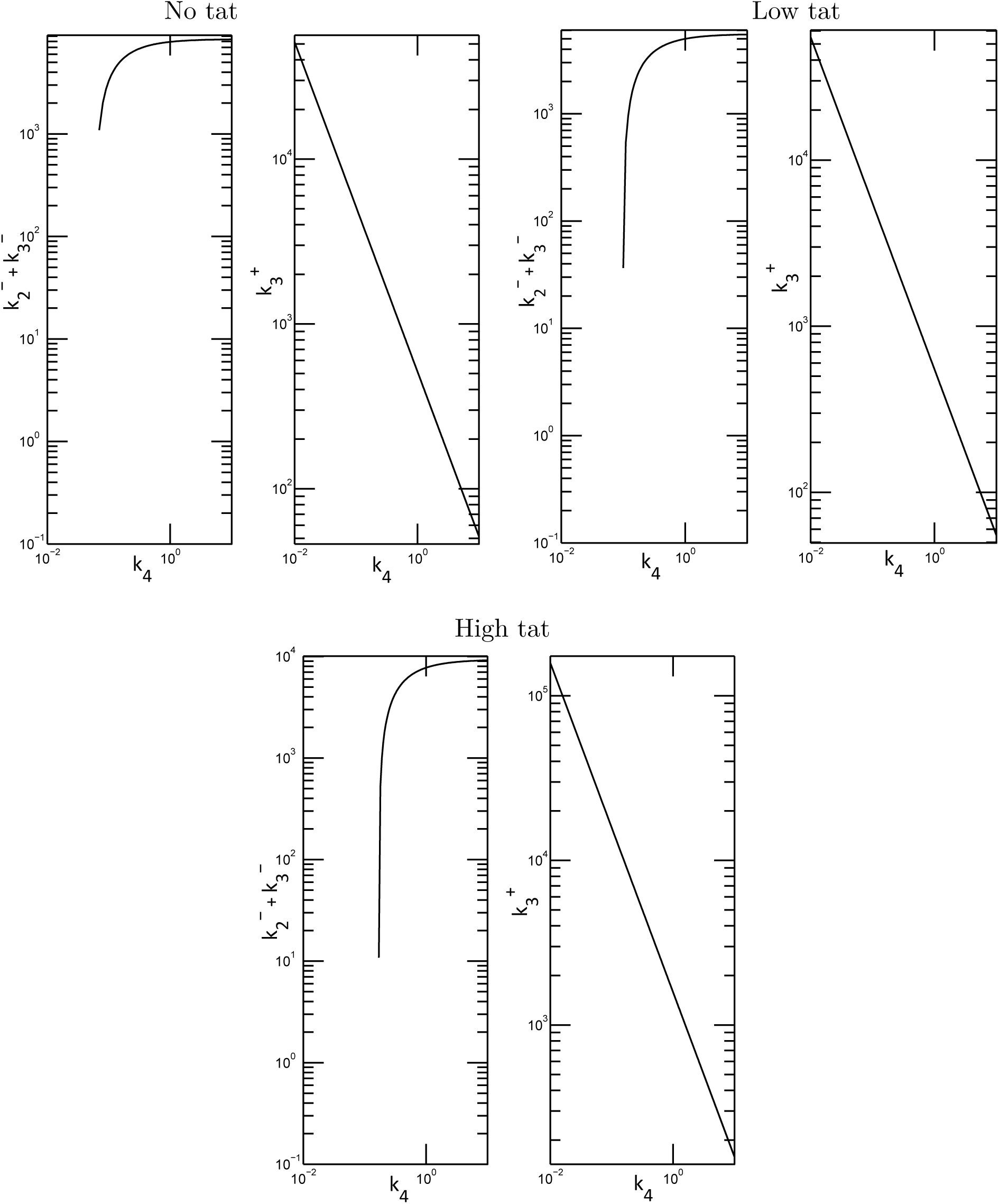
Results of the constrained four-exponential fit of the model M4. Parameter dependence on the undetermined parameter *k*_4_ (pause exit rate) for *α* = 0.30. The parameters 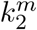 and 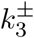 have very large values compared to other parameters and correspond to very rapid processes (timescales smaller than 0.1*s*). With such parameters the model M4 is equivalent to a three states model with the states ON and PAUSE pooled. For positivity of kinetic parameters one needs *k*_4_ > 0.1*s*^−1^.

